# MAAMOUL: Metabolic network-based discovery of microbiome-metabolome shifts in disease

**DOI:** 10.64898/2026.03.27.714614

**Authors:** Efrat Muller, Shiri Baum, Elhanan Borenstein

## Abstract

**Motivation:** A central goal in human gut microbiome research is to identify disease-associated functional shifts, an objective increasingly pursued through metagenomic and metabolomic assays. However, common differential abundance analyses of genes or metabolites often yield long and difficult-to-interpret feature lists. Aggregating features into predefined pathways can improve interpretability but relies on fixed pathway boundaries that may not reflect context-specific functional changes. Moreover, even when paired metagenomic-metabolomic data are available, they are often analyzed separately or linked only through simple statistical associations.

**Results:** We introduce MAAMOUL, a knowledge-based computational framework that integrates metagenomic and metabolomic data to identify disease-associated, data-driven microbial metabolic modules. Leveraging prior knowledge of bacterial metabolism, MAAMOUL maps disease-association scores onto a global microbiome-wide metabolic network and identifies custom modules enriched for altered genes and metabolites. Applying MAAMOUL to inflammatory bowel disease (IBD) and irritable bowel syndrome (IBS) datasets revealed significant disease-associated modules not detected by conventional pathway-level analysis. In IBD, modules reflected disrupted sulfur and aromatic amino acid metabolism and enhanced microbial nucleotide salvage, whereas in IBS they linked purine and nicotinate/nicotinamide metabolism. These results demonstrate that network-guided multi-omic integration can uncover coherent functional shifts in the gut microbiome overlooked by single-omic or purely statistical approaches.

**Availability:** MAAMOUL is available as an R package at https://github.com/borenstein-lab/MAAMOUL.

## 1 Introduction

A key task in many human gut metagenomic studies is to identify functional shifts in the gut community, i.e., increased or decreased capacity to perform specific (often metabolic) functions, that arise during disease states (Wilmanski *et al*. 2021). While taxonomic profiles are cost-effective to obtain, e.g., using 16S rRNA amplicon sequencing, functional analyses based on metagenomics or metabolomics often provide a more robust and nuanced characterization of gut microbiome health, independent of species composition (Li *et al*. 2014, Vieira-Silva *et al*. 2016).

Standard computational pipelines for metagenomic data typically quantify the abundance of gene families or enzyme commission (EC) numbers in each sample. Subsequently, differential abundance analysis is performed at the gene- or EC-level, often yielding long, difficult-to-interpret, lists of significant features. To enhance interpretability, researchers may aggregate features into predefined metabolic pathways, i.e., specific sets of genes/enzymes, which, collectively, are responsible for some metabolic activity with well-defined input and output metabolites (Khatri, Sirota, and Butte 2012). Such pathway definitions can be obtained through curated databases such as KEGG (Kanehisa *et al*. 2017), MetaCyc (Caspi *et al*. 2014), or human microbiome-specific resources (Pascal Andreu *et al*. 2023). Despite their utility, pathway-level analyses have significant drawbacks: they are commonly too coarse, potentially masking biologically meaningful variation due to irrelevant parts of the pathway, or failing to account for the interface between different pathways. Moreover, universal pathway definitions may fail to account for species-specific metabolism and are often biased toward model organisms (Darzi *et al*. 2016). Similar challenges arise in metabolomics, where metabolite grouping into predefined classes or pathways faces analogous interpretability and bias limitations (Ren *et al*. 2015).

The integration of gene- and metabolite-based microbiome profiles, in contrast to the independent analysis of each omic, may facilitate mechanistic interpretations that span molecular layers. Indeed, metagenomics and metabolomic assays of fecal samples are increasingly coupled, adopting a multi-omic approach and aiming to achieve a systems-level view of the gut microbial ecosystem (Turnbaugh and Gordon 2008, Muller, Algavi, and Borenstein 2022). Methods for integrating such multi-omic datasets, accounting both for the association of each omic with the disease of interest and for cross-omic relationships have been proposed (Singh *et al*. 2019, Muller, Shiryan, and Borenstein 2024, Newman *et al*. 2024), yet, they typically rely on statistical tests or machine learning and ignore prior knowledge about mechanistic links between features. As a result, integration results must be followed by tedious manual exploration to pinpoint biological hypotheses that are mechanistically feasible.

To address these challenges, we introduce MAAMOUL, a knowledge-based framework that integrates microbiome and metabolome data to identify custom microbial metabolic modules associated with disease. By projecting metagenomics and metabolomics data onto a comprehensive microbiome-wide metabolic network, MAAMOUL identifies custom subgraphs or ‘modules’ enriched for differentially abundant features, including both enzymes and metabolites. This approach provides a “middle ground” between single-feature and pathway-based analysis, capturing sets of mechanistically linked features that change cohesively in disease without relying on predefined pathway boundaries.

While this approach, often referred to as “active module” discovery, has been utilized in other domains (see for example (Nguyen *et al*. 2019)), existing microbiome-specific methods such as MetaPath (Liu and Pop 2011) and metaModules (May *et al*. 2016) do not explicitly support the integration of different omics or the complexities of networks containing both metabolite and enzyme nodes. Our ‘multi-omic’ approach thus offers a significant advantage, aiming to combine bacterial genomic potential with actual metabolic outputs, improving the confidence and interpretability of disease-associated functional shifts. Below, we describe the MAAMOUL method and demonstrate its utility across several case-control cohorts, identifying several significant disease-associated metabolic modules supported by both metagenomic and metabolomic data that are overlooked by traditional pathway-level analysis.

## 2 Methods

### 2.1. The MAAMOUL Algorithm

To improve the analysis of human-associated paired microbiome-metabolome data, and to better detect disease-associated functional shifts, we introduce **MAAMOUL** (**M**icrobiome **A**ssociation **A**nalysis of **M**ulti-**O**mic data using a **U**niversal metabolic mode**L**). MAAMOUL identifies *custom data-driven microbiome metabolic modules* that are significantly dysregulated in disease, integrating metagenomics and metabolomics data within the context of known microbial metabolism. The method takes three inputs: (a) p-values for metabolic reactions (ECs), representing their association with disease and derived from some statistical testing of a metagenomic dataset; (b) p-values for metabolites, similarly derived based on a paired metabolomic dataset; and (c) a global metabolic network represented as an undirected bipartite graph connecting reaction (EC) nodes to their substrate and product metabolite nodes (Figure 1A). The following steps are then performed:

**Fig. 1.**
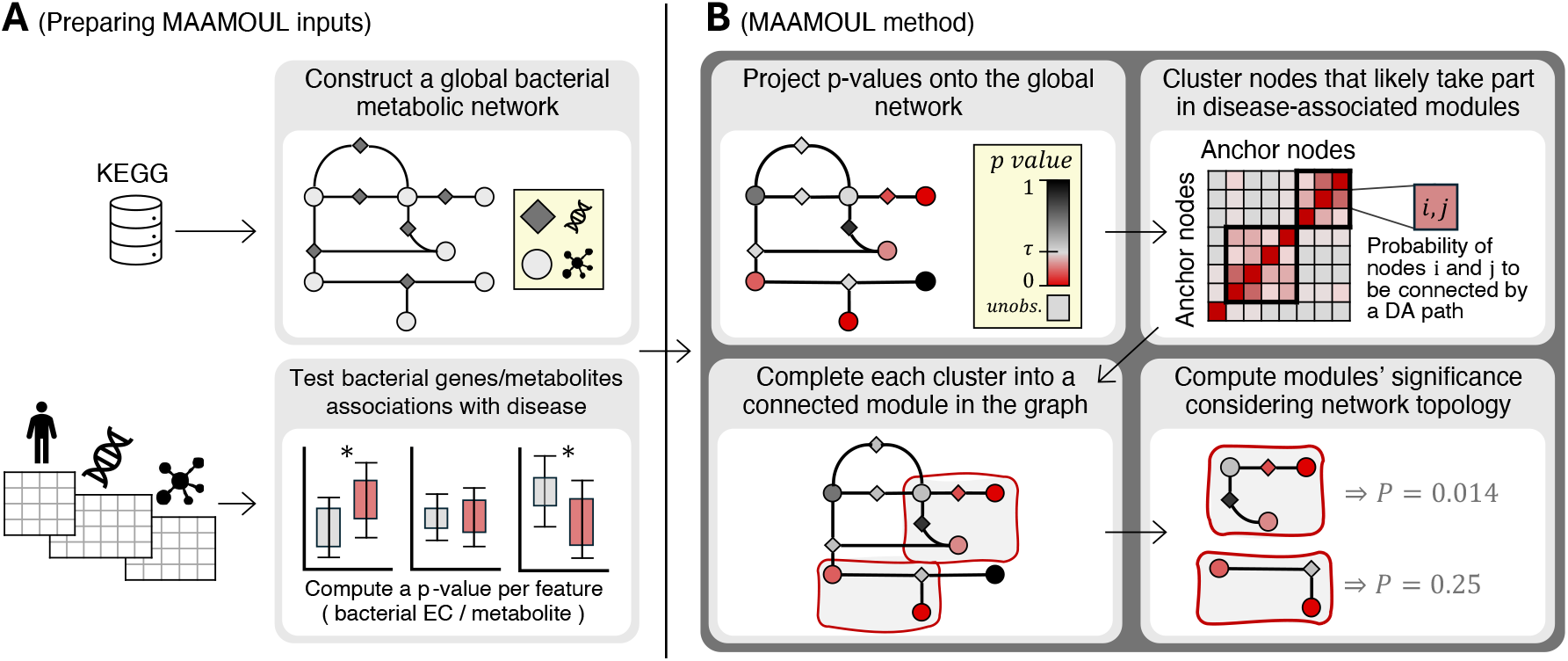
MAAMOUL method overview. **(A)** The MAAMOUL method receives as input a list of EC-metabolite edges, forming a global metabolic network (top), and a p-value for each feature observed in the data (EC or metabolite) reflecting its association with a disease/condition of interest (bottom). **(B)** Based on this input, MAAMOUL first projects p-values onto the global network, and then identifies clusters of proximate nodes that are likely to be involved in the same disease-associated module. Next, clusters are completed into connected subgraphs (“modules”) in the network based on the Steiner tree algorithm. Finally, a topology-aware permutation test is used to determine the statistical significance of each module.

#### Step 1: Projecting disease-association p-values onto the global metabolic network

First, each EC or metabolite node in the network with an available p-value (i.e., an “*observed*” node) is assigned its corresponding p-value (Figure 1B, top left). Nodes lacking a p-value (e.g., metabolites not assayed or ECs absent from the reference database), are referred to as “*unobserved*”.

#### Step 2: P-value modelling with a beta-uniform mixture model

Assuming p-values include a mixture of signal and noise, we modeled their distribution using the signal-noise decomposition approach of (Pounds and Morris 2003) and (Dittrich *et al*. 2008). Specifically, p-values are modeled as a mixture of a uniform distribution over [0,1] (noise; representing nodes not associated with the disease with p-values under the null hypothesis), and a Beta distribution *B*(*a*, 1) with a shape parameter *a* ∈ (0,1) (signal; representing disease-associated nodes), with a mixture parameter *λ* ∈ (0,1). The resulting beta-uniform mixture (BUM) distribution is *f*(*x*|*a, λ*) = *λ* + (1 − *λ*) ⋅ *a* ⋅ *x*^*a*-1^. Maximum likelihood estimates of *a* and *λ*, along with a p-value threshold τ corresponding to FDR=0.1, are then computed as previously described (Pounds and Morris 2003). This model is used below to impute p-values for the unobserved nodes.

#### Step 3: Identifying modules of disease-associated nodes

Given the global metabolic network with p-values assigned to observed nodes, MAAMOUL aims to detect small network regions (modules) that are enriched for disease-associated nodes. This task can be formulated in different ways, balancing factors such as module size, association strength, or the presence of unobserved nodes. MAAMOUL approaches this task with two assumptions: (i) Each EC or metabolite is either disease associated (DA) or not, with its P-value reflecting the probability of being DA, and (ii) DA nodes connected by short paths of other DA nodes are likely part of the same disease-associated module.

To estimate, for each pair of nodes, the likelihood that they are in the same DA module, MAAMOUL applies an iterated sampling approach. In each iteration, p-values are first assigned to unobserved nodes by sampling from the corresponding (ECs or metabolites) BUM model. Nodes are then randomly labeled as DA or non-DA, with the probability of being DA inversely-proportional to their p-value. Finally, MAAMOUL considers the subgraph induced by DA nodes only and uses a breadth-first search to determine for each pair of nodes whether they are connected by a path of length < *k*.

Across iterations, MAAMOUL records the fraction of times each node pair is connected, constructing a matrix *M* ∈ [0,1]^|*V*|×|*V*|^. To focus on high confidence modules, we define nodes that are both observed *and* have a p-value < τ as ‘anchors’, and filter *M* to include only these anchor nodes. The filtered matrix thus captures the likelihood that each pair of anchor nodes are connected by a short DA path and thus belong to the same disease-associated module. MAAMOUL then clusters these nodes using average-linkage hierarchical clustering and cuts the tree at a predefined threshold (0.8 in our analyses) to define modules (Figure 1B, top right).

#### Step 4: Completing modules using Steiner Trees

Since clusters of anchor nodes may not necessarily form a connected subgraph in *G* (which includes also unobserved or non-significant nodes), MAAMOUL next identifies a minimal subgraph of *G* connecting all anchors in each cluster (Figure 1B, bottom left), using a Steiner tree heuristic approach (Sadeghi and Fröhlich 2013): Briefly, each anchor node initially forms its own sub-tree; shortest paths between subtrees in *G* are computed and one of these paths is randomly selected, merging the corresponding subtrees. This process repeats until all anchors in a certain cluster are connected. The final module is defined as the subgraph induced by all nodes identified through this procedure (not necessarily a tree).

#### Step 5: Assessing modules’ significance

Certain topological properties of the global metabolic network (e.g., dense regions) may increase the chance of detecting modules enriched with low p-values purely due to the network’s structure rather than real biological signal. MAAMOUL therefore estimates the statistical significance of each module (Figure 1B, bottom right), accounting for the network’s topology. Specifically, for a module with *N*_*EC*_ anchor EC nodes, *N*_*met*_ anchor metabolite nodes, and average p-value 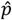, the nodes’ p-values are repeatedly shuffled (within each node type separately) and the module identification procedure is rerun. The module p-value is then computed as the fraction of shuffled networks resulting in a module with ≥ *N*_*EC*_ anchor EC’s, ≥ *N*_*met*_ anchor metabolites, and ≤ 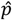 average p-value. To account for cases where multiple valid modules are detected in the real network, shuffled networks must also result in at least as many valid modules. Module p-values are FDR-corrected, and modules with FDR< 0.1 are considered significant. The final output is thus a list of significant modules: connected subgraphs in the global metabolic network likely associated with disease.

### 2.2. Implementation and Application

MAAMOUL is implemented as an open-source R package. We evaluated its performance using several publicly available microbiome-metabolome datasets, focusing on cohorts of patients with Crohn’s disease (CD), ulcerative colitis (UC) (Franzosa *et al*. 2019), irritable bowel syndrome (IBS) (Jeffery *et al*. 2020), and end stage renal disease (ESRD) (Wang *et al*. 2020) (Supplementary Table S1). The global bacterial metabolic network was constructed using the KEGG database and pre-processed to remove high-degree currency metabolites, ensuring a biologically informative topology. The resulting network contained 2,172 EC nodes, 2,539 metabolite nodes and 6,253 edges, grouped into 13 connected components (Supplementary Figure S1). Additional details on network construction, data processing, and parameter sensitivity analysis are provided in the Supplementary Methods. For comparison, we also performed disease-association analyses for each omic independently, both at the individual feature level and using predefined metabolic pathways (see Supplementary Methods).

## 3 Results

### 3.1. Discovery of data-driven, disease-associated metabolic modules using MAAMOUL

We applied MAAMOUL to the four microbiome-metabolome datasets to identify custom metabolic modules associated with each disease. After processing the metagenomic and metabolomic data into functional and metabolite profiles, we assigned disease-association p-values to each EC and metabolite feature using generalized linear models (Supplementary Methods; Supplementary Table S2). When projected onto the global metabolic network, the coverage of observed nodes (i.e., nodes for which a disease-association p-value is assigned) varied across datasets, with between 41.6% to 46.2% for EC nodes, and only 1% to 3.9% for metabolite nodes (Supplementary Table S4). Despite this sparse metabolite mapping, MAAMOUL identified a total of 70 significant modules (FDR < 0.1) across the four conditions: 14 in CD, 14 in UC, 13 in IBS and 29 in ESRD. Of these, 9, 6, 2 and 0 modules, in each of the four studies respectively, included both anchor EC nodes *and* anchor metabolite nodes.

We compared MAAMOUL’s modules to two common approaches for identifying microbiome functional shifts associated with disease: differential abundance tests of individual features (i.e., specific ECs or metabolites), and over-representation analysis (ORA) at the metabolic pathway-level (Figure 2; Supplementary Methods). In the UC cohort, for example, 524 EC features and 50 KEGG-annotated metabolites were significantly associated with disease (Figure 2A). At the pathway level, ORA identified five pathways significantly enriched in UC-associated ECs, but none enriched in UC-associated metabolites (Figure 2B; Supplementary Table S8). The disease-associated pathways included phenylalanine, tyrosine, and tryptophan biosynthesis, lipopolysaccharide biosynthesis, glutathione metabolism, histidine metabolism, and lipoic acid metabolism. Notably, while disease-associated as a whole, the majority of features in these pathways were either unobserved or non-significant (Figure 2B).

**Fig. 2.**
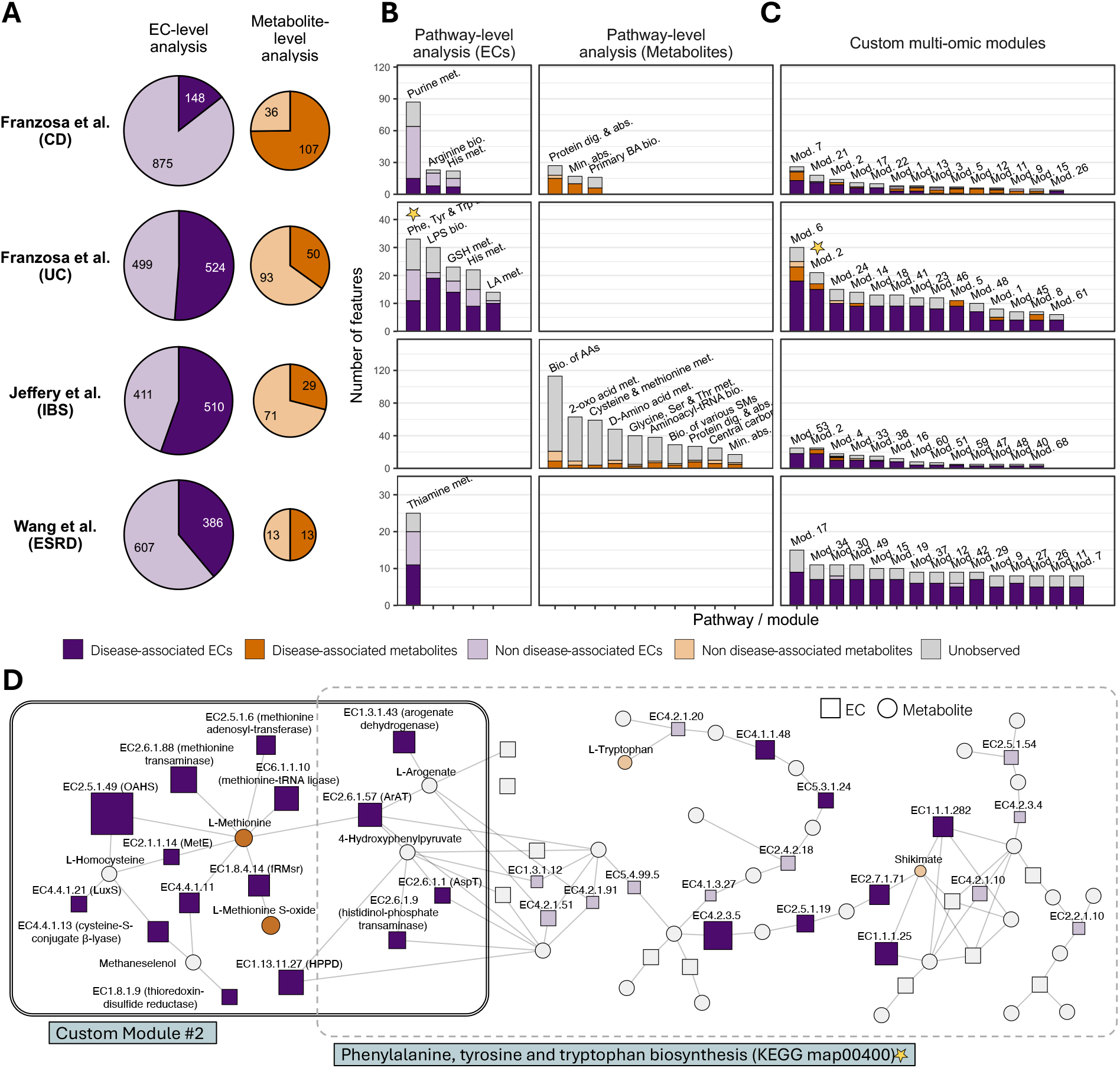
Single features, KEGG pathways, and custom metabolic modules associated with disease based on metagenomic and metabolomic data. **(A)** The number of significantly differentially abundant ECs based on fecal metagenomics (left), and KEGG-annotated metabolites based on fecal metabolomics (right). Pie sizes are proportional to the total number of features of each type, in each dataset. The method used for differential abundance testing, and the calculation of a p-value threshold for determining significance, are detailed in the Methods. **(B)** Disease-associated pathways based on either metagenomic EC profiles (left) or metabolomic profiles (right). Each bar represents a pathway that was significantly associated with the disease based on an over representation analysis (ORA) of significant ECs/metabolites. The dark portion of each bar represents ECs/metabolites that were significant in the pathway, the light portion represents non-significant ECs/metabolites in the pathway, and the grey portion represents ECs/metabolites that were unobserved in the data (i.e., their association with disease is unknown). **(C)** Significant disease-associated modules identified by MAAMOUL. Colors as in (B). In the ESRD dataset, only the top 15 modules are presented. **(D)** A part of the global metabolic network, with node colors and sizes reflecting associations with ulcerative colitis (UC) in the dataset from (Franzosa *et al*. 2019). This subgraph includes both the phenylalanine, tyrosine and tryptophan biosynthesis pathway (KEGG map00400), which is significantly associated with UC based on a simple ORA test, and a custom module identified by MAAMOUL that overlapped this pathway (see also Figure 3A). Nodes colors correspond to the same categories as in panels A-C and node sizes are inversely proportional to their p-values from differential abundance testing. Squares and circles represent EC and metabolite nodes, respectively. Abbreviations in pathway names: Phenylalanine: Phe; Tyrosine: Tyr; Tryptophan: Trp; Biosynthesis: bio.; metabolism: met.; Digestion: dig.; absorption: abs.; bile acids: BA; Lipopolysaccharide: LPS; Glutathione: GSH; Lipoic acid: LA; Amino acids: AAs; Mineral: Min.; Serine: Ser; Threonine: Thr; 2-Oxocarboxylic: 2-oxo; secondary metabolites: SMs; Histidine: His.

In contrast, MAAMOUL’s custom modules offer a more focused characterization of disease-related functional shifts by capturing network regions dense with disease-associated nodes (Figure 2C). In the UC cohort, MAAMOUL identified 14 significant modules, 6 of which included both anchor ECs and metabolites. Figure 2D illustrates this distinction, comparing the UC-associated phenylalanine, tyrosine, and tryptophan biosynthesis pathway (KEGG map00400) with custom Module #2 which partly overlaps this pathway. While only a third of the ECs and *none* of the metabolites in map00400 were significantly associated with UC, Module #2 involves both significant ECs and significant metabolites. Furthermore, this module spans both map00400 and the cysteine and methionine metabolism pathway, underscoring MAAMOUL’s ability to capture disease-associated functional units that cross canonical pathway boundaries.

To demonstrate the biological relevance of this approach, we next examine some of the detected modules that include both anchor ECs and anchor metabolites (which were detected in UC, CD, and IBS), as these provide the clearest examples of coordinated multi-omic shifts.

### 3.2. IBD-associated microbiome-metabolome modules

Analyzing the data from (Franzosa *et al*. 2019), MAAMOUL identified several significant modules associated with UC or CD. The UC-associated Module #2 (FDR = 0.02; Figure 2D), for example, reflects disrupted sulfur and aromatic amino acid metabolism, both of which are frequently altered in IBD (Lavelle and Sokol 2020, Metwaly *et al*. 2020, Tambovtseva *et al*. 2025). This module captures a depletion of L-methionine and reduced abundance of genes encoding for methionine biosynthetic and translational enzymes (such as EC 2.5.1.49 and EC 6.1.1.10). While it overlaps several KEGG pathways related to sulfur and aromatic amino acid metabolism (Figure 3A), the module is not fully contained within the boundaries of any single pathway, demonstrating MAAMOUL’s ability to detect functional modules that span several pre-defined pathways.

**Fig. 3.**
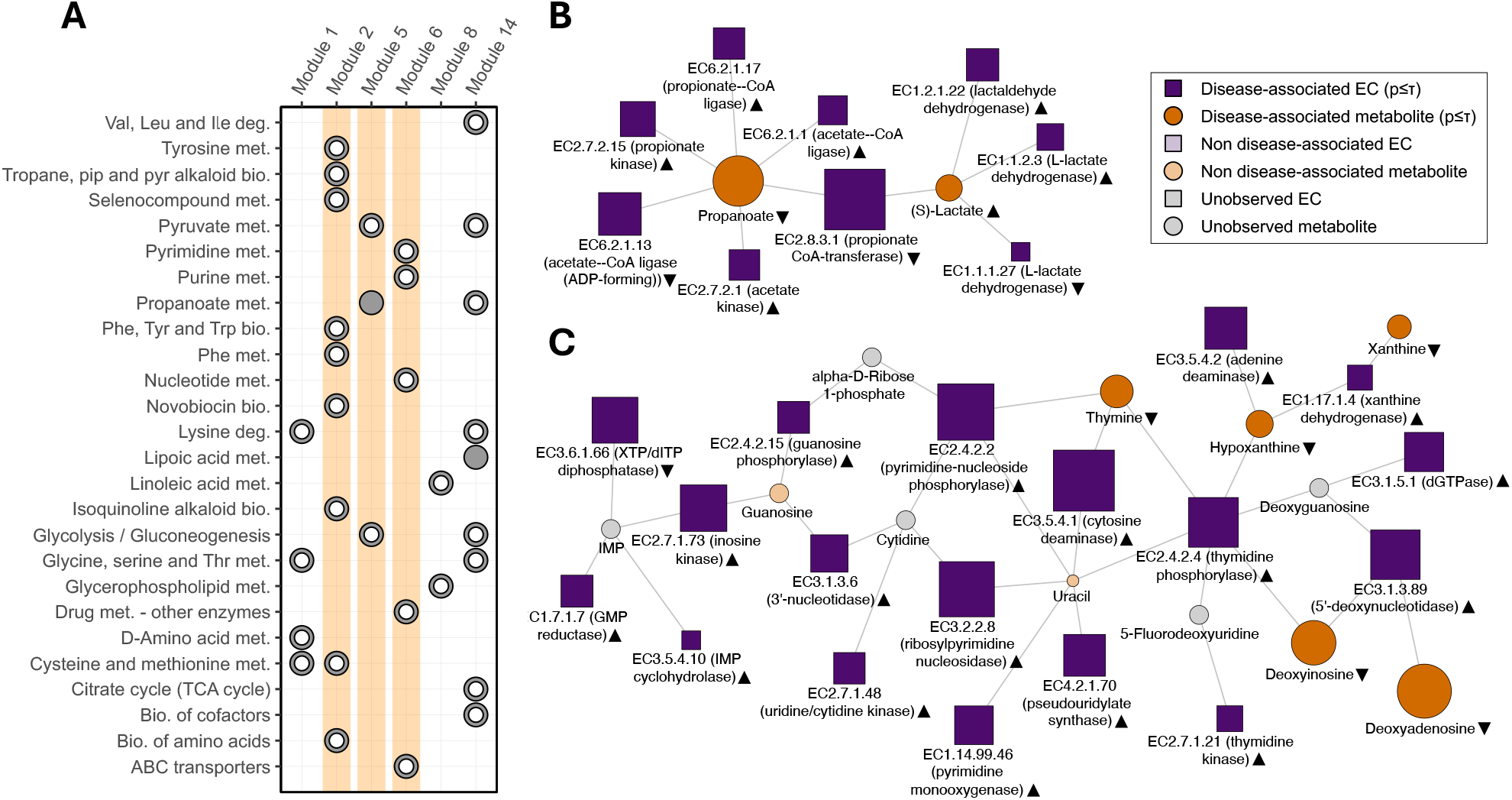
Microbiome-metabolome modules associated with ulcerative colitis (UC). **(A)** A comparison of custom modules associated with UC and identified by MAAMOUL, and KEGG metabolic pathways, as determined by a Fisher’s exact test (Methods). Each circle represents a significant overlap (FDR<0.05) between a module (columns) and a pathway (rows). Full points indicate that the module is mostly contained within the pathway (>80% of module nodes). Modules #2 (shown in Fig. 2D), #5 and #6 (shown in (B) and (C) respectively) are highlighted. **(B)**,**(C)** An illustration of UC-associated Module #5 and Module #6. Each module is a subgraph of the global metabolic network, with both enzyme nodes (squares) and metabolite nodes (circles), and with edges connecting each enzyme to its substrate and product metabolites. Nodes colored in dark shades indicate a significant association with disease, as determined using differential abundance tests. Grey-colored nodes represent unobserved features. Black triangles next to each node’s name indicate whether the corresponding feature was increased (triangles pointing up) or decreased (pointing down) in UC.

Other features in Module #2 are consistent with the oxidative burden characteristic of the inflamed UC colon. Specifically, the presence of L-methionine S-oxide and the repair enzyme methionine (R)-S-oxide reductase suggests an active microbial oxidative stress response (Ezraty *et al*. 2017, Walker and Schmitt-Kopplin 2021). Furthermore, the enrichment of transaminases (EC 2.6.1.88, EC 2.6.1.57) and sulfur lyases (EC 4.4.1.13) points toward metabolic reprogramming of the UC-associated microbiome toward a stress-adapted state (Ni *et al*. 2017). The module also includes homocysteine, a central intermediate in methionine cycling and transsulfuration (Bauchart-Thevret, Stoll, and Burrin 2009). Although homocysteine was not measured in this study, its inclusion in the module is consistent with reports linking elevated homocysteine levels to inflammatory activity in UC and IBD cohorts (Danese *et al*. 2005). This highlights how module-level analysis can capture biologically relevant intermediates even when they are absent from the initial assays.

Another UC-associated module (Module #5; FDR = 0.02; Figure 3B) highlights a metabolic shift from propionate to lactate. Propionate is an anti-inflammatory short-chain fatty acid typically produced from lactate via the acrylate pathway. However, in IBD, this conversion often breaks down (Lavelle and Sokol 2020). Module #5 reflects this disruption, showing decreased abundance of genes for propionate CoA-transferase and propionate-CoA ligase alongside an increase in lactate-producing dehydrogenases (EC 1.1.2.3) (Medina *et al*. 2021). Such shifts may be driven by lowered gut pH in severe UC (Vernia *et al*. 1988). Notably, while Module #5 is largely contained within the KEGG propanoate metabolism pathway, that pathway was not significant in standard single-omic analysis, illustrating MAAMOUL’s ability to detect localized perturbations masked at the pathway level.

Finally, UC-associated Module #6 (FDR = 0.02; Figure 3C) is centered on altered nucleotide metabolism. This module captures a depletion of nucleosides and nucleobases (e.g., xanthine, deoxyadenosine, and deoxyinosine) paired with an increased genomic potential for their degradation and salvage, involving enzymes such as GMP reductase and IMP cyclohydrolase. A highly similar module was detected in the CD cohort (Module #7; FDR = 0.03; Supplementary Figure S4A), indicating that this functional shift is characteristic of both IBD subtypes. While module compositions vary slightly between the two conditions, both reflect a microbiome adapting to an inflamed environment with high host cell turnover and the likely release of nucleotides into the gut lumen (Vincenzo Di *et al*. 2023). Module features are consistent with a dysbiotic microbiome that is rapidly scavenging these resources, potentially limiting the availability of salvageable precursors necessary for host epithelial renewal and mucosal repair (Lee *et al*. 2020).

### 3.3. IBS-associated microbiome-metabolome modules

Irritable bowel syndrome (IBS) is a highly heterogeneous and multifactorial condition with multiple disease subtypes and clinical manifestations (Singh *et al*. 2019). Although several studies have reported microbiome associations with IBS, symptom severity, or comorbidities (Pittayanon *et al*. 2019, Mars *et al*. 2020, Jacobs *et al*. 2023), these signatures are generally weaker than those observed in other gastrointestinal diseases, and underlying mechanisms remain largely unclear. Applying MAAMOUL to microbiome-metabolome data from (Jeffery *et al*. 2020) (with 78 IBS and 58 control samples), we found 13 IBS-associated modules (FDR<0.1), of which, two (Modules #2 and #4) were supported by both metabolite and EC features (Supplementary Figure S5). These modules highlighted shifts in amino acid utilization and proteolytic activity, as well as perturbations in purine and nicotinate metabolism, in line with previously hypothesized mechanisms. A detailed description of these modules and their biological interpretation is provided in Supplementary Note 1.

## 4 Discussion

In this work, we introduced MAAMOUL, a novel knowledge-based computational framework for identifying microbial metabolic modules perturbed in disease using paired metagenomic and metabolomic data. The method maps observed disease associations in these paired data onto a global bacterial metabolic network, and identifies data-driven disease-associated modules within this network, each representing a mechanistically interpretable hypothesis grounded in known biochemical relationships. In contrast to conventional functional analyses that rely on predefined metabolic pathways, MAAMOUL can detect shifts that may be missed by pathway-level approaches, including modules spanning several pathways or confined to a specific region within a broader pathway. To our knowledge, this is the first method to explicitly use metabolic networks to integrate metagenomic and metabolomic data in the context of host disease.

Applying MAAMOUL to microbiome-metabolome data from several case-control cohorts, we identified multiple disease-associated modules supported by both microbial genes and fecal metabolites. These included UC-associated modules reflecting disrupted sulfur and aromatic amino acid metabolism in oxidative stress, reduced conversion of lactate to propionate, and a nucleotide metabolism module (also found in CD). In IBS, MAAMOUL identified a module at the intersection of purine and nicotinate/nicotinamide metabolism, as well as a module reflecting shifts in amino acid utilization. Importantly, several of the highlighted modules represent coordinated functional shifts that were not detected using single-omic or pathway-level enrichment analyses.

Our approach has several key advantages over commonly used single-omic or pathway-level analyses. First, it allows flexibility in selecting statistical methods to test gene or metabolite associations with disease. In our analyses, we used linear models that accounted for cohort-specific confounders, but alternative methods could be applied for each omic. Nonetheless, alternative methods could have been used, including a different statistical test for each omic (as downstream modelling of p-value distributions is performed on each omic independently). This enables, for example, combining microbiome-specific differential abundance tests (Paulson *et al*. 2013), with statistical approaches tailored for metabolomics. Second, although developed for *paired* microbiome-metabolome datasets, the method does not require, in principle, matched samples, as p-values for each omic can be computed separately. This allows application to datasets with partial, or even no overlap between metagenomic and metabolomic samples. Another advantage is that MAAMOUL retains unobserved nodes in the metabolic network and assigns them a data-driven prior probability of disease association. This contrasts with methods such as Meta-Path (Liu and Pop 2011), which remove unscored nodes from the network. While this strategy may be reasonable for metagenomics data, it is problematic for metabolomics, where assays often measure only a subset of metabolites and many network metabolites remain unobserved. Removing these nodes can therefore result in a highly fragmented network.

Future work could further introduce several methodological improvements. For example, assigning p-values to unobserved nodes could consider their proximity to disease-associated nodes using network propagation or “guilt-by-association” approaches (Cowen *et al*. 2017). Anchor node clustering could also be improved by replacing hierarchical clustering with methods such as k-means or community detection algorithms (Thalamuthu *et al*. 2006). Lastly, a particularly promising direction is the identification of the taxonomic drivers of disease-associated metabolic modules, which would help distinguish shifts driven by a specific taxon from broader community-level shifts. This could be done using a “leave-one-species-out” approach or methods based on Shapley values (Manor and Borenstein 2017).

In summary, we suggest that integrating microbiome and metabolome data using biological knowledge offers advantages over purely statistical approaches. Such methods generate mechanistic hypotheses grounded in known biology and are likely less susceptible to noise or artifact. As reference databases and cohort sizes grow, these approaches will further advance our understanding of the gut microbiome’s role in human disease.

## Supplementary data

Supplementary data are available online.

## Data & code availability

Data used in this study were obtained from supplementary files and deposited data from previously published studies (see Supplementary Table S1). MAAMOUL is available as an R package at https://github.com/borenstein-lab/MAAMOUL. Scripts used for misc. analysis and for generating manuscript figures are available on GitHub at https://github.com/borenstein-lab/MAAMOUL_analysis.

## Acknowledgements

We would like to thank the Borenstein lab members for helpful feedback and discussions. We thank all authors of the studies whose data was included in our analysis, for generating invaluable data and making them publicly available.

