## Supplementary Information for "MAAMOUL: Metabolic network-based discovery of microbiome-metabolome shifts in disease"

### Supplementary data

Supplementary Figure S3: Sensitivity analysis – Cumulative size of output modules using different pipeline parameters .. 8

### Supplementary Note 1: IBS-associated microbiome-metabolome modules

As mentioned in the main text, we utilized MAAMOUL to investigate the metabolic shifts associated with Irritable Bowel Syndrome (IBS). IBS is a highly heterogeneous and multifactorial condition with multiple disease subtypes and clinical manifestations. Although several studies have reported microbiome taxonomic and functional associations with IBS symptom severity, or comorbidities[1–3], these signatures are generally weaker than those observed in other gastrointestinal diseases, and underlying mechanisms remain largely unclear. Applying MAAMOUL to microbiome-metabolome data from Jeffery et al.[4] (with 78 IBS and 58 control samples), we found 13 IBS-associated modules (FDR<0.1), of which, two (Modules #2 and #4) were supported by both metabolite and metagenomic EC features (Supplementary Figure 5).

Module #2 (Supplementary Figure 5A) includes 4 amino acids increased in IBS, namely lysine, glutamine, serine and tryptophan (Trp), alongside a decrease in arginine and multiple genes involved in amino acid biosynthesis and utilization. This pattern aligns with reports of elevated proteolytic activity in the IBS gut, where bacteria increasingly bypass carbohydrate fermentation in favor of protein breakdown[5,6]. Arginine depletion may reflect increased microbial utilization, consistent with enrichment of arginase and arginine decarboxylase genes, and potentially linked to the expansion of *Enterobacteriaceae* that utilize a "linear arginine pathway" to outcompete commensals[7]. Similarly, the enrichment of Trp and Trp-related genes (e.g., tryptophanase and tryptophan synthase) aligns well with previous implications of dysregulated Trp metabolism in IBS pathogenesis and symptoms[8,9], potentially via microbial modulation of Trp flux and activation of the host's kynurenine pathway, which in turn influence intestinal motility and visceral sensitivity in IBS[10].

Module #4 (Supplementary Figure 5B), in contrast, captures a different shift involving purine and nicotinate metabolism. It includes decreased levels of adenosine, adenine, and nicotinamide alongside increased abundance of multiple ECs overlapping the purine and nicotinate/nicotinamide pathways (Supplementary Table S7). Disturbances in purine metabolism have recently been proposed as a novel mechanism underlying IBS[11], potentially reflecting elevated microbial utilization of purine nucleotides that may contribute to epithelial metabolic stress and impaired mucosal repair. Consistent with this hypothesis, the module contains xanthine dehydrogenase genes (EC 1.17.1.4), previously reported to be elevated in IBS patients with predominant constipation[12]. The module also includes multiple enzymes at the intersection of purine metabolism and nicotinate/nicotinamide metabolism (e.g., EC 2.4.2.1), as well as nicotinamidase (EC 3.5.1.19), consistent with reduced fecal nicotinamide (a form of vitamin B3 that enters the gut lumen through circulating host nicotinamide[13]) and increased microbial conversion of nicotinamide to nicotinate in IBS[14].

In summary, these results demonstrate MAAMOUL's ability to identify mechanistically coherent, multi-omic functional units even within the heterogeneous landscape of IBS.

### Supplementary Methods

#### *Construction of the global metabolic network*

We constructed a metabolic network of enzyme nodes (represented by EC codes) and metabolite nodes (represented by KEGG compound ID's), using data from the KEGG database (FTP 2021-05-10 release)[15]. Notably, in contrast to the common metabolic network representation, where nodes represent metabolites and metabolic reactions are denoted by hyperedges, our representation includes metabolic reactions (coded by corresponding EC numbers) as nodes as well, resulting in a bipartite graph with two types of nodes (ECs and metabolites) and edges connecting EC nodes to their substrate and product metabolite nodes.

The network was constructed as follows: First, KEGG reaction ID's were mapped to their main substrates and products (given by compound ID's) using the file "reaction_mapformula.lst" from the "ligand" subdirectory. Then, each reaction was mapped to its corresponding EC (one or more) using the file "reaction_enzyme.list". Metabolite and enzyme names, as well as other properties, were extracted from the "compound" and "enzyme" files, respectively. The unique list of EC-metabolite pairs was then used as edges in the bipartite network. This preliminary global metabolic network included 3,793 enzyme nodes, 4,516 metabolite nodes, and 12,241 edges.

We further pre-processed the network by performing the following steps: Nodes with a rank (number of neighbors in the network) > 20 were removed from the network alongside all their connected edges. Overall, 27 metabolites were removed in this step (including, for example, Acetyl-CoA, ammonia, glucose and CO2), and 24 EC nodes. This removal of "currency" nodes (mostly metabolites) is a common practice in metabolic modeling[16], resulting in a network with a more informative topology. Singleton nodes were also removed. After this step, the network included 3,752 EC nodes, 4,321 metabolite nodes, and 10,697 edges. Next, EC nodes that were not linked in KEGG to any bacterial organism, as determined using the KEGG files "enzyme" ("ligand" subdirectory) and "taxonomic_rank" ("genes" subdirectory), were omitted from the network, as well as metabolite nodes that became singletons, resulting in 2,348 EC nodes and 2,861 metabolite nodes remaining in the graph. Lastly, connected components in the graph that included less than 10 nodes were removed. We refer to the resulting network, after the preprocessing steps described above, as the "global metabolic network", containing 2,172 EC nodes, 2,539 metabolite nodes and 6,253 edges, grouped into 13 connected components, the largest of which including 4,439 nodes. Supplementary Figure S1 presents node rank distributions before and after network processing, and a visualization of the final graph.

#### *Microbiome-metabolome datasets acquisition and processing*

In this work, we used data from the following studies: Franzosa et al.[17], Jeffery et al.[18] and Wang et al.[19]. For each cohort we obtained: (a) Metagenomics-based enzyme profiles; (b) Metabolite profiles as published or shared by the authors of the original studies; (c) Subject/sample metadata including demographics, disease state (healthy/disease), etc. One exception was the data from Jeffery et al., for which instead of metabolite profiles (that were unavailable), we obtained p-values per metabolite based on differential abundance testing from the original publication.

Enzyme profiles, given by Enzyme Commission (EC) numbers, were computed using HUMANn3[20], with default parameters, after quality filtering using the 'fastp' tool (version 0.23)[21], and host-reads removal using bowtie2 and the GRCh38 human reference genome (https://www.ncbi.nlm.nih.gov/datasets/genome/GCF_000001405.26), with the "sensitive" flag. As some of the EC codes obtained by HUMANn3 have been officially deprecated, we mapped deprecated EC numbers to their updated ones using the ENZYME database from the Expasy resource (https://ftp.expasy.org/databases/enzyme/enzyme.dat). The resulting EC profiles were normalized using MUSiCC[22]. Finally, rare EC's, defined as those observed in less than 10% of the samples were removed.

Metabolites were mapped, where possible, to their KEGG compound identifiers, either by the authors of the original studies or using the id conversion utility of MetaboAnalyst[23]. Metabolites that could not be mapped into KEGG identifiers were discarded. Rare metabolites, appearing in less than 10% of samples, were removed as well. Missing metabolite values were imputed using half the minimal observed value of each metabolite. Lastly, metabolite values were log-transformed to account for heteroscedasticity[24]. Further details and processing notes for each dataset, as well as accession codes for obtaining the raw data, are provided in Supplementary Table S1.

#### *Differential abundance analysis*

For each dataset, we ran differential abundance testing for each EC/metabolite feature using linear mixed models, as implemented in the "MaAsLin2" R package (version 1.8)[25]. Briefly, in each model, the EC/metabolite feature was the dependent variable, the study group (i.e. case/control label) was an independent (predictor) variable, and additional covariates/confounders, as controlled for in the original studies, were added as additional covariates. The p-value of the "study-group" variable coefficient was then recorded. The list of covariates used per dataset, the number of features (after pre-processing) that underwent testing, and additional statistics are given in Supplementary Table S2.

Notably, by using only p-values (rather than effect sizes), we focused solely on the significance of each feature's association with disease, ignoring the direction of association (i.e. whether the feature's abundance increased or decreased in disease). Our motivation for this choice was to relax the assumption that a functional module perturbed in disease is consistently upregulated or consistently downregulated, and rather allow identification of any module demonstrating shifts in activity in disease states. Similar methods for finding disease-associated modules in biological networks have adopted the same heuristic[26,27]. The method could, however, easily be adapted to find only consistent up-/down-regulated modules using one-sided statistical tests.

#### *MAAMOUL implementation, parameters and sensitivity analysis*

The MAAMOUL method is implemented in R (version 4.1). We use the "fitBumModel" function from the "BioNet" R package (version 1.54)[28] to obtain maximum-likelihood estimations for mixture model parameters, and the R package "igraph" (version 1.3.4)[29] for miscellaneous graph-related computations.

The method requires three main parameters: (a) An FDR threshold, determining the threshold used for defining which nodes will be considered as module "anchors" (default: 0.1); (b) $k$, determining a maximal distance between nodes for them to be considered as being part of the same disease-associated module (default: 4); and (c) The height at which we "cut" the tree obtained by hierarchical clustering of all anchor nodes (default: 0.8).

To assess how different parameter settings impact the identified modules, we ran MAAMOUL with several different combinations of parameter values (FDR threshold parameter, and $k$ parameter; see Supplementary Figure S3). As expected, less stringent FDR thresholds, and higher levels of $k$, both resulted in an overall increase in the number of nodes included in the obtained modules. Importantly, however, across all datasets, and across all parameter settings, the cumulative size of all obtained modules was significantly higher compared to modules identified in node-permuted graphs. Overall, different settings may produce more/ fewer modules, of increased/decreased size, and parameter choice should be made based on data characteristics and desired sensitivity. The parameters chosen for the analysis presented (which are the default parameters described above) were selected to maintain a balance between discovery and stringency.

##### *Identification of overlaps between modules and KEGG pathways*

For each MAAMOUL-identified module (complete modules, not only anchor nodes), we identified all the KEGG-pathways that it significantly overlapped. To this end, we created a table mapping each node in the global metabolic network (either EC or metabolite) to all pathways in which they appear in KEGG, using the tables "enzyme_pathway.list" and "compound_pathway.list" from the "ligand" KEGG directory. Per module, we identified all pathways for which an overlap of at least 3 nodes existed, and then used a hypergeometric test to obtain an "enrichment p-value". P-values were then FDR-corrected. All overlaps with an FDR < 0.1 are listed in Supplementary Table S7, and visualized in Figures 3B and S4B.

##### *Pathway-based analysis*

To compare the custom modules identified by MAAMOUL to the common practice of pathway-based analysis, we implemented an Over Representation Analysis (ORA). Specifically, we used the same categorization of EC and metabolite features into significantly disease-associated (i.e., "anchors" in MAAMOUL's terminology) and non-disease-associated. We then used a hypergeometric test to examine, for each pathway, whether it was significantly enriched with disease associated ECs/metabolites. For consistency, for each KEGG pathway we used only the nodes that were also included in our global network (removing, for example, currency metabolites).

For metagenomic data only, we also report results from a standard differential abundance test, but applied at the whole pathway level. Specifically, the abundance of each pathway was computed as the sum of the relative abundances of all the ECs in that pathway, where the abundance of an EC that was included in multiple pathways is evenly distributed among them (the "uniform fractional mapping" approach[30]). We then used a linear mixed model (as described above for the individual features) to determine whether the abundance of each pathway was significantly different in cases versus controls. All results from these pathway-level analyses are provided in Supplementary Table S8.

##### *Visualizations*

Network visualizations were created using the R package "igraph" (version 1.3.4)[29], the RCy3 package (version 2.26.0)[31], and the Cytoscape software (version 3.10.2)[32].

### Supplementary Figures

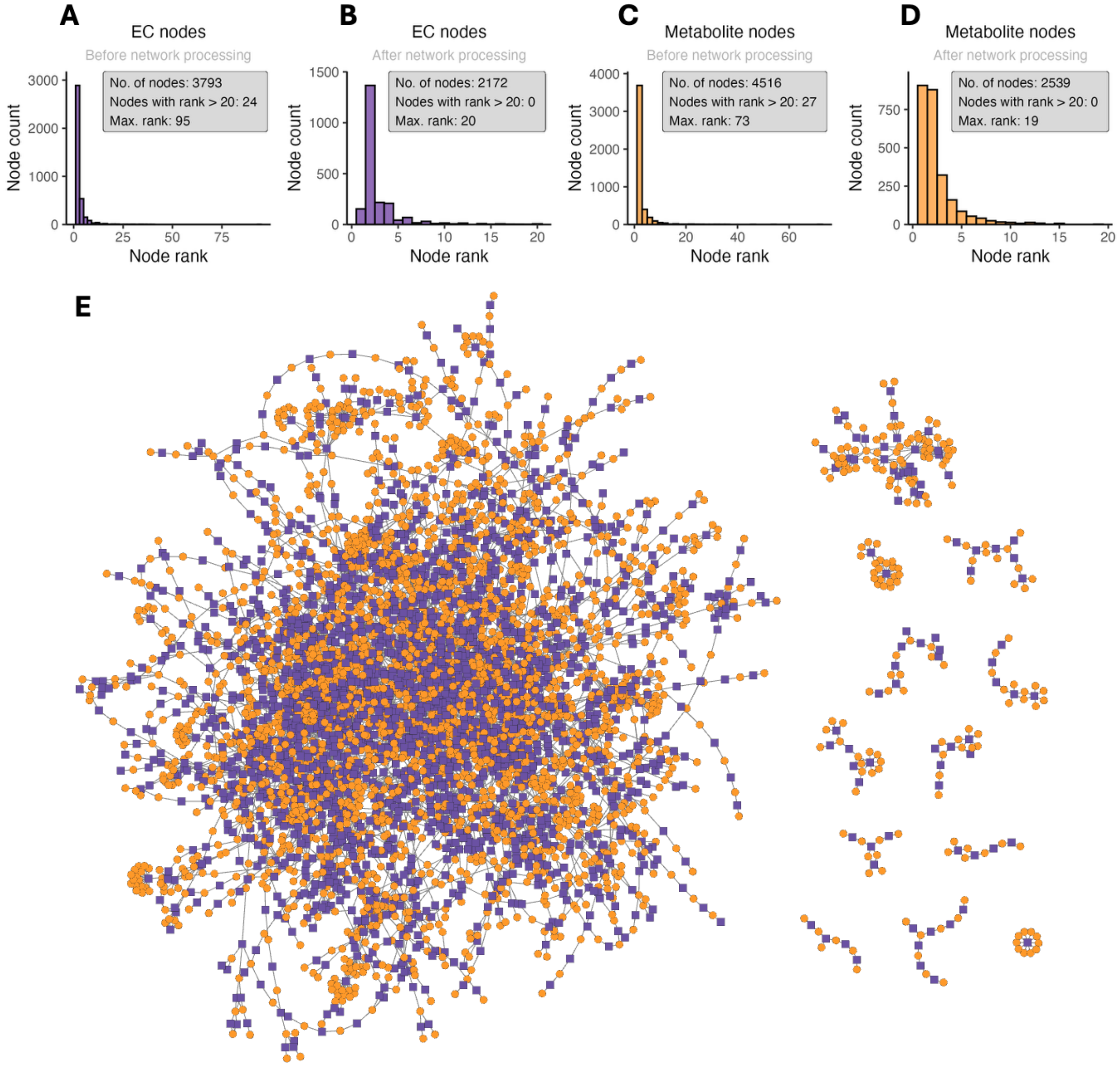

*Supplementary Figure S1: A bipartite global metabolic network*

**(A-D)** The distribution of node degrees (ranks) in the global bipartite network. Left panels (A,C) describe EC nodes, right panels (B, D) describe metabolite nodes, upper panels (A, B) describe the full network before any processing, and the lower panels (C, D) describe the final network after various processing steps; **(E)** A cytoscape visualization of the entire bipartite global metabolic network after processing.

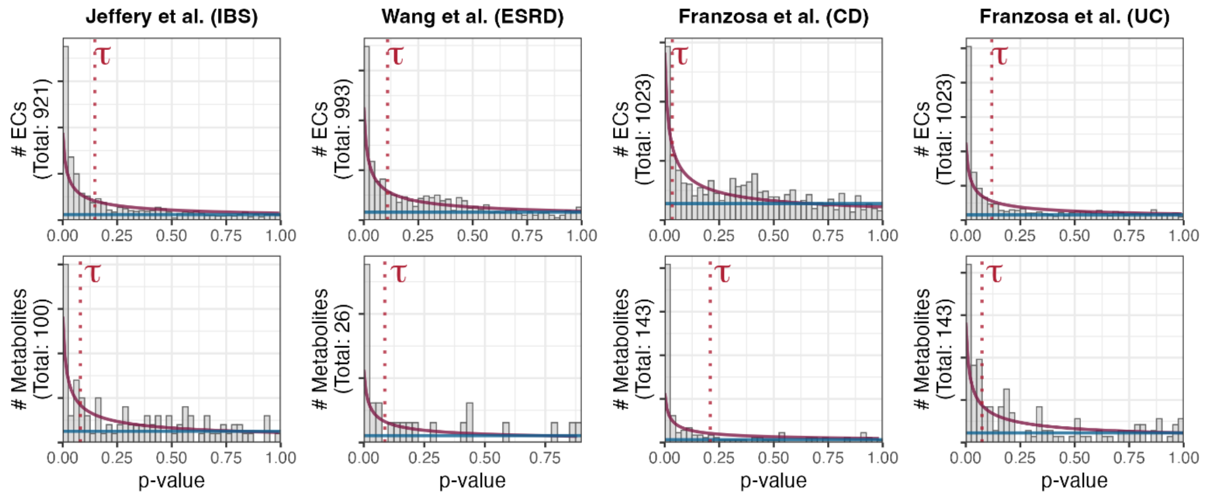

*Supplementary Figure S2: Distribution of p-values from differential abundance analysis and the corresponding fitted mixture models*

For each dataset, the distribution of p-values from the differential abundance testing (Methods) is presented, for EC features (upper panels) and metabolite features (lower panels) separately. Purple and blue overlaying lines illustrate the fitted beta distribution and uniform distribution, respectively, of the beta-uniform mixture (BUM) models. Red dashed line represent the selected p-value cutoff for defining "anchor" nodes, $\tau$, signifying a false discovery rate of 0.1 and using the formulation by Pounds and Morris[33].

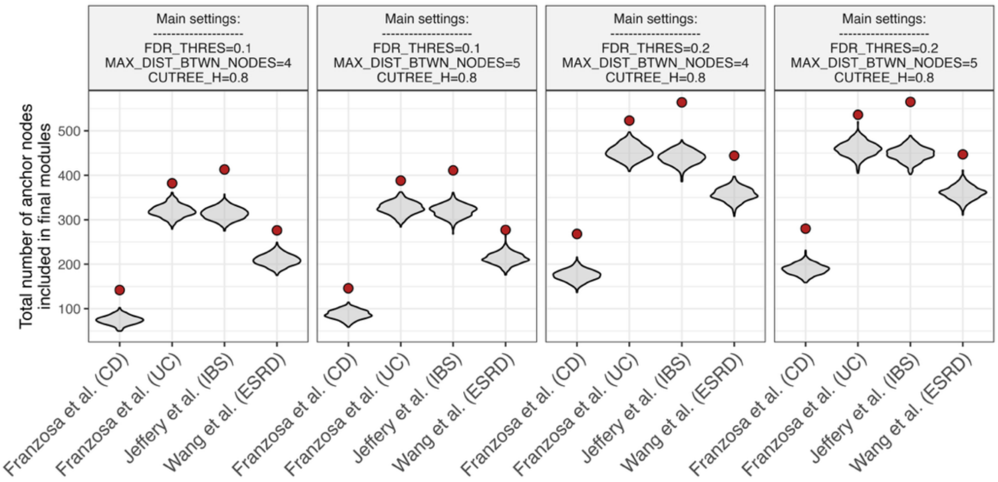

*Supplementary Figure S3: Sensitivity analysis – Cumulative size of output modules using different pipeline parameters*

Each panel represents a different set of pipeline parameters. See Methods for parameter descriptions. For each dataset (x axis), the red point represents the cumulative number of nodes included in the identified modules (regardless of the modules' statistical significance as determined by the topology-aware permutation test). Grey violins represent the distribution of cumulative numbers of nodes for node-permuted networks, as a null distribution.

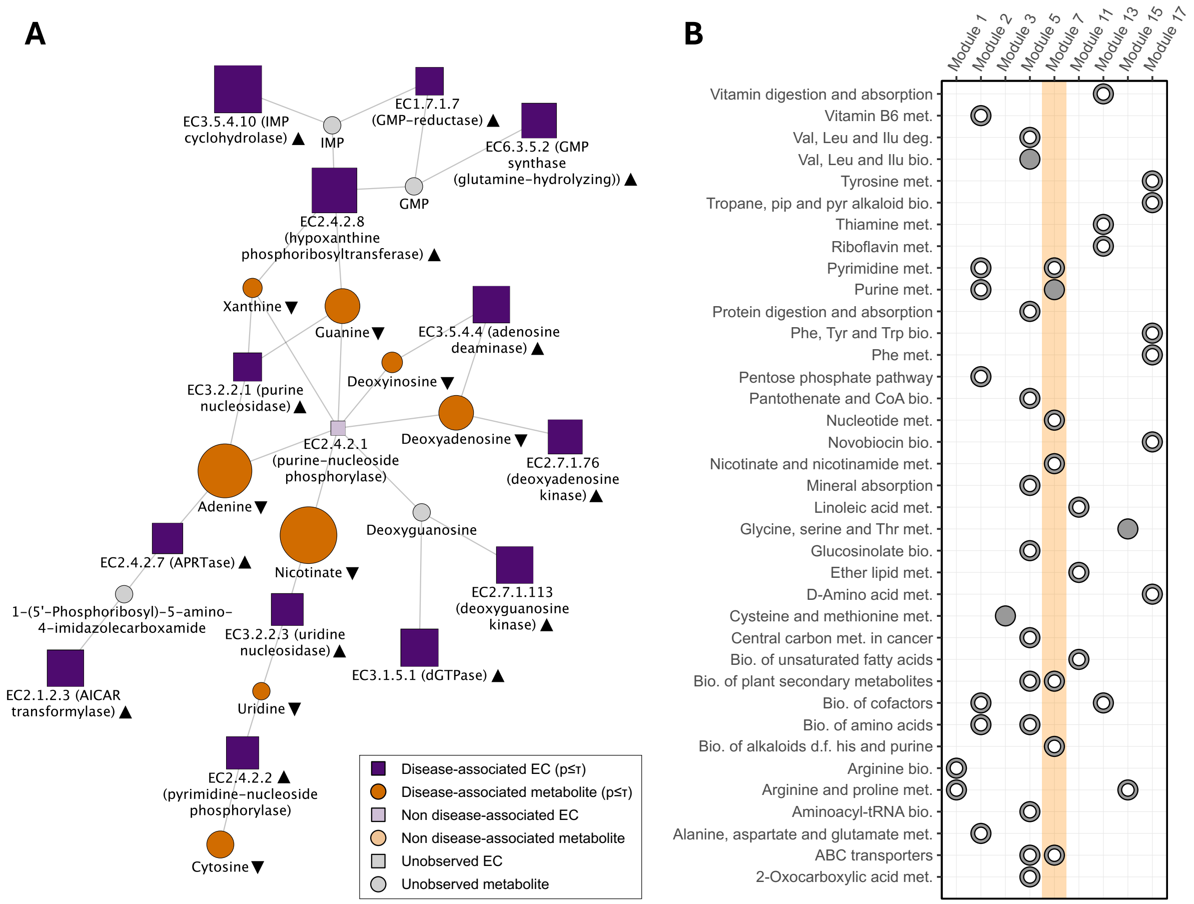

#### *Supplementary Figure S4: Microbiome-metabolome modules associated with Crohn's disease*

**(A)** An illustration of Module #7 identified by MAAMOUL using data from Crohn's disease (CD) patients and healthy controls from Franzosa et al.[17]. (B) Significant overlaps between CD-associated modules identified by MAAMOUL (columns) and pre-defined KEGG metabolic pathways (rows). Overlaps with Module #7 presented in panel (A) are highlighted. See legend of Figure 3.

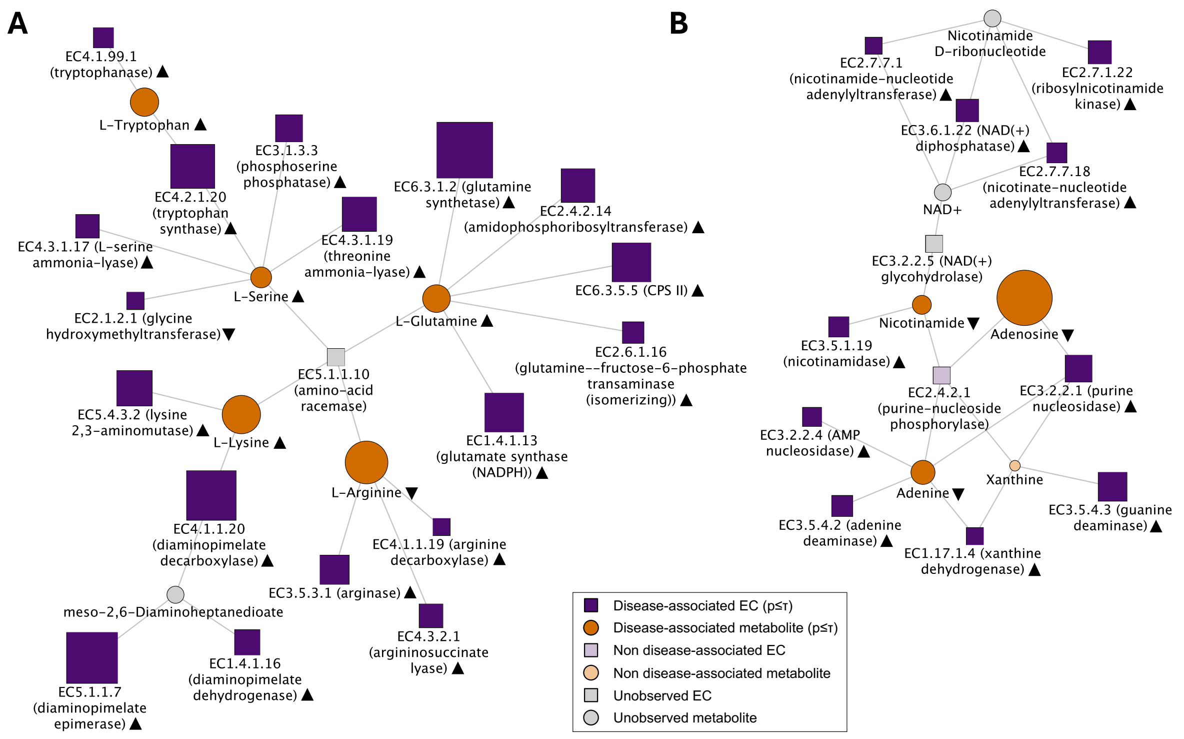

*Supplementary Figure S50: Microbiome functional shifts associated with IBS*

An illustration of Module #2 **(A)**, and Module #4 **(B)** identified by MAAMOUL applied to the IBS dataset from Jeffery et al.[18]. See legend of Figure 3.

### Supplementary Tables

#### *Supplementary Table S1: Description of studies included in the analysis*

| Dataset | Cohort description | No. samples | | Microbiome data | Metabolome data | | Metadata location | Publication | Comments |
|---|---|---|---|---|---|---|---|---|---|
| | | Control | Case | Raw data location | Instrument | Data location | | | |
| Franzosa et al. (CD) | Inflammatory bowel disease patients and controls (PRISM cohort). - CD and healthy only. | 56 | 73 | BioProject: PRJNA400072 / Paper SI | Four complimentary LC-MS methods | Paper SI | Paper SI | DOI: 10.1038/s41564-018-0306-4 | The dataset includes two different cohorts: the main PRISM cohort (from Boston, USA) and a validation cohort from the Netherlands. |
| Franzosa et al. (UC) | Inflammatory bowel disease patients and controls (PRISM cohort). - UC and healthy only. | 56 | 70 | BioProject: PRJNA400072 / Paper SI | Four complimentary LC-MS methods | Paper SI | Paper SI | DOI: 10.1038/s41564-018-0306-4 | The dataset includes two different cohorts: the main PRISM cohort (from Boston, USA) and a validation cohort from the Netherlands. |
| Wang et al. (ESRD) | Adults with end-stage renal disease (ESRD) and controls. | 67 | 220 | BioProject: PRJNA449784 | GC-MS | MetaboLights database: accession number MTBLS700 | MetaboLights database: accession number MTBLS700 | DOI: 10.1136/gutjnl-2019-319766 | |
| Jeffery et al. (IBS) | Adults with and without IBS (Rome IV criteria) | 58 | 78 | BioProject: PRJEB42304 | LC-MS | Paper SI | Shared by authors | DOI: 10.1053/j.gastro.2019.11.301 | Metabolite levels were unavailable. Instead, pre-computed differential abundance p-values were obtained from supplementary tables. |

#### *Supplementary Table S2: Differential abundance analysis statistics per dataset*

| Dataset | Fixed effects for DA analysis | Feature type | No. of samples | No. of features | No. of p-values < 0.05 | No. of FDR values < 0.05 | No. of p-values < 0.1 | No. of FDR values < 0.1 |
|---|---|---|---|---|---|---|---|---|
| Wang et al. (ESRD) | Age, Gender, BMI | ECs | 287 | 993 | 289 | 87 | 388 | 123 |
| Wang et al. (ESRD) | Age, Gender, BMI | Metabolites | 287 | 26 | 11 | 6 | 13 | 7 |
| Franzosa et al. (CD) | Age, immunosuppressant, mesalamine, steroids, Batch | ECs | 129 | 1023 | 206 | 19 | 281 | 34 |
| Franzosa et al. (CD) | Age, immunosuppressant, mesalamine, steroids, Batch | Metabolites | 129 | 143 | 84 | 48 | 93 | 66 |
| Franzosa et al. (UC) | Age, immunosuppressant, mesalamine, steroids, Batch | ECs | 126 | 1023 | 412 | 6 | 513 | 41 |
| Franzosa et al. (UC) | Age, immunosuppressant, mesalamine, steroids, Batch | Metabolites | 126 | 143 | 43 | 12 | 63 | 19 |
| Jeffery et al. (IBS) | | ECs | 136 | 921 | 353 | 190 | 462 | 291 |
| Jeffery et al. (IBS) | | Metabolites | 139 | 100 | 23 | 15 | 35 | 23 |

* Note 1: Numbers include all observed features in the data, after pre-processing (e.g. removal of rare features). Some of these features may have been missing from the global network (see Table S4).
* Note 2: Significance thresholds in this table are based on standard FDR thresholds, and not on the mixture models (see Table S3).

#### *Supplementary Table S3: Beta-uniform mixture model parameters per dataset.*

| Dataset | Feature type | BUM parameter: a (shape) | BUM parameter: $\lambda$ (mixture) | Pipeline parameter: FDR threshold | Final p-value threshold based on BUM | Percent of features below p-value threshold (“anchors”) |
|---|---|---|---|---|---|---|
| Jeffery et al. (IBS) | EC | 0.3579 | 0 | 0.1 | 0.1372 | 54.23 |
| Jeffery et al. (IBS) | Metabolite | 0.2719 | 0.3156 | 0.1 | 0.0709 | 34.8 |
| Wang et al. (ESRD) | EC | 0.4002 | 0 | 0.1 | 0.0991 | 38.74 |
| Wang et al. (ESRD) | Metabolite | 0.1527 | 0.4463 | 0.1 | 0.077 | 38.04 |
| Franzosa et al. (CD) | EC | 0.5161 | 0.086 | 0.1 | 0.0246 | 14.22 |
| Franzosa et al. (CD) | Metabolite | 0.1955 | 0.1597 | 0.1 | 0.1989 | 65.52 |
| Franzosa et al. (UC) | EC | 0.3876 | 0 | 0.1 | 0.1094 | 50.13 |
| Franzosa et al. (UC) | Metabolite | 0.4507 | 0 | 0.1 | 0.0645 | 31.61 |

* BUM = Beta-uniform mixture model

#### *Supplementary Table S4: Metabolic network statistics per dataset*

| Dataset | No. of metabolite nodes with a p-value | No. of EC nodes with a p-value | No. of 'anchor' metabolite nodes | No. of 'anchor' EC nodes | No. of metabolite nodes in global network | No. of EC nodes in global network |
|---|---|---|---|---|---|---|
| Wang et al. (ESRD) | 26 | 977 | 16 | 547 | 2539 | 2172 |
| Franzosa et al. (CD) | 99 | 1004 | 66 | 255 | 2539 | 2172 |
| Franzosa et al. (UC) | 99 | 1004 | 31 | 499 | 2539 | 2172 |
| Jeffery et al. (IBS) | 98 | 904 | 29 | 393 | 2539 | 2172 |

### *Supplementary Table S5: Modules overview per dataset*

| Dataset | Module ID | Module p-value (network permutation test) | Module FDR (network permutation test) | No. of DA nodes in module | No. of DA metabolites in module | No. of DA ECs in module | Full module size |
|---|---|---|---|---|---|---|---|
| Franzosa et al. (CD) | 1 | 0.036 | **0.0780** | 5 | 2 | 3 | 8 |
| Franzosa et al. (CD) | 2 | 0.004 | **0.0347** | 11 | 2 | 9 | 14 |
| Franzosa et al. (CD) | 3 | 0.006 | **0.0390** | 4 | 3 | 1 | 7 |
| Franzosa et al. (CD) | 4 | 0.064 | 0.1040 | 8 | 3 | 5 | 11 |
| Franzosa et al. (CD) | 5 | 0.026 | **0.0662** | 5 | 4 | 1 | 7 |
| Franzosa et al. (CD) | 6 | 0.17 | 0.2009 | 4 | 2 | 2 | 7 |
| Franzosa et al. (CD) | 7 | 0.002 | **0.0347** | 21 | 8 | 13 | 26 |
| Franzosa et al. (CD) | 8 | 0.302 | 0.3141 | 3 | 1 | 2 | 5 |
| Franzosa et al. (CD) | 9 | 0.004 | **0.0347** | 3 | 3 | 0 | 5 |
| Franzosa et al. (CD) | 10 | 0.274 | 0.2968 | 3 | 1 | 2 | 6 |
| Franzosa et al. (CD) | 11 | 0.012 | **0.0446** | 4 | 3 | 1 | 6 |
| Franzosa et al. (CD) | 12 | 0.028 | **0.0662** | 5 | 5 | 0 | 6 |
| Franzosa et al. (CD) | 13 | 0.01 | **0.0433** | 6 | 3 | 3 | 8 |
| Franzosa et al. (CD) | 14 | 0.06 | 0.1040 | 5 | 1 | 4 | 9 |
| Franzosa et al. (CD) | 15 | 0.028 | **0.0662** | 3 | 2 | 1 | 5 |
| Franzosa et al. (CD) | 16 | 0.092 | 0.1259 | 4 | 1 | 3 | 5 |
| Franzosa et al. (CD) | 17 | 0.042 | **0.0840** | 7 | 1 | 6 | 11 |
| Franzosa et al. (CD) | 18 | 0.214 | 0.2419 | 4 | 1 | 3 | 5 |
| Franzosa et al. (CD) | 19 | 0.078 | 0.1193 | 4 | 0 | 4 | 6 |
| Franzosa et al. (CD) | 20 | 0.136 | 0.1684 | 3 | 0 | 3 | 6 |
| Franzosa et al. (CD) | 21 | 0.018 | **0.0585** | 11 | 0 | 11 | 18 |
| Franzosa et al. (CD) | 22 | 0.008 | **0.0416** | 6 | 0 | 6 | 10 |
| Franzosa et al. (CD) | 23 | 0.086 | 0.1242 | 3 | 0 | 3 | 7 |
| Franzosa et al. (CD) | 24 | 0.346 | 0.3460 | 3 | 0 | 3 | 5 |
| Franzosa et al. (CD) | 25 | 0.124 | 0.1612 | 4 | 0 | 4 | 7 |
| Franzosa et al. (CD) | 26 | 0.046 | **0.0854** | 3 | 0 | 3 | 4 |
| Franzosa et al. (UC) | 1 | 0.008 | **0.0596** | 5 | 1 | 4 | 8 |
| Franzosa et al. (UC) | 2 | 0.002 | **0.0223** | 17 | 2 | 15 | 21 |
| Franzosa et al. (UC) | 3 | 0.07 | 0.2315 | 5 | 1 | 4 | 8 |
| Franzosa et al. (UC) | 4 | 0.28 | 0.3633 | 3 | 1 | 2 | 5 |
| Franzosa et al. (UC) | 5 | 0.002 | **0.0223** | 11 | 2 | 9 | 11 |
| Franzosa et al. (UC) | 6 | 0.002 | **0.0223** | 23 | 5 | 18 | 30 |
| Franzosa et al. (UC) | 7 | 0.282 | 0.3633 | 4 | 1 | 3 | 5 |
| Franzosa et al. (UC) | 8 | 0.02 | **0.0957** | 6 | 2 | 4 | 7 |
| Franzosa et al. (UC) | 9 | 0.352 | 0.4288 | 3 | 1 | 2 | 5 |
| Franzosa et al. (UC) | 10 | 0.156 | 0.2773 | 11 | 1 | 10 | 16 |
| Franzosa et al. (UC) | 11 | 0.34 | 0.4219 | 4 | 1 | 3 | 5 |
| Franzosa et al. (UC) | 12 | 0.526 | 0.5594 | 3 | 1 | 2 | 5 |
| Franzosa et al. (UC) | 13 | 0.484 | 0.5230 | 3 | 1 | 2 | 3 |
| Franzosa et al. (UC) | 14 | 0.01 | **0.0670** | 10 | 1 | 9 | 14 |
| Franzosa et al. (UC) | 15 | 0.124 | 0.2773 | 5 | 0 | 5 | 7 |
| Franzosa et al. (UC) | 16 | 0.038 | 0.1675 | 4 | 0 | 4 | 6 |
| Franzosa et al. (UC) | 17 | 0.208 | 0.2959 | 3 | 0 | 3 | 6 |
| Franzosa et al. (UC) | 18 | 0.002 | **0.0223** | 9 | 0 | 9 | 13 |
| Franzosa et al. (UC) | 19 | 0.436 | 0.4869 | 3 | 0 | 3 | 5 |
| Franzosa et al. (UC) | 20 | 0.368 | 0.4326 | 5 | 0 | 5 | 9 |
| Franzosa et al. (UC) | 21 | 0.658 | 0.6580 | 3 | 0 | 3 | 5 |
| Franzosa et al. (UC) | 22 | 0.156 | 0.2773 | 4 | 0 | 4 | 5 |
| Franzosa et al. (UC) | 23 | 0.002 | **0.0223** | 9 | 0 | 9 | 12 |
| Franzosa et al. (UC) | 24 | 0.014 | **0.0782** | 10 | 0 | 10 | 15 |
| Franzosa et al. (UC) | 25 | 0.25 | 0.3418 | 3 | 0 | 3 | 5 |
| Franzosa et al. (UC) | 26 | 0.194 | 0.2954 | 12 | 0 | 12 | 16 |
| Franzosa et al. (UC) | 27 | 0.074 | 0.2315 | 13 | 0 | 13 | 17 |
| Franzosa et al. (UC) | 28 | 0.124 | 0.2773 | 3 | 0 | 3 | 4 |
| Franzosa et al. (UC) | 29 | 0.148 | 0.2773 | 4 | 0 | 4 | 6 |
| Franzosa et al. (UC) | 30 | 0.154 | 0.2773 | 4 | 0 | 4 | 7 |
| Franzosa et al. (UC) | 31 | 0.388 | 0.4482 | 3 | 0 | 3 | 4 |
| Franzosa et al. (UC) | 32 | 0.076 | 0.2315 | 23 | 0 | 23 | 30 |
| Franzosa et al. (UC) | 33 | 0.59 | 0.5989 | 3 | 0 | 3 | 5 |
| Franzosa et al. (UC) | 34 | 0.168 | 0.2773 | 3 | 0 | 3 | 5 |
| Franzosa et al. (UC) | 35 | 0.36 | 0.4307 | 4 | 0 | 4 | 5 |
| Franzosa et al. (UC) | 36 | 0.202 | 0.2959 | 7 | 0 | 7 | 9 |
| Franzosa et al. (UC) | 37 | 0.118 | 0.2773 | 5 | 0 | 5 | 7 |
| Franzosa et al. (UC) | 38 | 0.176 | 0.2773 | 3 | 0 | 3 | 5 |
| Franzosa et al. (UC) | 39 | 0.06 | 0.2233 | 7 | 0 | 7 | 12 |
| Franzosa et al. (UC) | 40 | 0.456 | 0.5009 | 3 | 0 | 3 | 5 |
| Franzosa et al. (UC) | 41 | 0.002 | **0.0223** | 9 | 0 | 9 | 13 |
| Franzosa et al. (UC) | 42 | 0.11 | 0.2773 | 4 | 0 | 4 | 7 |
| Franzosa et al. (UC) | 43 | 0.148 | 0.2773 | 3 | 0 | 3 | 5 |
| Franzosa et al. (UC) | 44 | 0.258 | 0.3457 | 3 | 0 | 3 | 6 |
| Franzosa et al. (UC) | 45 | 0.014 | **0.0782** | 4 | 0 | 4 | 7 |
| Franzosa et al. (UC) | 46 | 0.004 | **0.0383** | 8 | 0 | 8 | 12 |
| Franzosa et al. (UC) | 47 | 0.16 | 0.2773 | 4 | 0 | 4 | 5 |
| Franzosa et al. (UC) | 48 | 0.006 | **0.0503** | 7 | 0 | 7 | 10 |
| Franzosa et al. (UC) | 49 | 0.04 | 0.1675 | 4 | 0 | 4 | 6 |
| Franzosa et al. (UC) | 50 | 0.172 | 0.2773 | 4 | 0 | 4 | 7 |
| Franzosa et al. (UC) | 51 | 0.05 | 0.1971 | 7 | 0 | 7 | 10 |
| Franzosa et al. (UC) | 52 | 0.17 | 0.2773 | 4 | 0 | 4 | 6 |
| Franzosa et al. (UC) | 53 | 0.17 | 0.2773 | 3 | 0 | 3 | 4 |
| Franzosa et al. (UC) | 54 | 0.308 | 0.3894 | 3 | 0 | 3 | 4 |
| Franzosa et al. (UC) | 55 | 0.178 | 0.2773 | 5 | 0 | 5 | 7 |
| Franzosa et al. (UC) | 56 | 0.56 | 0.5863 | 5 | 0 | 5 | 8 |
| Franzosa et al. (UC) | 57 | 0.426 | 0.4838 | 3 | 0 | 3 | 5 |
| Franzosa et al. (UC) | 58 | 0.212 | 0.2959 | 5 | 0 | 5 | 7 |
| Franzosa et al. (UC) | 59 | 0.148 | 0.2773 | 5 | 0 | 5 | 8 |
| Franzosa et al. (UC) | 60 | 0.17 | 0.2773 | 3 | 0 | 3 | 5 |
| Franzosa et al. (UC) | 61 | 0.02 | **0.0957** | 4 | 0 | 4 | 6 |
| Franzosa et al. (UC) | 62 | 0.13 | 0.2773 | 5 | 0 | 5 | 8 |

| | | | | | | | |
|---|---|---|---|---|---|---|---|
| Franzosa et al. (UC) | 63 | 0.072 | 0.2315 | 3 | 0 | 3 | 4 |
| Franzosa et al. (UC) | 64 | 0.156 | 0.2773 | 4 | 0 | 4 | 6 |
| Franzosa et al. (UC) | 65 | 0.1 | 0.2773 | 4 | 0 | 4 | 6 |
| Franzosa et al. (UC) | 66 | 0.21 | 0.2959 | 3 | 0 | 3 | 4 |
| Franzosa et al. (UC) | 67 | 0.59 | 0.5989 | 3 | 0 | 3 | 4 |
| Jeffery et al. (IBS) | 1 | 0.254 | 0.3651 | 5 | 1 | 4 | 7 |
| Jeffery et al. (IBS) | 2 | 0.002 | **0.0345** | 23 | 5 | 18 | 25 |
| Jeffery et al. (IBS) | 3 | 0.048 | 0.1274 | 12 | 1 | 11 | 15 |
| Jeffery et al. (IBS) | 4 | 0.002 | **0.0345** | 13 | 3 | 10 | 18 |
| Jeffery et al. (IBS) | 5 | 0.512 | 0.5791 | 8 | 1 | 7 | 11 |
| Jeffery et al. (IBS) | 6 | 0.1 | 0.2156 | 4 | 2 | 2 | 6 |
| Jeffery et al. (IBS) | 7 | 0.314 | 0.4167 | 5 | 1 | 4 | 5 |
| Jeffery et al. (IBS) | 8 | 0.792 | 0.8280 | 3 | 1 | 2 | 3 |
| Jeffery et al. (IBS) | 9 | 0.194 | 0.3187 | 7 | 1 | 6 | 9 |
| Jeffery et al. (IBS) | 10 | 0.642 | 0.7031 | 4 | 1 | 3 | 5 |
| Jeffery et al. (IBS) | 11 | 0.394 | 0.4904 | 4 | 1 | 3 | 7 |
| Jeffery et al. (IBS) | 12 | 0.832 | 0.8568 | 5 | 1 | 4 | 7 |
| Jeffery et al. (IBS) | 13 | 0.094 | 0.2137 | 6 | 0 | 6 | 9 |
| Jeffery et al. (IBS) | 14 | 0.246 | 0.3651 | 3 | 0 | 3 | 5 |
| Jeffery et al. (IBS) | 15 | 0.032 | 0.1227 | 26 | 0 | 26 | 36 |
| Jeffery et al. (IBS) | 16 | 0.01 | **0.0767** | 8 | 0 | 8 | 12 |
| Jeffery et al. (IBS) | 17 | 0.464 | 0.5336 | 6 | 0 | 6 | 9 |
| Jeffery et al. (IBS) | 18 | 0.042 | 0.1270 | 4 | 0 | 4 | 7 |
| Jeffery et al. (IBS) | 19 | 0.046 | 0.1270 | 9 | 0 | 9 | 15 |
| Jeffery et al. (IBS) | 20 | 0.096 | 0.2137 | 16 | 0 | 16 | 22 |
| Jeffery et al. (IBS) | 21 | 0.038 | 0.1270 | 11 | 0 | 11 | 16 |
| Jeffery et al. (IBS) | 22 | 0.406 | 0.4915 | 9 | 0 | 9 | 12 |
| Jeffery et al. (IBS) | 23 | 0.046 | 0.1270 | 7 | 0 | 7 | 11 |
| Jeffery et al. (IBS) | 24 | 0.082 | 0.1951 | 3 | 0 | 3 | 5 |
| Jeffery et al. (IBS) | 25 | 0.172 | 0.3036 | 3 | 0 | 3 | 5 |
| Jeffery et al. (IBS) | 26 | 0.614 | 0.6833 | 5 | 0 | 5 | 6 |
| Jeffery et al. (IBS) | 27 | 0.022 | 0.1084 | 6 | 0 | 6 | 10 |
| Jeffery et al. (IBS) | 28 | 0.424 | 0.5044 | 4 | 0 | 4 | 6 |
| Jeffery et al. (IBS) | 29 | 0.062 | 0.1584 | 8 | 0 | 8 | 12 |
| Jeffery et al. (IBS) | 30 | 0.254 | 0.3651 | 4 | 0 | 4 | 7 |
| Jeffery et al. (IBS) | 31 | 0.024 | 0.1104 | 5 | 0 | 5 | 8 |
| Jeffery et al. (IBS) | 32 | 0.042 | 0.1270 | 4 | 0 | 4 | 6 |
| Jeffery et al. (IBS) | 33 | 0.018 | **0.0955** | 10 | 0 | 10 | 16 |
| Jeffery et al. (IBS) | 34 | 0.174 | 0.3036 | 3 | 0 | 3 | 4 |
| Jeffery et al. (IBS) | 35 | 0.858 | 0.8706 | 5 | 0 | 5 | 9 |
| Jeffery et al. (IBS) | 36 | 0.272 | 0.3809 | 3 | 0 | 3 | 5 |
| Jeffery et al. (IBS) | 37 | 0.908 | 0.9080 | 7 | 0 | 7 | 11 |
| Jeffery et al. (IBS) | 38 | 0.006 | **0.0518** | 9 | 0 | 9 | 15 |
| Jeffery et al. (IBS) | 39 | 0.276 | 0.3809 | 3 | 0 | 3 | 5 |
| Jeffery et al. (IBS) | 40 | 0.014 | **0.0878** | 3 | 0 | 3 | 5 |
| Jeffery et al. (IBS) | 41 | 0.16 | 0.3036 | 3 | 0 | 3 | 4 |
| Jeffery et al. (IBS) | 42 | 0.248 | 0.3651 | 3 | 0 | 3 | 5 |
| Jeffery et al. (IBS) | 43 | 0.328 | 0.4270 | 4 | 0 | 4 | 6 |
| Jeffery et al. (IBS) | 44 | 0.104 | 0.2175 | 4 | 0 | 4 | 7 |
| Jeffery et al. (IBS) | 45 | 0.398 | 0.4904 | 4 | 0 | 4 | 7 |
| Jeffery et al. (IBS) | 46 | 0.778 | 0.8259 | 5 | 0 | 5 | 7 |
| Jeffery et al. (IBS) | 47 | 0.002 | **0.0345** | 3 | 0 | 3 | 5 |
| Jeffery et al. (IBS) | 48 | 0.014 | **0.0878** | 3 | 0 | 3 | 5 |
| Jeffery et al. (IBS) | 49 | 0.306 | 0.4140 | 5 | 0 | 5 | 7 |
| Jeffery et al. (IBS) | 50 | 0.176 | 0.3036 | 7 | 0 | 7 | 11 |
| Jeffery et al. (IBS) | 51 | 0.016 | **0.0920** | 4 | 0 | 4 | 7 |
| Jeffery et al. (IBS) | 52 | 0.174 | 0.3036 | 3 | 0 | 3 | 5 |
| Jeffery et al. (IBS) | 53 | 0.004 | **0.0460** | 18 | 0 | 18 | 25 |
| Jeffery et al. (IBS) | 54 | 0.368 | 0.4702 | 4 | 0 | 4 | 6 |
| Jeffery et al. (IBS) | 55 | 0.03 | 0.1218 | 7 | 0 | 7 | 9 |
| Jeffery et al. (IBS) | 56 | 0.144 | 0.2839 | 3 | 0 | 3 | 5 |
| Jeffery et al. (IBS) | 57 | 0.67 | 0.7223 | 6 | 0 | 6 | 9 |
| Jeffery et al. (IBS) | 58 | 0.204 | 0.3199 | 3 | 0 | 3 | 5 |
| Jeffery et al. (IBS) | 59 | 0.006 | **0.0518** | 4 | 0 | 4 | 5 |
| Jeffery et al. (IBS) | 60 | 0.002 | **0.0345** | 4 | 0 | 4 | 8 |
| Jeffery et al. (IBS) | 61 | 0.204 | 0.3199 | 5 | 0 | 5 | 7 |
| Jeffery et al. (IBS) | 62 | 0.08 | 0.1951 | 3 | 0 | 3 | 5 |
| Jeffery et al. (IBS) | 63 | 0.044 | 0.1270 | 4 | 0 | 4 | 6 |
| Jeffery et al. (IBS) | 64 | 0.44 | 0.5146 | 6 | 0 | 6 | 10 |
| Jeffery et al. (IBS) | 65 | 0.114 | 0.2314 | 3 | 0 | 3 | 5 |
| Jeffery et al. (IBS) | 66 | 0.034 | 0.1235 | 3 | 0 | 3 | 4 |
| Jeffery et al. (IBS) | 67 | 0.19 | 0.3187 | 3 | 0 | 3 | 6 |
| Jeffery et al. (IBS) | 68 | 0.004 | **0.0460** | 3 | 0 | 3 | 5 |
| Jeffery et al. (IBS) | 69 | 0.026 | 0.1121 | 3 | 0 | 3 | 5 |
| Wang et al. (ESRD) | 1 | 0.154 | 0.1953 | 3 | 2 | 1 | 4 |
| Wang et al. (ESRD) | 2 | 0.516 | 0.5160 | 3 | 1 | 2 | 4 |
| Wang et al. (ESRD) | 3 | 0.128 | 0.1707 | 7 | 1 | 6 | 10 |
| Wang et al. (ESRD) | 4 | 0.38 | 0.3875 | 7 | 1 | 6 | 10 |
| Wang et al. (ESRD) | 5 | 0.122 | 0.1669 | 5 | 1 | 4 | 8 |
| Wang et al. (ESRD) | 6 | 0.004 | **0.0130** | 4 | 0 | 4 | 6 |
| Wang et al. (ESRD) | 7 | 0.002 | **0.0087** | 5 | 0 | 5 | 8 |
| Wang et al. (ESRD) | 8 | 0.036 | **0.0693** | 4 | 0 | 4 | 6 |
| Wang et al. (ESRD) | 9 | 0.004 | **0.0130** | 5 | 0 | 5 | 8 |
| Wang et al. (ESRD) | 10 | 0.002 | **0.0087** | 4 | 0 | 4 | 6 |
| Wang et al. (ESRD) | 11 | 0.012 | **0.0328** | 5 | 0 | 5 | 8 |
| Wang et al. (ESRD) | 12 | 0.002 | **0.0087** | 6 | 0 | 6 | 9 |
| Wang et al. (ESRD) | 13 | 0.25 | 0.2674 | 3 | 0 | 3 | 5 |
| Wang et al. (ESRD) | 14 | 0.004 | **0.0130** | 4 | 0 | 4 | 6 |
| Wang et al. (ESRD) | 15 | 0.018 | **0.0446** | 7 | 0 | 7 | 10 |
| Wang et al. (ESRD) | 16 | 0.032 | **0.0640** | 4 | 0 | 4 | 6 |
| Wang et al. (ESRD) | 17 | 0.024 | **0.0567** | 9 | 0 | 9 | 15 |
| Wang et al. (ESRD) | 18 | 0.11 | 0.1546 | 4 | 0 | 4 | 7 |
| Wang et al. (ESRD) | 19 | 0.002 | **0.0087** | 7 | 0 | 7 | 10 |

| | | | | | | | |
|---|---|---|---|---|---|---|---|
| Wang et al. (ESRD) | 20 | 0.188 | 0.2328 | 14 | 0 | 14 | 22 |
| Wang et al. (ESRD) | 21 | 0.058 | 0.1005 | 9 | 0 | 9 | 14 |
| Wang et al. (ESRD) | 22 | 0.076 | 0.1156 | 3 | 0 | 3 | 5 |
| Wang et al. (ESRD) | 23 | 0.216 | 0.2553 | 3 | 0 | 3 | 5 |
| Wang et al. (ESRD) | 24 | 0.04 | **0.0717** | 3 | 0 | 3 | 4 |
| Wang et al. (ESRD) | 25 | 0.264 | 0.2746 | 5 | 0 | 5 | 7 |
| Wang et al. (ESRD) | 26 | 0.002 | **0.0087** | 5 | 0 | 5 | 8 |
| Wang et al. (ESRD) | 27 | 0.004 | **0.0130** | 6 | 0 | 6 | 8 |
| Wang et al. (ESRD) | 28 | 0.07 | 0.1135 | 7 | 0 | 7 | 11 |
| Wang et al. (ESRD) | 29 | 0.002 | **0.0087** | 7 | 0 | 7 | 9 |
| Wang et al. (ESRD) | 30 | 0.002 | **0.0087** | 7 | 0 | 7 | 11 |
| Wang et al. (ESRD) | 31 | 0.212 | 0.2553 | 3 | 0 | 3 | 4 |
| Wang et al. (ESRD) | 32 | 0.142 | 0.1846 | 9 | 0 | 9 | 13 |
| Wang et al. (ESRD) | 33 | 0.23 | 0.2658 | 3 | 0 | 3 | 5 |
| Wang et al. (ESRD) | 34 | 0.006 | **0.0173** | 7 | 0 | 7 | 11 |
| Wang et al. (ESRD) | 35 | 0.252 | 0.2674 | 16 | 0 | 16 | 23 |
| Wang et al. (ESRD) | 36 | 0.016 | **0.0416** | 3 | 0 | 3 | 5 |
| Wang et al. (ESRD) | 37 | 0.002 | **0.0087** | 6 | 0 | 6 | 9 |
| Wang et al. (ESRD) | 38 | 0.002 | **0.0087** | 4 | 0 | 4 | 7 |
| Wang et al. (ESRD) | 39 | 0.078 | 0.1156 | 9 | 0 | 9 | 14 |
| Wang et al. (ESRD) | 40 | 0.026 | **0.0588** | 3 | 0 | 3 | 4 |
| Wang et al. (ESRD) | 41 | 0.038 | **0.0706** | 3 | 0 | 3 | 5 |
| Wang et al. (ESRD) | 42 | 0.006 | **0.0173** | 5 | 0 | 5 | 9 |
| Wang et al. (ESRD) | 43 | 0.002 | **0.0087** | 6 | 0 | 6 | 8 |
| Wang et al. (ESRD) | 44 | 0.25 | 0.2674 | 3 | 0 | 3 | 5 |
| Wang et al. (ESRD) | 45 | 0.03 | **0.0640** | 3 | 0 | 3 | 5 |
| Wang et al. (ESRD) | 46 | 0.032 | **0.0640** | 3 | 0 | 3 | 6 |
| Wang et al. (ESRD) | 47 | 0.25 | 0.2674 | 4 | 0 | 4 | 6 |
| Wang et al. (ESRD) | 48 | 0.072 | 0.1135 | 3 | 0 | 3 | 5 |
| Wang et al. (ESRD) | 49 | 0.002 | **0.0087** | 7 | 0 | 7 | 11 |
| Wang et al. (ESRD) | 50 | 0.064 | 0.1074 | 3 | 0 | 3 | 5 |
| Wang et al. (ESRD) | 51 | 0.002 | **0.0087** | 5 | 0 | 5 | 8 |
| Wang et al. (ESRD) | 52 | 0.08 | 0.1156 | 3 | 0 | 3 | 6 |

### *Supplementary Table S6: Detailed modules per dataset*

| Dataset | Module ID | Node name | Node type | DA p-value | Is an 'anchor' node $(p < \tau)$ | Node description |
|---|---|---|---|---|---|---|
| Franzosa et al. (CD) | 1 | C00062 | Metabolite | 0.0070891 | TRUE | L-Arginine |
| | | C01035 | Metabolite | 0.0360768 | TRUE | 4-Guanidinobutanoate |
| | | C00122 | Metabolite | NA | FALSE | Fumarate |
| | | EC1.3.1.6 | EC | 0.0126986 | TRUE | fumarate reductase (NADH) |
| | | EC2.1.4.1 | EC | 0.4632806 | FALSE | glycine amidinotransferase |
| | | EC3.5.3.7 | EC | 0.0156365 | TRUE | guanidinobutyrase |
| | | EC4.3.2.1 | EC | 0.0758779 | FALSE | argininosuccinate lyase |
| | | EC6.1.1.19 | EC | 0.0181257 | TRUE | arginine--tRNA ligase |
| Franzosa et al. (CD) | 2 | C00064 | Metabolite | 0.0070931 | TRUE | L-Glutamine |
| | | C00534 | Metabolite | 0.0000044 | TRUE | Pyridoxamine |
| | | C00018 | Metabolite | NA | FALSE | Pyridoxal phosphate |
| | | C00117 | Metabolite | NA | FALSE | D-Ribose 5-phosphate |
| | | C00119 | Metabolite | NA | FALSE | 5-Phospho-alpha-D-ribose 1-diphosphate |
| | | EC2.4.2.17 | EC | 0.0000068 | TRUE | ATP phosphoribosyltransferase |
| | | EC2.7.1.15 | EC | 0.0009125 | TRUE | ribokinase |
| | | EC2.7.1.35 | EC | 0.0139983 | TRUE | pyridoxal kinase |
| | | EC2.7.6.1 | EC | 0.0022334 | TRUE | ribose-phosphate diphosphokinase |
| | | EC4.3.3.6 | EC | 0.0121749 | TRUE | pyridoxal 5'-phosphate synthase (glutamine hydrolyzing) |
| | | EC5.3.1.6 | EC | 0.007103 | TRUE | ribose-5-phosphate isomerase |
| | | EC6.1.1.18 | EC | 0.0004638 | TRUE | glutamine--tRNA ligase |
| | | EC6.3.1.2 | EC | 0.0143194 | TRUE | glutamine synthetase |
| | | EC6.3.5.5 | EC | 0.0001033 | TRUE | carbamoyl-phosphate synthase (glutamine-hydrolyzing) |
| Franzosa et al. (CD) | 3 | C00073 | Metabolite | 0.1207388 | TRUE | L-Methionine |
| | | C00491 | Metabolite | 0.0000001 | TRUE | L-Cystine |
| | | C02989 | Metabolite | 0.0524836 | TRUE | L-Methionine S-oxide |
| | | C00155 | Metabolite | NA | FALSE | L-Homocysteine |
| | | EC1.8.4.14 | EC | 0.2782024 | FALSE | L-methionine (R)-S-oxide reductase |
| | | EC2.5.1.49 | EC | 0.0112205 | TRUE | O-acetylhomoserine aminocarboxypropyltransferase |
| | | EC4.4.1.13 | EC | 0.0459387 | FALSE | cysteine-S-conjugate beta-lyase |
| Franzosa et al. (CD) | 5 | C00123 | Metabolite | 0.0083732 | TRUE | L-Leucine |
| | | C00141 | Metabolite | 0.1351861 | TRUE | 3-Methyl-2-oxobutanoic acid |
| | | C00183 | Metabolite | 0.0080825 | TRUE | L-Valine |
| | | C00407 | Metabolite | 0.171816 | TRUE | L-Isoleucine |
| | | C00109 | Metabolite | NA | FALSE | 2-Oxobutanoate |
| | | EC1.1.1.85 | EC | 0.0011663 | TRUE | 3-isopropylmalate dehydrogenase |
| | | EC2.6.1.42 | EC | 0.783 | FALSE | branched-chain-amino-acid transaminase |
| Franzosa et al. (CD) | 7 | C00147 | Metabolite | 0.0001013 | TRUE | Adenine |
| | | C00242 | Metabolite | 0.0052261 | TRUE | Guanine |
| | | C00253 | Metabolite | 0.0000555 | TRUE | Nicotinate |
| | | C00299 | Metabolite | 0.1938766 | TRUE | Uridine |
| | | C00380 | Metabolite | 0.0260621 | TRUE | Cytosine |
| | | C00385 | Metabolite | 0.1310927 | TRUE | Xanthine |
| | | C00559 | Metabolite | 0.0053919 | TRUE | Deoxyadenosine |
| | | C05512 | Metabolite | 0.0997919 | TRUE | Deoxyinosine |
| | | C00130 | Metabolite | NA | FALSE | IMP |
| | | C00144 | Metabolite | NA | FALSE | GMP |
| | | C04677 | Metabolite | NA | FALSE | 1-(5'-Phosphoribosyl)-5-amino-4-imidazolecarboxamide |
| | | C00330 | Metabolite | NA | FALSE | Deoxyguanosine |
| | | EC1.7.1.7 | EC | 0.0233303 | TRUE | GMP reductase |
| | | EC2.1.2.3 | EC | 0.0037219 | TRUE | phosphoribosylaminoimidazolecarboxamide formyltransferase |
| | | EC2.4.2.1 | EC | 0.8752706 | FALSE | purine-nucleoside phosphorylase |
| | | EC2.4.2.2 | EC | 0.0088412 | TRUE | pyrimidine-nucleoside phosphorylase |
| | | EC2.4.2.7 | EC | 0.0129047 | TRUE | adenine phosphoribosyltransferase |
| | | EC2.4.2.8 | EC | 0.0006729 | TRUE | hypoxanthine phosphoribosyltransferase |
| | | EC2.7.1.113 | EC | 0.0036545 | TRUE | deoxyguanosine kinase |
| | | EC2.7.1.76 | EC | 0.0061342 | TRUE | deoxyadenosine kinase |
| | | EC3.1.5.1 | EC | 0.0031159 | TRUE | dGTPase |
| | | EC3.2.2.1 | EC | 0.0208245 | TRUE | purine nucleosidase |
| | | EC3.2.2.3 | EC | 0.0095846 | TRUE | uridine nucleosidase |
| | | EC3.5.4.10 | EC | 0.0004162 | TRUE | IMP cyclohydrolase |
| | | EC3.5.4.4 | EC | 0.0036506 | TRUE | adenosine deaminase |
| | | EC6.3.5.2 | EC | 0.0054071 | TRUE | GMP synthase (glutamine-hydrolyzing) |
| Franzosa et al. (CD) | 9 | C00163 | Metabolite | 0.1354209 | TRUE | Propanoate |
| | | C00186 | Metabolite | 0.0001325 | TRUE | (S)-Lactate |
| | | C05984 | Metabolite | 0.0019751 | TRUE | 2-Hydroxybutanoic acid |
| | | EC1.1.1.27 | EC | 0.1285315 | FALSE | L-lactate dehydrogenase |
| | | EC2.8.3.1 | EC | 0.1141363 | FALSE | propionate CoA-transferase |
| Franzosa et al. (CD) | 11 | C00219 | Metabolite | 0.0000921 | TRUE | Arachidonate |
| Franzosa et al. (CD) | 11 | C01595 | Metabolite | 0.1006297 | TRUE | Linoleate |
| | | C06427 | Metabolite | 0.0085016 | TRUE | (9Z,12Z,15Z)-Octadecatrienoic acid |
| | | C04317 | Metabolite | NA | FALSE | 1-Organyl-2-lyso-sn-glycero-3-phosphocholine |
| | | EC3.1.1.4 | EC | 0.5028989 | FALSE | phospholipase A2 |
| | | EC3.1.1.47 | EC | 0.0200866 | TRUE | 1-alkyl-2-acetylglycerophosphocholine esterase |
| Franzosa et al. (CD) | 12 | C00245 | Metabolite | 0.0004625 | TRUE | Taurine |
| | | C00695 | Metabolite | 0.0000955 | TRUE | Cholic acid |
| | | C01921 | Metabolite | 0.0109106 | TRUE | Glycocholate |
| | | C02528 | Metabolite | 0.000013 | TRUE | Chenodeoxycholate |
| | | C05465 | Metabolite | 0.10341 | TRUE | Taurochenodeoxycholate |
| | | EC3.5.1.24 | EC | 0.1483136 | FALSE | choloylglycine hydrolase |
| Franzosa et al. (CD) | 13 | C00255 | Metabolite | 0.0223581 | TRUE | Riboflavin |
| | | C00378 | Metabolite | 0.1602965 | TRUE | Thiamine |
| | | C00870 | Metabolite | 0.0745124 | TRUE | 4-Nitrophenol |
| | | C00016 | Metabolite | NA | FALSE | FAD |
| | | EC1.5.1.36 | EC | 0.003229 | TRUE | flavin reductase (NADH) |

| | | | | | | |
|---|---|---|---|---|---|---|
| | | EC2.7.7.2 | EC | 0.0194212 | TRUE | FAD synthase |
| | | EC3.1.3.1 | EC | 0.0021371 | TRUE | alkaline phosphatase |
| | | EC3.1.3.2 | EC | 0.073548 | FALSE | acid phosphatase |
| Franzosa et al. (CD) | 15 | C00300 | Metabolite | 0.0133647 | TRUE | Creatine |
| | | C00719 | Metabolite | 0.0956604 | TRUE | Betaine |
| | | C00213 | Metabolite | NA | FALSE | Sarcosine |
| | | EC2.1.1.157 | EC | 0.0063813 | TRUE | sarcosine/dimethylglycine N-methyltransferase |
| | | EC3.5.3.3 | EC | NA | FALSE | creatinase |
| Franzosa et al. (CD) | 17 | C05607 | Metabolite | 0.0074119 | TRUE | Phenyllactate |
| | | C00166 | Metabolite | NA | FALSE | Phenylpyruvate |
| | | C01179 | Metabolite | NA | FALSE | 3-(4-Hydroxyphenyl)pyruvate |
| | | C00133 | Metabolite | NA | FALSE | D-Alanine |
| | | EC1.13.11.27 | EC | 0.0101889 | TRUE | 4-hydroxyphenylpyruvate dioxygenase |
| | | EC1.3.1.12 | EC | 0.0121343 | TRUE | prephenate dehydrogenase |
| | | EC2.6.1.1 | EC | 0.0108993 | TRUE | aspartate transaminase |
| | | EC2.6.1.21 | EC | 0.0141202 | TRUE | D-amino-acid transaminase |
| | | EC4.2.1.51 | EC | 0.0058454 | TRUE | prephenate dehydratase |
| | | EC6.1.1.13 | EC | 0.0052979 | TRUE | D-alanine--poly(phosphoribitol) ligase |
| | | EC1.1.1.110 | EC | NA | FALSE | aromatic 2-oxoacid reductase |
| Franzosa et al. (CD) | 21 | C00063 | Metabolite | NA | FALSE | CTP |
| | | C00143 | Metabolite | NA | FALSE | 5,10-Methylenetetrahydrofolate |
| | | C00415 | Metabolite | NA | FALSE | Dihydrofolate |
| | | C00364 | Metabolite | NA | FALSE | dTMP |
| | | C00103 | Metabolite | NA | FALSE | D-Glucose 1-phosphate |
| | | C00857 | Metabolite | NA | FALSE | Deamino-NAD+ |
| | | EC1.17.4.2 | EC | 0.0107003 | TRUE | ribonucleoside-triphosphate reductase (thioredoxin) |
| | | EC1.5.1.3 | EC | 0.0000004 | TRUE | dihydrofolate reductase |
| | | EC1.5.1.5 | EC | 0.0176342 | TRUE | methylenetetrahydrofolate dehydrogenase (NADP(+)) |
| | | EC2.1.1.148 | EC | 0.0007374 | TRUE | thymidylate synthase (FAD) |
| | | EC2.1.1.45 | EC | 0.0000025 | TRUE | thymidylate synthase |
| | | EC2.4.1.20 | EC | 0.0000072 | TRUE | cellobiose phosphorylase |
| | | EC2.4.1.7 | EC | 0.0069104 | TRUE | sucrose phosphorylase |
| | | EC2.7.7.18 | EC | 0.0001487 | TRUE | nicotinate-nucleotide adenylyltransferase |
| | | EC2.7.7.40 | EC | 0.0180712 | TRUE | D-ribitol-5-phosphate cytidylyltransferase |
| | | EC2.7.7.9 | EC | 0.0071367 | TRUE | UTP--glucose-1-phosphate uridylyltransferase |
| | | EC3.6.1.9 | EC | 0.0449158 | FALSE | nucleotide diphosphatase |
| | | EC6.3.5.1 | EC | 0.0158834 | TRUE | NAD(+) synthase (glutamine-hydrolyzing) |
| Franzosa et al. (CD) | 22 | C00085 | Metabolite | NA | FALSE | D-Fructose 6-phosphate |
| | | C00092 | Metabolite | NA | FALSE | D-Glucose 6-phosphate |
| | | C00275 | Metabolite | NA | FALSE | D-Mannose 6-phosphate |
| | | EC2.2.1.2 | EC | 0.0194758 | TRUE | transaldolase |
| | | EC2.7.1.191 | EC | 0.0032159 | TRUE | protein-N(pi)-phosphohistidine--D-mannose phosphotransferase |
| | | EC3.2.1.86 | EC | 0.0069758 | TRUE | 6-phospho-beta-glucosidase |
| | | EC3.5.99.6 | EC | 0.0130802 | TRUE | glucosamine-6-phosphate deaminase |
| | | EC5.3.1.8 | EC | 0.008899 | TRUE | mannose-6-phosphate isomerase |
| | | EC5.4.2.6 | EC | 0.0055886 | TRUE | beta-phosphoglucomutase |
| | | EC2.7.1.1 | EC | NA | FALSE | hexokinase |
| Franzosa et al. (CD) | 26 | C00357 | Metabolite | NA | FALSE | N-Acetyl-D-glucosamine 6-phosphate |
| | | EC3.5.1.25 | EC | 0.0011891 | TRUE | N-acetylglucosamine-6-phosphate deacetylase |
| | | EC4.2.1.126 | EC | 0.0005033 | TRUE | N-acetylmuramic acid 6-phosphate etherase |
| | | EC5.1.3.9 | EC | 0.0005362 | TRUE | N-acylglucosamine-6-phosphate 2-epimerase |
| Franzosa et al. (UC) | 1 | C00047 | Metabolite | 0.0487764 | TRUE | L-Lysine |
| | | C00740 | Metabolite | NA | FALSE | D-Serine |
| | | C00793 | Metabolite | NA | FALSE | D-Cysteine |
| | | EC4.1.1.18 | EC | 0.0200206 | TRUE | lysine decarboxylase |
| | | EC4.3.1.18 | EC | 0.0023189 | TRUE | D-serine ammonia-lyase |
| | | EC4.4.1.15 | EC | 0.0010792 | TRUE | D-cysteine desulfhydrase |
| | | EC6.1.1.6 | EC | 0.0411433 | TRUE | lysine--tRNA ligase |
| | | EC5.1.1.10 | EC | NA | FALSE | amino-acid racemase |
| Franzosa et al. (UC) | 2 | C00073 | Metabolite | 0.0456204 | TRUE | L-Methionine |
| | | C02989 | Metabolite | 0.039172 | TRUE | L-Methionine S-oxide |
| | | C01179 | Metabolite | NA | FALSE | 3-(4-Hydroxyphenyl)pyruvate |
| | | C00826 | Metabolite | NA | FALSE | L-Arogenate |
| | | C05703 | Metabolite | NA | FALSE | Methaneselenol |
| | | C00155 | Metabolite | NA | FALSE | L-Homocysteine |
| | | EC1.13.11.27 | EC | 0.0056707 | TRUE | 4-hydroxyphenylpyruvate dioxygenase |
| | | EC1.3.1.43 | EC | 0.013479 | TRUE | arogenate dehydrogenase |
| | | EC1.8.1.9 | EC | 0.0873579 | TRUE | thioredoxin-disulfide reductase |
| | | EC1.8.4.14 | EC | 0.015776 | TRUE | L-methionine (R)-S-oxide reductase |
| | | EC2.1.1.14 | EC | 0.0819209 | TRUE | 5-methyltetrahydropteroyltriglutamate--homocysteine S-methyltransferase |
| | | EC2.5.1.49 | EC | 0.0000364 | TRUE | O-acetylhomoserine aminocarboxypropyltransferase |
| | | EC2.5.1.6 | EC | 0.0320355 | TRUE | methionine adenosyltransferase |
| | | EC2.6.1.1 | EC | 0.0806571 | TRUE | aspartate transaminase |
| | | EC2.6.1.57 | EC | 0.0082221 | TRUE | aromatic-amino-acid transaminase |
| | | EC2.6.1.88 | EC | 0.0038696 | TRUE | methionine transaminase |
| | | EC2.6.1.9 | EC | 0.0537561 | TRUE | histidinol-phosphate transaminase |
| | | EC4.4.1.11 | EC | 0.0232202 | TRUE | methionine gamma-lyase |
| | | EC4.4.1.13 | EC | 0.0189139 | TRUE | cysteine-S-conjugate beta-lyase |
| | | EC4.4.1.21 | EC | 0.0767176 | TRUE | S-ribosylhomocysteine lyase |
| | | EC6.1.1.10 | EC | 0.0068558 | TRUE | methionine--tRNA ligase |
| Franzosa et al. (UC) | 5 | C00163 | Metabolite | 0.0018882 | TRUE | Propanoate |
| | | C00186 | Metabolite | 0.0299332 | TRUE | (S)-Lactate |
| | | EC1.1.1.27 | EC | 0.072757 | TRUE | L-lactate dehydrogenase |
| | | EC1.1.2.3 | EC | 0.02873 | TRUE | L-lactate dehydrogenase (cytochrome) |
| | | EC1.2.1.22 | EC | 0.015001 | TRUE | lactaldehyde dehydrogenase |
| | | EC2.7.2.1 | EC | 0.0185007 | TRUE | acetate kinase |
| | | EC2.7.2.15 | EC | 0.0110678 | TRUE | propionate kinase |
| | | EC2.8.3.1 | EC | 0.0005889 | TRUE | propionate CoA-transferase |
| | | EC6.2.1.1 | EC | 0.0220136 | TRUE | acetate--CoA ligase |
| | | EC6.2.1.13 | EC | 0.0046118 | TRUE | acetate--CoA ligase (ADP-forming) |

| | | | | | | |
|---|---|---|---|---|---|---|
| | | EC6.2.1.17 | EC | 0.0167088 | TRUE | propionate--CoA ligase |
| Franzosa et al. (UC) | 6 | C00106 | Metabolite | 0.1467285 | FALSE | Uracil |
| | | C00178 | Metabolite | 0.0228594 | TRUE | Thymine |
| | | C00262 | Metabolite | 0.0372418 | TRUE | Hypoxanthine |
| | | C00385 | Metabolite | 0.048851 | TRUE | Xanthine |
| | | C00387 | Metabolite | 0.0819116 | FALSE | Guanosine |
| | | C00559 | Metabolite | 0.0036766 | TRUE | Deoxyadenosine |
| | | C05512 | Metabolite | 0.0083269 | TRUE | Deoxyinosine |
| | | C00130 | Metabolite | NA | FALSE | IMP |
| | | C00620 | Metabolite | NA | FALSE | alpha-D-Ribose 1-phosphate |
| | | C00330 | Metabolite | NA | FALSE | Deoxyguanosine |
| | | C00475 | Metabolite | NA | FALSE | Cytidine |
| | | C11736 | Metabolite | NA | FALSE | 5-Fluorodeoxyuridine |
| | | EC1.14.99.46 | EC | 0.0137342 | TRUE | pyrimidine monooxygenase |
| | | EC1.17.1.4 | EC | 0.0444586 | TRUE | xanthine dehydrogenase |
| | | EC1.7.1.7 | EC | 0.0243602 | TRUE | GMP reductase |
| | | EC2.4.2.15 | EC | 0.0246241 | TRUE | guanosine phosphorylase |
| | | EC2.4.2.2 | EC | 0.0030429 | TRUE | pyrimidine-nucleoside phosphorylase |
| | | EC2.4.2.4 | EC | 0.0053371 | TRUE | thymidine phosphorylase |
| | | EC2.7.1.21 | EC | 0.0412729 | TRUE | thymidine kinase |
| | | EC2.7.1.48 | EC | 0.0255492 | TRUE | uridine/cytidine kinase |
| | | EC2.7.1.73 | EC | 0.0070133 | TRUE | inosine kinase |
| | | EC3.1.3.6 | EC | 0.0154348 | TRUE | 3'-nucleotidase |
| | | EC3.1.3.89 | EC | 0.0054584 | TRUE | 5'-deoxynucleotidase |
| | | EC3.1.5.1 | EC | 0.0131713 | TRUE | dGTPase |
| | | EC3.2.2.8 | EC | 0.0033516 | TRUE | ribosylpyrimidine nucleosidase |
| | | EC3.5.4.1 | EC | 0.0020373 | TRUE | cytosine deaminase |
| | | EC3.5.4.10 | EC | 0.0757455 | TRUE | IMP cyclohydrolase |
| | | EC3.5.4.2 | EC | 0.0102749 | TRUE | adenine deaminase |
| | | EC3.6.1.66 | EC | 0.0074313 | TRUE | XTP/dITP diphosphatase |
| | | EC4.2.1.70 | EC | 0.010587 | TRUE | pseudouridylate synthase |
| Franzosa et al. (UC) | 8 | C00219 | Metabolite | 0.0009147 | TRUE | Arachidonate |
| | | C01595 | Metabolite | 0.0131372 | TRUE | Linoleate |
| | | C00350 | Metabolite | NA | FALSE | Phosphatidylethanolamine |
| | | EC2.3.1.40 | EC | 0.0132239 | TRUE | acyl-[acyl-carrier-protein]--phospholipid O-acyltransferase |
| | | EC3.1.1.32 | EC | 0.0042317 | TRUE | phospholipase A1 |
| | | EC3.1.1.4 | EC | 0.0112875 | TRUE | phospholipase A2 |
| | | EC4.1.1.65 | EC | 0.0027152 | TRUE | phosphatidylserine decarboxylase |
| Franzosa et al. (UC) | 14 | C06423 | Metabolite | 0.005357 | TRUE | Octanoic acid |
| | | C15972 | Metabolite | NA | FALSE | Enzyme N6-(lipoyl)lysine |
| | | C02051 | Metabolite | NA | FALSE | Lipoylprotein |
| | | C15973 | Metabolite | NA | FALSE | Enzyme N6-(dihydrolipoyl)lysine |
| | | C22160 | Metabolite | NA | FALSE | [Lipoyl-carrier protein E2]-N6-octanoyl-L-lysine |
| | | EC1.2.4.1 | EC | 0.0020826 | TRUE | pyruvate dehydrogenase (acetyl-transferring) |
| | | EC1.2.4.2 | EC | 0.0045247 | TRUE | oxoglutarate dehydrogenase (succinyl-transferring) |
| | | EC1.4.4.2 | EC | 0.0009302 | TRUE | glycine dehydrogenase (aminomethyl-transferring) |
| | | EC1.8.1.4 | EC | 0.000527 | TRUE | dihydrolipoyl dehydrogenase |
| | | EC2.3.1.12 | EC | 0.0037579 | TRUE | dihydrolipoyllysine-residue acetyltransferase |
| | | EC2.3.1.181 | EC | 0.0542052 | TRUE | lipoyl(octanoyl) transferase |
| | | EC2.3.1.61 | EC | 0.0063099 | TRUE | dihydrolipoyllysine-residue succinyltransferase |
| | | EC2.8.1.8 | EC | 0.0068629 | TRUE | lipoyl synthase |
| | | EC6.3.1.20 | EC | 0.0018009 | TRUE | lipoate--protein ligase |
| Franzosa et al. (UC) | 18 | C00085 | Metabolite | NA | FALSE | D-Fructose 6-phosphate |
| | | C01172 | Metabolite | NA | FALSE | beta-D-Glucose 6-phosphate |
| | | C00275 | Metabolite | NA | FALSE | D-Mannose 6-phosphate |
| | | EC1.1.1.140 | EC | 0.0085311 | TRUE | sorbitol-6-phosphate 2-dehydrogenase |
| | | EC1.1.1.17 | EC | 0.0469013 | TRUE | mannitol-1-phosphate 5-dehydrogenase |
| | | EC2.2.1.1 | EC | 0.026545 | TRUE | transketolase |
| | | EC2.2.1.2 | EC | 0.0462893 | TRUE | transaldolase |
| | | EC2.7.1.191 | EC | 0.0029294 | TRUE | protein-N(pi)-phosphohistidine--D-mannose phosphotransferase |
| | | EC2.7.1.90 | EC | 0.0691716 | TRUE | diphosphate--fructose-6-phosphate 1-phosphotransferase |
| | | EC3.2.1.86 | EC | 0.0068317 | TRUE | 6-phospho-beta-glucosidase |
| | | EC5.1.3.15 | EC | 0.012739 | TRUE | glucose-6-phosphate 1-epimerase |
| | | EC5.3.1.8 | EC | 0.0124942 | TRUE | mannose-6-phosphate isomerase |
| | | EC2.7.1.1 | EC | NA | FALSE | hexokinase |
| Franzosa et al. (UC) | 23 | C00168 | Metabolite | NA | FALSE | Hydroxypyruvate |
| | | C00160 | Metabolite | NA | FALSE | Glycolate |
| | | C01146 | Metabolite | NA | FALSE | 2-Hydroxy-3-oxopropanoate |
| | | EC1.1.1.29 | EC | 0.0447201 | TRUE | glycerate dehydrogenase |
| | | EC1.1.1.60 | EC | 0.023852 | TRUE | 2-hydroxy-3-oxopropionate reductase |
| | | EC1.1.1.79 | EC | 0.0168131 | TRUE | glyoxylate reductase (NADP(+)) |
| | | EC1.1.1.81 | EC | 0.0231383 | TRUE | hydroxypyruvate reductase |
| | | EC3.1.3.18 | EC | 0.0766582 | TRUE | phosphoglycolate phosphatase |
| | | EC3.8.1.2 | EC | 0.0857366 | TRUE | (S)-2-haloacid dehalogenase |
| | | EC3.8.1.3 | EC | 0.0023936 | TRUE | haloacetate dehalogenase |
| | | EC4.1.1.47 | EC | 0.0088281 | TRUE | tartronate-semialdehyde synthase |
| | | EC5.3.1.22 | EC | 0.0068332 | TRUE | hydroxypyruvate isomerase |
| Franzosa et al. (UC) | 24 | C00099 | Metabolite | 0.1843589 | FALSE | beta-Alanine |
| | | C00222 | Metabolite | NA | FALSE | 3-Oxopropanoate |
| | | C00080 | Metabolite | NA | FALSE | H+ |
| | | C12455 | Metabolite | NA | FALSE | 5-Aminopentanal |
| | | C00334 | Metabolite | NA | FALSE | 4-Aminobutanoate |
| | | EC1.1.1.298 | EC | 0.0304935 | TRUE | 3-hydroxypropionate dehydrogenase (NADP(+)) |
| | | EC1.17.1.9 | EC | 0.0047978 | TRUE | formate dehydrogenase |
| | | EC1.2.1.19 | EC | 0.0016446 | TRUE | aminobutyraldehyde dehydrogenase |
| | | EC1.2.1.27 | EC | 0.0810232 | TRUE | methylmalonate-semialdehyde dehydrogenase (CoA acylating) |
| | | EC2.6.1.19 | EC | 0.0052149 | TRUE | 4-aminobutyrate--2-oxoglutarate transaminase |
| | | EC2.6.1.82 | EC | 0.0061111 | TRUE | putrescine--2-oxoglutarate transaminase |
| | | EC3.5.3.7 | EC | 0.0075329 | TRUE | guanidinobutyrase |
| | | EC4.1.1.11 | EC | 0.0387885 | TRUE | aspartate 1-decarboxylase |

| | | | | | | |
|---|---|---|---|---|---|---|
| | | EC4.1.1.15 | EC | 0.0054334 | TRUE | glutamate decarboxylase |
| | | EC4.1.2.29 | EC | 0.03134 | TRUE | 5-dehydro-2-deoxyphosphogluconate aldolase |
| Franzosa et al. (UC) | 41 | C00136 | Metabolite | NA | FALSE | Butanoyl-CoA |
| | | C05271 | Metabolite | NA | FALSE | trans-Hex-2-enoyl-CoA |
| | | C00877 | Metabolite | NA | FALSE | Crotonoyl-CoA |
| | | C00894 | Metabolite | NA | FALSE | Propenoyl-CoA |
| | | EC1.2.1.10 | EC | 0.0133554 | TRUE | acetaldehyde dehydrogenase (acetylating) |
| | | EC1.3.1.84 | EC | 0.0857759 | TRUE | acrylyl-CoA reductase (NADPH) |
| | | EC1.3.8.1 | EC | 0.0598369 | TRUE | short-chain acyl-CoA dehydrogenase |
| | | EC1.3.8.7 | EC | 0.0041613 | TRUE | medium-chain acyl-CoA dehydrogenase |
| | | EC1.3.8.8 | EC | 0.0041613 | TRUE | long-chain-acyl-CoA dehydrogenase |
| | | EC2.8.3.12 | EC | 0.0932126 | TRUE | glutaconate CoA-transferase |
| | | EC2.8.3.8 | EC | 0.0313603 | TRUE | acetate CoA-transferase |
| | | EC2.8.3.9 | EC | 0.0036001 | TRUE | butyrate--acetoacetate CoA-transferase |
| | | EC7.2.4.5 | EC | 0.0044418 | TRUE | glutaconyl-CoA decarboxylase |
| Franzosa et al. (UC) | 45 | C04171 | Metabolite | NA | FALSE | (2S,3S)-2,3-Dihydro-2,3-dihydroxybenzoate |
| | | C00885 | Metabolite | NA | FALSE | Isochorismate |
| | | C16519 | Metabolite | NA | FALSE | 2-Succinyl-5-enolpyruvyl-6-hydroxy-3-cyclohexene-1-carboxylate |
| | | EC1.3.1.28 | EC | 0.014745 | TRUE | 2,3-dihydro-2,3-dihydroxybenzoate dehydrogenase |
| | | EC2.2.1.9 | EC | 0.0441052 | TRUE | 2-succinyl-5-enolpyruvyl-6-hydroxy-3-cyclohexene-1-carboxylic-acid |
| | | EC3.3.2.1 | EC | 0.0047293 | TRUE | isochorismatase |
| | | EC4.2.99.20 | EC | 0.0154522 | TRUE | 2-succinyl-6-hydroxy-2,4-cyclohexadiene-1-carboxylate synthase |
| Franzosa et al. (UC) | 46 | C05761 | Metabolite | NA | FALSE | Tetradecanoyl-[acp] |
| | | C06025 | Metabolite | NA | FALSE | KDO2-lipid IVA |
| | | C00229 | Metabolite | NA | FALSE | Acyl-carrier protein |
| | | C00010 | Metabolite | NA | FALSE | CoA |
| | | EC1.3.1.9 | EC | 0.0921544 | TRUE | enoyl-[acyl-carrier-protein] reductase (NADH) |
| | | EC2.3.1.241 | EC | 0.0508634 | TRUE | Kdo2-lipid IVA acyltransferase |
| | | EC2.3.1.242 | EC | 0.0057361 | TRUE | Kdo2-lipid IVA palmitoleoyltransferase |
| | | EC2.3.1.243 | EC | 0.0245015 | TRUE | acyl-Kdo2-lipid IVA acyltransferase |
| | | EC2.7.1.24 | EC | 0.0553571 | TRUE | dephospho-CoA kinase |
| | | EC2.7.8.7 | EC | 0.0119738 | TRUE | holo-[acyl-carrier-protein] synthase |
| | | EC3.1.4.14 | EC | 0.0057125 | TRUE | [acyl-carrier-protein] phosphodiesterase |
| | | EC6.2.1.20 | EC | 0.0320389 | TRUE | long-chain-fatty-acid--[acyl-carrier-protein] ligase |
| Franzosa et al. (UC) | 48 | C00143 | Metabolite | NA | FALSE | 5,10-Methylenetetrahydrofolate |
| | | C00445 | Metabolite | NA | FALSE | 5,10-Methenyltetrahydrofolate |
| | | C00365 | Metabolite | NA | FALSE | dUMP |
| | | EC1.5.1.5 | EC | 0.0260344 | TRUE | methylenetetrahydrofolate dehydrogenase (NADP(+)) |
| | | EC2.1.1.148 | EC | 0.0103823 | TRUE | thymidylate synthase (FAD) |
| | | EC2.1.1.45 | EC | 0.0478156 | TRUE | thymidylate synthase |
| | | EC2.1.2.10 | EC | 0.0023197 | TRUE | aminomethyltransferase |
| | | EC3.5.4.12 | EC | 0.0913597 | TRUE | dCMP deaminase |
| | | EC3.5.4.9 | EC | 0.0825722 | TRUE | methenyltetrahydrofolate cyclohydrolase |
| | | EC6.3.3.2 | EC | 0.0021834 | TRUE | 5-formyltetrahydrofolate cyclo-ligase |
| Franzosa et al. (UC) | 61 | C00288 | Metabolite | NA | FALSE | HCO3- |
| | | C01563 | Metabolite | NA | FALSE | Carbamate |
| | | EC2.7.2.2 | EC | 0.0221094 | TRUE | carbamate kinase |
| | | EC3.5.1.110 | EC | 0.0084051 | TRUE | ureidoacrylate amidohydrolase |
| | | EC4.2.1.1 | EC | 0.0310636 | TRUE | carbonic anhydrase |
| | | EC6.4.1.2 | EC | 0.0137271 | TRUE | acetyl-CoA carboxylase |
| Jeffery et al. (IBS) | 2 | C00047 | Metabolite | 0.00455 | TRUE | L-Lysine |
| | | C00062 | Metabolite | 0.00239 | TRUE | L-Arginine |
| | | C00064 | Metabolite | 0.022 | TRUE | L-Glutamine |
| | | C00065 | Metabolite | 0.0598 | TRUE | L-Serine |
| | | C00078 | Metabolite | 0.0197 | TRUE | L-Tryptophan |
| | | C00680 | Metabolite | NA | FALSE | meso-2,6-Diaminoheptanedioate |
| | | EC1.4.1.13 | EC | 0.0043725 | TRUE | glutamate synthase (NADPH) |
| | | EC1.4.1.16 | EC | 0.0315269 | TRUE | diaminopimelate dehydrogenase |
| | | EC2.1.2.1 | EC | 0.1012129 | TRUE | glycine hydroxymethyltransferase |
| | | EC2.4.2.14 | EC | 0.0091258 | TRUE | amidophosphoribosyltransferase |
| | | EC2.6.1.16 | EC | 0.0550353 | TRUE | glutamine--fructose-6-phosphate transaminase (isomerizing) |
| | | EC3.1.3.3 | EC | 0.0246651 | TRUE | phosphoserine phosphatase |
| | | EC3.5.3.1 | EC | 0.0177749 | TRUE | arginase |
| | | EC4.1.1.19 | EC | 0.1049725 | TRUE | arginine decarboxylase |
| | | EC4.1.1.20 | EC | 0.0008834 | TRUE | diaminopimelate decarboxylase |
| | | EC4.1.99.1 | EC | 0.0679459 | TRUE | tryptophanase |
| | | EC4.2.1.20 | EC | 0.0019985 | TRUE | tryptophan synthase |
| | | EC4.3.1.17 | EC | 0.0407892 | TRUE | L-serine ammonia-lyase |
| | | EC4.3.1.19 | EC | 0.0083417 | TRUE | threonine ammonia-lyase |
| | | EC4.3.2.1 | EC | 0.0394089 | TRUE | argininosuccinate lyase |
| | | EC5.1.1.7 | EC | 0.0007209 | TRUE | diaminopimelate epimerase |
| | | EC5.4.3.2 | EC | 0.0070809 | TRUE | lysine 2,3-aminomutase |
| | | EC6.3.1.2 | EC | 0.000353 | TRUE | glutamine synthetase |
| | | EC6.3.5.5 | EC | 0.004354 | TRUE | carbamoyl-phosphate synthase (glutamine-hydrolyzing) |
| | | EC5.1.1.10 | EC | NA | FALSE | amino-acid racemase |
| Jeffery et al. (IBS) | 4 | C00147 | Metabolite | 0.00362 | TRUE | Adenine |
| | | C00153 | Metabolite | 0.0436 | TRUE | Nicotinamide |
| | | C00212 | Metabolite | 0 | TRUE | Adenosine |
| | | C00385 | Metabolite | 0.992 | FALSE | Xanthine |
| | | C00003 | Metabolite | NA | FALSE | NAD+ |
| | | C00455 | Metabolite | NA | FALSE | Nicotinamide D-ribonucleotide |
| | | EC1.17.1.4 | EC | 0.1086373 | TRUE | xanthine dehydrogenase |
| | | EC2.4.2.1 | EC | 0.1863915 | FALSE | purine-nucleoside phosphorylase |
| | | EC2.7.1.22 | EC | 0.0057711 | TRUE | ribosylnicotinamide kinase |
| | | EC2.7.7.1 | EC | 0.1142293 | TRUE | nicotinamide-nucleotide adenylyltransferase |
| | | EC2.7.7.18 | EC | 0.0268878 | TRUE | nicotinate-nucleotide adenylyltransferase |
| | | EC3.2.2.1 | EC | 0.0009868 | TRUE | purine nucleosidase |
| | | EC3.2.2.4 | EC | 0.04242 | TRUE | AMP nucleosidase |
| | | EC3.5.1.19 | EC | 0.024392 | TRUE | nicotinamidase |
| | | EC3.5.4.2 | EC | 0.0160879 | TRUE | adenine deaminase |

| | | | | | | |
|---|---|---|---|---|---|---|
| | | EC3.5.4.3 | EC | 0.0003737 | TRUE | guanine deaminase |
| | | EC3.6.1.22 | EC | 0.0080601 | TRUE | NAD(+) diphosphatase |
| | | EC3.2.2.5 | EC | NA | FALSE | NAD(+) glycohydrolase |
| Jeffery et al. (IBS) | 16 | C01144 | Metabolite | NA | FALSE | (S)-3-Hydroxybutanoyl-CoA |
| | | C00332 | Metabolite | NA | FALSE | Acetoacetyl-CoA |
| | | C00136 | Metabolite | NA | FALSE | Butanoyl-CoA |
| | | C00246 | Metabolite | NA | FALSE | Butanoic acid |
| | | EC1.1.1.157 | EC | 0.0603314 | TRUE | 3-hydroxybutyryl-CoA dehydrogenase |
| | | EC1.1.1.36 | EC | 0.0707496 | TRUE | acetoacetyl-CoA reductase |
| | | EC1.3.8.1 | EC | 0.0639693 | TRUE | short-chain acyl-CoA dehydrogenase |
| | | EC2.7.2.7 | EC | 0.0583767 | TRUE | butyrate kinase |
| | | EC2.8.3.8 | EC | 0.0093017 | TRUE | acetate CoA-transferase |
| | | EC2.8.3.9 | EC | 0.0100055 | TRUE | butyrate--acetoacetate CoA-transferase |
| | | EC5.1.2.3 | EC | 0.1233479 | TRUE | 3-hydroxybutyryl-CoA epimerase |
| | | EC6.2.1.16 | EC | 0.0463745 | TRUE | acetoacetate--CoA ligase |
| Jeffery et al. (IBS) | 33 | C00106 | Metabolite | 0.411 | FALSE | Uracil |
| | | C00299 | Metabolite | 0.736 | FALSE | Uridine |
| | | C21028 | Metabolite | NA | FALSE | (R)-5,6-Dihydrothymine |
| | | C00365 | Metabolite | NA | FALSE | dUMP |
| | | C00526 | Metabolite | NA | FALSE | Deoxyuridine |
| | | C00105 | Metabolite | NA | FALSE | UMP |
| | | EC1.3.1.1 | EC | 0.0459956 | TRUE | dihydrouracil dehydrogenase (NAD(+)) |
| | | EC2.1.1.148 | EC | 0.1097752 | TRUE | thymidylate synthase (FAD) |
| | | EC2.4.2.3 | EC | 0.0075441 | TRUE | uridine phosphorylase |
| | | EC2.4.2.9 | EC | 0.1003691 | TRUE | uracil phosphoribosyltransferase |
| | | EC2.7.1.21 | EC | 0.0771583 | TRUE | thymidine kinase |
| | | EC3.2.2.8 | EC | 0.104194 | TRUE | ribosylpyrimidine nucleosidase |
| | | EC3.5.2.2 | EC | 0.0112293 | TRUE | dihydropyrimidinase |
| | | EC3.5.4.12 | EC | 0.0486858 | TRUE | dCMP deaminase |
| | | EC3.5.4.5 | EC | 0.0234026 | TRUE | cytidine deaminase |
| | | EC4.1.1.23 | EC | 0.0209007 | TRUE | orotidine-5'-phosphate decarboxylase |
| Jeffery et al. (IBS) | 38 | C00255 | Metabolite | 0.0716 | FALSE | Riboflavin |
| | | C01847 | Metabolite | NA | FALSE | Reduced FMN |
| | | C00870 | Metabolite | NA | FALSE | 4-Nitrophenol |
| | | C00061 | Metabolite | NA | FALSE | FMN |
| | | C00378 | Metabolite | NA | FALSE | Thiamine |
| | | C01081 | Metabolite | NA | FALSE | Thiamin monophosphate |
| | | EC1.5.1.38 | EC | 0.1278402 | TRUE | FMN reductase (NADPH) |
| | | EC1.5.1.39 | EC | 0.0000592 | TRUE | FMN reductase [NAD(P)H] |
| | | EC2.5.1.129 | EC | 0.0442198 | TRUE | flavin prenyltransferase |
| | | EC2.5.1.3 | EC | 0.0195021 | TRUE | thiamine phosphate synthase |
| | | EC2.5.1.9 | EC | 0.1279267 | TRUE | riboflavin synthase |
| | | EC2.7.6.2 | EC | 0.0029873 | TRUE | thiamine diphosphokinase |
| | | EC3.1.3.1 | EC | 0.0076504 | TRUE | alkaline phosphatase |
| | | EC3.1.3.2 | EC | 0.0204965 | TRUE | acid phosphatase |
| | | EC3.1.8.1 | EC | 0.0035917 | TRUE | aryldialkylphosphatase |
| Jeffery et al. (IBS) | 40 | C02463 | Metabolite | NA | FALSE | Precorrin 2 |
| | | C01051 | Metabolite | NA | FALSE | Uroporphyrinogen III |
| | | EC2.1.1.107 | EC | 0.0165905 | TRUE | uroporphyrinogen-III C-methyltransferase |
| | | EC2.1.1.130 | EC | 0.1071368 | TRUE | precorrin-2 C(20)-methyltransferase |
| | | EC4.1.1.37 | EC | 0.027818 | TRUE | uroporphyrinogen decarboxylase |
| Jeffery et al. (IBS) | 47 | C19848 | Metabolite | NA | FALSE | Adenylated molybdopterin |
| | | C18237 | Metabolite | NA | FALSE | Molybdoenzyme molybdenum cofactor |
| | | EC2.10.1.1 | EC | 0.0095764 | TRUE | molybdopterin molybdotransferase |
| | | EC2.7.7.75 | EC | 0.0032074 | TRUE | molybdopterin adenylyltransferase |
| | | EC2.7.7.77 | EC | 0.000755 | TRUE | molybdenum cofactor guanylyltransferase |
| Jeffery et al. (IBS) | 48 | C11437 | Metabolite | NA | FALSE | 1-Deoxy-D-xylulose 5-phosphate |
| | | C15809 | Metabolite | NA | FALSE | Iminoglycine |
| | | EC2.2.1.7 | EC | 0.0680482 | TRUE | 1-deoxy-D-xylulose-5-phosphate synthase |
| | | EC2.8.1.10 | EC | 0.0507634 | TRUE | thiazole synthase |
| | | EC4.1.99.19 | EC | 0.0003403 | TRUE | 2-iminoacetate synthase |
| Jeffery et al. (IBS) | 51 | C01092 | Metabolite | NA | FALSE | 8-Amino-7-oxononanoate |
| | | C01037 | Metabolite | NA | FALSE | 7,8-Diaminononanoate |
| | | C01909 | Metabolite | NA | FALSE | Dethiobiotin |
| | | EC2.3.1.47 | EC | 0.0049038 | TRUE | 8-amino-7-oxononanoate synthase |
| | | EC2.6.1.62 | EC | 0.0001355 | TRUE | adenosylmethionine--8-amino-7-oxononanoate transaminase |
| | | EC2.8.1.6 | EC | 0.0040822 | TRUE | biotin synthase |
| | | EC6.3.3.3 | EC | 0.0036338 | TRUE | dethiobiotin synthase |
| Jeffery et al. (IBS) | 53 | C00029 | Metabolite | NA | FALSE | UDP-glucose |
| | | C00369 | Metabolite | NA | FALSE | Starch |
| | | C00103 | Metabolite | NA | FALSE | D-Glucose 1-phosphate |
| | | C00089 | Metabolite | NA | FALSE | Sucrose |
| | | C00718 | Metabolite | NA | FALSE | Amylose |
| | | C00052 | Metabolite | NA | FALSE | UDP-alpha-D-galactose |
| | | C01935 | Metabolite | NA | FALSE | Maltodextrin |
| | | EC2.4.1.1 | EC | 0.065038 | TRUE | glycogen phosphorylase |
| | | EC2.4.1.11 | EC | 0.0891718 | TRUE | glycogen(starch) synthase |
| | | EC2.4.1.18 | EC | 0.0520558 | TRUE | 1,4-alpha-glucan branching enzyme |
| | | EC2.4.1.20 | EC | 0.0561356 | TRUE | cellobiose phosphorylase |
| | | EC2.4.1.21 | EC | 0.0206606 | TRUE | starch synthase |
| | | EC2.4.1.4 | EC | 0.0084425 | TRUE | amylosucrase |
| | | EC2.4.1.5 | EC | 0.0094693 | TRUE | dextransucrase |
| | | EC2.4.1.7 | EC | 0.0537858 | TRUE | sucrose phosphorylase |
| | | EC2.4.99.16 | EC | 0.1190859 | TRUE | starch synthase (maltosyl-transferring) |
| | | EC2.7.7.27 | EC | 0.0990762 | TRUE | glucose-1-phosphate adenylyltransferase |
| | | EC2.7.7.9 | EC | 0.0412613 | TRUE | UTP--glucose-1-phosphate uridylyltransferase |
| | | EC3.2.1.1 | EC | 0.0042664 | TRUE | alpha-amylase |
| | | EC3.2.1.3 | EC | 0.106752 | TRUE | glucan 1,4-alpha-glucosidase |
| | | EC3.2.1.54 | EC | 0.0055335 | TRUE | cyclomaltodextrinase |
| | | EC3.2.1.68 | EC | 0.0819147 | TRUE | isoamylase |

| | | | | | | |
|---|---|---|---|---|---|---|
| | | EC3.6.1.45 | EC | 0.1182435 | TRUE | UDP-sugar diphosphatase |
| | | EC5.1.3.2 | EC | 0.0002219 | TRUE | UDP-glucose 4-epimerase |
| | | EC5.4.99.9 | EC | 0.0005754 | TRUE | UDP-galactopyranose mutase |
| Jeffery et al. (IBS) | 59 | C00129 | Metabolite | NA | FALSE | Isopentenyl diphosphate |
| | | EC2.5.1.1 | EC | 0.0095195 | TRUE | dimethylallyltranstransferase |
| | | EC2.5.1.10 | EC | 0.0014252 | TRUE | (2E,6E)-farnesyl diphosphate synthase |
| | | EC2.5.1.29 | EC | 0.0024533 | TRUE | geranylgeranyl diphosphate synthase |
| | | EC5.3.3.2 | EC | 0.0040147 | TRUE | isopentenyl-diphosphate Delta-isomerase |
| Jeffery et al. (IBS) | 60 | C06508 | Metabolite | NA | FALSE | Adenosyl cobinamide |
| | | C06509 | Metabolite | NA | FALSE | Adenosyl cobinamide phosphate |
| | | C04122 | Metabolite | NA | FALSE | D-1-Aminopropan-2-ol O-phosphate |
| | | EC2.5.1.17 | EC | 0.0007645 | TRUE | corrinoid adenosyltransferase |
| | | EC2.7.1.156 | EC | 0.0060653 | TRUE | adenosylcobinamide kinase |
| | | EC2.7.7.62 | EC | 0.0133206 | TRUE | adenosylcobinamide-phosphate guanylyltransferase |
| | | EC4.1.1.81 | EC | 0.0004845 | TRUE | threonine-phosphate decarboxylase |
| | | EC6.3.1.10 | EC | NA | FALSE | adenosylcobinamide-phosphate synthase |
| Jeffery et al. (IBS) | 68 | C00086 | Metabolite | NA | FALSE | Urea |
| | | C01010 | Metabolite | NA | FALSE | Urea-1-carboxylate |
| | | EC3.5.1.5 | EC | 0.0012855 | TRUE | urease |
| | | EC3.5.1.54 | EC | 0.0280354 | TRUE | allophanate hydrolase |
| | | EC6.3.4.6 | EC | 0.1367414 | TRUE | urea carboxylase |
| Wang et al. (ESRD) | 6 | C00188 | Metabolite | NA | FALSE | L-Threonine |
| | | C03508 | Metabolite | NA | FALSE | L-2-Amino-3-oxobutanoic acid |
| | | EC1.1.1.103 | EC | 0.0011409 | TRUE | L-threonine 3-dehydrogenase |
| | | EC2.3.1.29 | EC | 0.0011881 | TRUE | glycine C-acetyltransferase |
| | | EC4.1.2.48 | EC | 0.0715074 | TRUE | low-specificity L-threonine aldolase |
| | | EC4.2.3.1 | EC | 0.0017932 | TRUE | threonine synthase |
| Wang et al. (ESRD) | 7 | C00310 | Metabolite | NA | FALSE | D-Xylulose |
| | | C00181 | Metabolite | NA | FALSE | D-Xylose |
| | | C00267 | Metabolite | NA | FALSE | alpha-D-Glucose |
| | | EC1.1.1.14 | EC | 0.0812683 | TRUE | L-iditol 2-dehydrogenase |
| | | EC1.1.1.179 | EC | 0.0505813 | TRUE | D-xylose 1-dehydrogenase (NADP(+), D-xylono-1,5-lactone-forming) |
| | | EC3.2.1.23 | EC | 0.0787603 | TRUE | beta-galactosidase |
| | | EC5.1.3.3 | EC | 0.0443571 | TRUE | aldose 1-epimerase |
| | | EC5.3.1.5 | EC | 0.006861 | TRUE | xylose isomerase |
| Wang et al. (ESRD) | 8 | C00116 | Metabolite | NA | FALSE | Glycerol |
| | | C06103 | Metabolite | NA | FALSE | 6-Hydroxyhexanoic acid |
| | | EC1.1.1.2 | EC | 0.0001528 | TRUE | alcohol dehydrogenase (NADP(+)) |
| | | EC2.7.1.30 | EC | 0.0116437 | TRUE | glycerol kinase |
| | | EC3.1.1.83 | EC | 0.0294566 | TRUE | monoterpene epsilon-lactone hydrolase |
| | | EC4.2.1.30 | EC | 0.0112488 | TRUE | glycerol dehydratase |
| Wang et al. (ESRD) | 9 | C00135 | Metabolite | NA | FALSE | L-Histidine |
| | | C00785 | Metabolite | NA | FALSE | Urocanate |
| | | C03680 | Metabolite | NA | FALSE | 4-Imidazolone-5-propanoate |
| | | EC1.1.1.23 | EC | 0.0192425 | TRUE | histidinol dehydrogenase |
| | | EC3.5.2.7 | EC | 0.0012146 | TRUE | imidazolonepropionase |
| | | EC4.2.1.49 | EC | 0.0006472 | TRUE | urocanate hydratase |
| | | EC4.3.1.3 | EC | 0.0001554 | TRUE | histidine ammonia-lyase |
| | | EC6.1.1.21 | EC | 0.0818091 | TRUE | histidine--tRNA ligase |
| Wang et al. (ESRD) | 10 | C11638 | Metabolite | NA | FALSE | 3-Amino-2-oxopropyl phosphate |
| | | C00627 | Metabolite | NA | FALSE | Pyridoxine phosphate |
| | | EC1.1.1.262 | EC | 0.0534713 | TRUE | 4-hydroxythreonine-4-phosphate dehydrogenase |
| | | EC1.4.3.5 | EC | 0.003315 | TRUE | pyridoxal 5'-phosphate synthase |
| | | EC2.6.99.2 | EC | 0.0007355 | TRUE | pyridoxine 5'-phosphate synthase |
| | | EC2.7.1.35 | EC | 0.0979069 | TRUE | pyridoxal kinase |
| Wang et al. (ESRD) | 11 | C11437 | Metabolite | NA | FALSE | 1-Deoxy-D-xylulose 5-phosphate |
| | | C11434 | Metabolite | NA | FALSE | 2-C-Methyl-D-erythritol 4-phosphate |
| | | C15809 | Metabolite | NA | FALSE | Iminoglycine |
| | | EC1.1.1.267 | EC | 0.0334971 | TRUE | 1-deoxy-D-xylulose-5-phosphate reductoisomerase |
| | | EC2.2.1.7 | EC | 0.0281836 | TRUE | 1-deoxy-D-xylulose-5-phosphate synthase |
| | | EC2.7.7.60 | EC | 0.0884026 | TRUE | 2-C-methyl-D-erythritol 4-phosphate cytidylyltransferase |
| | | EC2.8.1.10 | EC | 0.0001998 | TRUE | thiazole synthase |
| | | EC4.1.99.19 | EC | 0.0262437 | TRUE | 2-iminoacetate synthase |
| Wang et al. (ESRD) | 12 | C00258 | Metabolite | NA | FALSE | D-Glycerate |
| | | C00160 | Metabolite | NA | FALSE | Glycolate |
| | | C00546 | Metabolite | NA | FALSE | Methylglyoxal |
| | | EC1.1.1.283 | EC | 0.0435836 | TRUE | methylglyoxal reductase (NADPH) |
| | | EC1.1.1.29 | EC | 0.0109704 | TRUE | glycerate dehydrogenase |
| | | EC1.1.1.79 | EC | 0.0740202 | TRUE | glyoxylate reductase (NADP(+)) |
| | | EC1.1.1.81 | EC | 0.0110843 | TRUE | hydroxypyruvate reductase |
| | | EC3.8.1.2 | EC | 0.0515842 | TRUE | (S)-2-haloacid dehalogenase |
| | | EC4.4.1.5 | EC | 0.0368793 | TRUE | lactoylglutathione lyase |
| Wang et al. (ESRD) | 14 | C11537 | Metabolite | NA | FALSE | (2R)-3-Sulfolactate |
| | | C00149 | Metabolite | NA | FALSE | (S)-Malate |
| | | EC1.1.1.37 | EC | 0.0001231 | TRUE | malate dehydrogenase |
| | | EC1.1.1.38 | EC | 0.0503857 | TRUE | malate dehydrogenase (oxaloacetate-decarboxylating) |
| | | EC1.1.1.40 | EC | 0.0000001 | TRUE | malate dehydrogenase (oxaloacetate-decarboxylating) (NADP(+)) |
| | | EC3.1.3.71 | EC | 0.0489055 | TRUE | 2-phosphosulfolactate phosphatase |
| Wang et al. (ESRD) | 15 | C00199 | Metabolite | NA | FALSE | D-Ribulose 5-phosphate |
| | | C00345 | Metabolite | NA | FALSE | 6-Phospho-D-gluconate |
| | | C00006 | Metabolite | NA | FALSE | NADP+ |
| | | EC1.1.1.42 | EC | 0.0196171 | TRUE | isocitrate dehydrogenase (NADP(+)) |
| | | EC1.1.1.44 | EC | 0.003237 | TRUE | phosphogluconate dehydrogenase (NADP(+)-dependent, decarboxylating) |
| | | EC1.1.1.49 | EC | 0.0013033 | TRUE | glucose-6-phosphate dehydrogenase (NADP(+)) |
| | | EC1.8.1.7 | EC | 0.0936285 | TRUE | glutathione-disulfide reductase |
| | | EC2.7.1.12 | EC | 0.0863611 | TRUE | gluconokinase |
| | | EC2.7.1.16 | EC | 0.0336438 | TRUE | ribulokinase |
| | | EC3.1.1.31 | EC | 0.0040088 | TRUE | 6-phosphogluconolactonase |
| Wang et al. (ESRD) | 16 | C00232 | Metabolite | NA | FALSE | Succinate semialdehyde |
| | | C00091 | Metabolite | NA | FALSE | Succinyl-CoA |

| | | | | | | |
|---|---|---|---|---|---|---|
| | | EC1.1.1.61 | EC | 0.0024204 | TRUE | 4-hydroxybutyrate dehydrogenase |
| | | EC1.2.1.76 | EC | 0.0218723 | TRUE | succinate-semialdehyde dehydrogenase (acetylating) |
| | | EC1.2.7.3 | EC | 0.0068597 | TRUE | 2-oxoglutarate synthase |
| | | EC5.4.99.2 | EC | 0.0088491 | TRUE | methylmalonyl-CoA mutase |
| Wang et al. (ESRD) | 17 | C00233 | Metabolite | NA | FALSE | 4-Methyl-2-oxopentanoate |
| | | C15972 | Metabolite | NA | FALSE | Enzyme N6-(lipoyl)lysine |
| | | C00141 | Metabolite | NA | FALSE | 3-Methyl-2-oxobutanoic acid |
| | | C15980 | Metabolite | NA | FALSE | (S)-2-Methylbutanoyl-CoA |
| | | C15973 | Metabolite | NA | FALSE | Enzyme N6-(dihydrolipoyl)lysine |
| | | C22159 | Metabolite | NA | FALSE | [Glycine cleavage system H]-N6-octanoyl-L-lysine |
| | | EC1.1.1.85 | EC | 0.0102001 | TRUE | 3-isopropylmalate dehydrogenase |
| | | EC1.2.4.4 | EC | 0.0083933 | TRUE | 3-methyl-2-oxobutanoate dehydrogenase (2-methylpropanoyl-transferring) |
| | | EC1.2.7.7 | EC | 0.0368822 | TRUE | 3-methyl-2-oxobutanoate dehydrogenase (ferredoxin) |
| | | EC1.3.8.1 | EC | 0.029919 | TRUE | short-chain acyl-CoA dehydrogenase |
| | | EC1.8.1.4 | EC | 0.0003511 | TRUE | dihydrolipoyl dehydrogenase |
| | | EC2.1.2.11 | EC | 0.0086436 | TRUE | 3-methyl-2-oxobutanoate hydroxymethyltransferase |
| | | EC2.3.1.12 | EC | 0.0833308 | TRUE | dihydrolipoyllysine-residue acetyltransferase |
| | | EC2.3.1.181 | EC | 0.0001125 | TRUE | lipoyl(octanoyl) transferase |
| | | EC2.8.1.8 | EC | 0.0005155 | TRUE | lipoyl synthase |
| Wang et al. (ESRD) | 19 | C03972 | Metabolite | NA | FALSE | 2,3,4,5-Tetrahydrodipicolinate |
| | | C04462 | Metabolite | NA | FALSE | N-Succinyl-2-L-amino-6-oxoheptanedioate |
| | | C00666 | Metabolite | NA | FALSE | LL-2,6-Diaminoheptanedioate |
| | | EC1.17.1.8 | EC | 0.0009654 | TRUE | 4-hydroxy-tetrahydrodipicolinate reductase |
| | | EC2.3.1.117 | EC | 0.0322356 | TRUE | 2,3,4,5-tetrahydropyridine-2,6-dicarboxylate N-succinyltransferase |
| | | EC2.6.1.17 | EC | 0.0295176 | TRUE | succinyldiaminopimelate transaminase |
| | | EC2.6.1.83 | EC | 0.0000008 | TRUE | LL-diaminopimelate aminotransferase |
| | | EC3.5.1.18 | EC | 0.0059892 | TRUE | succinyl-diaminopimelate desuccinylase |
| | | EC3.5.1.47 | EC | 0.0341039 | TRUE | N-acetyldiaminopimelate deacetylase |
| | | EC5.1.1.7 | EC | 0.0327022 | TRUE | diaminopimelate epimerase |
| Wang et al. (ESRD) | 24 | C00254 | Metabolite | NA | FALSE | Prephenate |
| | | EC1.3.1.12 | EC | 0.0289341 | TRUE | prephenate dehydrogenase |
| | | EC4.2.1.51 | EC | 0.0502756 | TRUE | prephenate dehydratase |
| | | EC4.2.1.91 | EC | 0.0556655 | TRUE | arogenate dehydratase |
| Wang et al. (ESRD) | 26 | C02463 | Metabolite | NA | FALSE | Precorrin 2 |
| | | C05778 | Metabolite | NA | FALSE | Sirohydrochlorin |
| | | C01051 | Metabolite | NA | FALSE | Uroporphyrinogen III |
| | | EC1.3.1.76 | EC | 0.0188292 | TRUE | precorrin-2 dehydrogenase |
| | | EC2.1.1.107 | EC | 0.060875 | TRUE | uroporphyrinogen-III C-methyltransferase |
| | | EC2.1.1.130 | EC | 0.0000032 | TRUE | precorrin-2 C(20)-methyltransferase |
| | | EC4.1.1.37 | EC | 0.064483 | TRUE | uroporphyrinogen decarboxylase |
| | | EC4.99.1.4 | EC | 0.0002904 | TRUE | sirohydrochlorin ferrochelatase |
| Wang et al. (ESRD) | 27 | C00111 | Metabolite | NA | FALSE | Glycerone phosphate |
| | | C05840 | Metabolite | NA | FALSE | Iminoaspartate |
| | | EC1.4.3.16 | EC | 0.0016007 | TRUE | L-aspartate oxidase |
| | | EC2.5.1.72 | EC | 0.0002034 | TRUE | quinolinate synthase |
| | | EC2.7.1.121 | EC | 0.0139601 | TRUE | phosphoenolpyruvate--glycerone phosphotransferase |
| | | EC2.7.1.29 | EC | 0.0736325 | TRUE | glycerone kinase |
| | | EC4.1.2.40 | EC | 0.0170603 | TRUE | tagatose-bisphosphate aldolase |
| | | EC5.3.1.1 | EC | 0 | TRUE | triose-phosphate isomerase |
| Wang et al. (ESRD) | 29 | C00064 | Metabolite | NA | FALSE | L-Glutamine |
| | | C00117 | Metabolite | NA | FALSE | D-Ribose 5-phosphate |
| | | EC1.4.7.1 | EC | 0 | TRUE | glutamate synthase (ferredoxin) |
| | | EC2.7.1.15 | EC | 0.0002112 | TRUE | ribokinase |
| | | EC2.7.6.1 | EC | 0.0008138 | TRUE | ribose-phosphate diphosphokinase |
| | | EC3.6.1.13 | EC | 0.0000009 | TRUE | ADP-ribose diphosphatase |
| | | EC4.3.3.6 | EC | 0 | TRUE | pyridoxal 5'-phosphate synthase (glutamine hydrolyzing) |
| | | EC6.1.1.18 | EC | 0.0005931 | TRUE | glutamine--tRNA ligase |
| | | EC6.3.1.2 | EC | 0.0032289 | TRUE | glutamine synthetase |
| Wang et al. (ESRD) | 30 | C00870 | Metabolite | NA | FALSE | 4-Nitrophenol |
| | | C00061 | Metabolite | NA | FALSE | FMN |
| | | C00378 | Metabolite | NA | FALSE | Thiamine |
| | | EC1.5.1.38 | EC | 0.0853902 | TRUE | FMN reductase (NADPH) |
| | | EC1.5.1.39 | EC | 0.0017566 | TRUE | FMN reductase [NAD(P)H] |
| | | EC2.7.1.26 | EC | 0.000868 | TRUE | riboflavin kinase |
| | | EC2.7.6.2 | EC | 0.0265151 | TRUE | thiamine diphosphokinase |
| | | EC2.7.7.2 | EC | 0.0008373 | TRUE | FAD synthase |
| | | EC3.1.3.1 | EC | 0.0007542 | TRUE | alkaline phosphatase |
| | | EC3.1.3.2 | EC | 0.1449444 | FALSE | acid phosphatase |
| | | EC3.1.8.1 | EC | 0.0021473 | TRUE | aryldialkylphosphatase |
| Wang et al. (ESRD) | 34 | C00062 | Metabolite | NA | FALSE | L-Arginine |
| | | C00327 | Metabolite | NA | FALSE | L-Citrulline |
| | | C00077 | Metabolite | NA | FALSE | L-Ornithine |
| | | C15532 | Metabolite | NA | FALSE | N-Acetyl-L-citrulline |
| | | EC2.1.3.9 | EC | 0.077758 | TRUE | N-acetylornithine carbamoyltransferase |
| | | EC2.6.1.13 | EC | 0.0472936 | TRUE | ornithine aminotransferase |
| | | EC3.5.1.16 | EC | 0.0110068 | TRUE | acetylornithine deacetylase |
| | | EC3.5.3.1 | EC | 0.0239319 | TRUE | arginase |
| | | EC3.5.3.6 | EC | 0.0505194 | TRUE | arginine deiminase |
| | | EC4.3.2.1 | EC | 0.0056782 | TRUE | argininosuccinate lyase |
| | | EC6.3.4.5 | EC | 0.0750683 | TRUE | argininosuccinate synthase |
| Wang et al. (ESRD) | 36 | C01063 | Metabolite | NA | FALSE | Pimeloyl-CoA |
| | | C01092 | Metabolite | NA | FALSE | 8-Amino-7-oxononanoate |
| | | EC2.3.1.47 | EC | 0.0588412 | TRUE | 8-amino-7-oxononanoate synthase |
| | | EC2.6.1.62 | EC | 0.0893766 | TRUE | adenosylmethionine--8-amino-7-oxononanoate transaminase |
| | | EC6.2.1.14 | EC | 0.0026619 | TRUE | 6-carboxyhexanoate--CoA ligase |
| Wang et al. (ESRD) | 37 | C00641 | Metabolite | NA | FALSE | 1,2-Diacyl-sn-glycerol |
| | | C00416 | Metabolite | NA | FALSE | Phosphatidate |
| | | C01177 | Metabolite | NA | FALSE | Inositol 1-phosphate |
| | | EC2.3.1.51 | EC | 0.0149035 | TRUE | 1-acylglycerol-3-phosphate O-acyltransferase |
| | | EC2.7.1.107 | EC | 0.082929 | TRUE | diacylglycerol kinase (ATP) |

| | | | | | | |
|---|---|---|---|---|---|---|
| | | EC3.1.1.3 | EC | 0.0005783 | TRUE | triacylglycerol lipase |
| | | EC3.1.3.25 | EC | 0.0000931 | TRUE | inositol-phosphate phosphatase |
| | | EC3.1.3.4 | EC | 0.0600234 | TRUE | phosphatidate phosphatase |
| | | EC4.6.1.13 | EC | 0.0027144 | TRUE | phosphatidylinositol diacylglycerol-lyase |
| Wang et al. (ESRD) | 38 | C00134 | Metabolite | NA | FALSE | Putrescine |
| | | C00315 | Metabolite | NA | FALSE | Spermidine |
| | | EC2.3.1.57 | EC | 0.0190643 | TRUE | diamine N-acetyltransferase |
| | | EC2.5.1.16 | EC | 0.2210627 | FALSE | spermidine synthase |
| | | EC3.5.1.53 | EC | 0.0318271 | TRUE | N-carbamoylputrescine amidase |
| | | EC3.5.3.11 | EC | 0.035426 | TRUE | agmatinase |
| | | EC4.1.1.96 | EC | 0.0010214 | TRUE | carboxynorspermidine decarboxylase |
| Wang et al. (ESRD) | 40 | C00140 | Metabolite | NA | FALSE | N-Acetyl-D-glucosamine |
| | | EC2.4.1.280 | EC | 0.045516 | TRUE | N,N'-diacetylchitobiose phosphorylase |
| | | EC2.7.1.162 | EC | 0.0758786 | TRUE | N-acetylhexosamine 1-kinase |
| | | EC3.2.1.14 | EC | 0.0168406 | TRUE | chitinase |
| Wang et al. (ESRD) | 41 | G13167 | Metabolite | NA | FALSE | NA |
| | | C00133 | Metabolite | NA | FALSE | D-Alanine |
| | | EC2.4.1.52 | EC | 0.0912361 | TRUE | poly(glycerol-phosphate) alpha-glucosyltransferase |
| | | EC6.1.1.13 | EC | 0.0013603 | TRUE | D-alanine--poly(phosphoribitol) ligase |
| | | EC6.3.2.4 | EC | 0.0327166 | TRUE | D-alanine--D-alanine ligase |
| Wang et al. (ESRD) | 42 | C00106 | Metabolite | NA | FALSE | Uracil |
| | | C21028 | Metabolite | NA | FALSE | (R)-5,6-Dihydrothymine |
| | | C00299 | Metabolite | NA | FALSE | Uridine |
| | | EC1.3.1.2 | EC | 0.2442189 | FALSE | dihydropyrimidine dehydrogenase (NADP(+)) |
| | | EC2.4.2.9 | EC | 0.0254023 | TRUE | uracil phosphoribosyltransferase |
| | | EC3.2.2.3 | EC | 0.0861802 | TRUE | uridine nucleosidase |
| | | EC3.5.2.2 | EC | 0.0003186 | TRUE | dihydropyrimidinase |
| | | EC3.5.4.5 | EC | 0.0000754 | TRUE | cytidine deaminase |
| | | EC4.2.1.70 | EC | 0.0763505 | TRUE | pseudouridylate synthase |
| Wang et al. (ESRD) | 43 | C00074 | Metabolite | NA | FALSE | Phosphoenolpyruvate |
| | | C04691 | Metabolite | NA | FALSE | 2-Dehydro-3-deoxy-D-arabino-heptonate 7-phosphate |
| | | EC2.5.1.54 | EC | 0.0227864 | TRUE | 3-deoxy-7-phosphoheptulonate synthase |
| | | EC2.7.9.2 | EC | 0.0337732 | TRUE | pyruvate, water dikinase |
| | | EC4.1.1.31 | EC | 0.0553958 | TRUE | phosphoenolpyruvate carboxylase |
| | | EC4.1.1.32 | EC | 0.0196531 | TRUE | phosphoenolpyruvate carboxykinase (GTP) |
| | | EC4.1.1.49 | EC | 0.0231634 | TRUE | phosphoenolpyruvate carboxykinase (ATP) |
| | | EC4.2.3.4 | EC | 0.0003653 | TRUE | 3-dehydroquinate synthase |
| Wang et al. (ESRD) | 45 | C19972 | Metabolite | NA | FALSE | 2,4-Bis(acetamido)-2,4,6-trideoxy-beta-L-altropyranose |
| | | C20082 | Metabolite | NA | FALSE | Pseudaminic acid |
| | | EC2.5.1.97 | EC | 0.0616637 | TRUE | pseudaminic acid synthase |
| | | EC2.7.7.81 | EC | 0.0374947 | TRUE | pseudaminic acid cytidylyltransferase |
| | | EC3.6.1.57 | EC | 0.0963297 | TRUE | UDP-2,4-diacetamido-2,4,6-trideoxy-beta-L-altropyranose hydrolase |
| Wang et al. (ESRD) | 46 | C04188 | Metabolite | NA | FALSE | S-Methyl-5-thio-D-ribose 1-phosphate |
| | | C19787 | Metabolite | NA | FALSE | 5'-S-Methyl-5'-thioinosine |
| | | EC2.7.1.100 | EC | 0.0004615 | TRUE | S-methyl-5-thioribose kinase |
| | | EC3.5.4.31 | EC | 0.0701095 | TRUE | S-methyl-5'-thioadenosine deaminase |
| | | EC5.3.1.23 | EC | 0.0775393 | TRUE | S-methyl-5-thioribose-1-phosphate isomerase |
| | | EC2.4.2.44 | EC | NA | FALSE | S-methyl-5'-thioinosine phosphorylase |
| Wang et al. (ESRD) | 49 | C04556 | Metabolite | NA | FALSE | 4-Amino-2-methyl-5-(phosphooxymethyl)pyrimidine |
| | | C04751 | Metabolite | NA | FALSE | 1-(5-Phospho-D-ribosyl)-5-amino-4-imidazolecarboxylate |
| | | C03373 | Metabolite | NA | FALSE | Aminoimidazole ribotide |
| | | C15667 | Metabolite | NA | FALSE | 5-Carboxyamino-1-(5-phospho-D-ribosyl)imidazole |
| | | EC2.7.1.49 | EC | 0.0008382 | TRUE | hydroxymethylpyrimidine kinase |
| | | EC2.7.4.7 | EC | 0.0065149 | TRUE | phosphooxymethylpyrimidine kinase |
| | | EC4.1.99.17 | EC | 0.0033584 | TRUE | phosphomethylpyrimidine synthase |
| | | EC5.4.99.18 | EC | 0.0119016 | TRUE | 5-(carboxyamino)imidazole ribonucleotide mutase |
| | | EC6.3.2.6 | EC | 0.0403274 | TRUE | phosphoribosylaminoimidazolesuccinocarboxamide synthase |
| | | EC6.3.3.1 | EC | 0.0528415 | TRUE | phosphoribosylformylglycinamidine cyclo-ligase |
| | | EC6.3.4.18 | EC | 0.0076346 | TRUE | 5-(carboxyamino)imidazole ribonucleotide synthase |
| Wang et al. (ESRD) | 51 | C00245 | Metabolite | NA | FALSE | Taurine |
| | | C00099 | Metabolite | NA | FALSE | beta-Alanine |
| | | C00334 | Metabolite | NA | FALSE | 4-Aminobutanoate |
| | | EC3.5.1.24 | EC | 0.0939251 | TRUE | choloylglycine hydrolase |
| | | EC3.5.1.94 | EC | 0.0905836 | TRUE | gamma-glutamyl-gamma-aminobutyrate hydrolase |
| | | EC4.1.1.11 | EC | 0.000978 | TRUE | aspartate 1-decarboxylase |
| | | EC4.1.1.15 | EC | 0.0007686 | TRUE | glutamate decarboxylase |
| | | EC6.3.2.1 | EC | 0.0125214 | TRUE | pantoate--beta-alanine ligase (AMP-forming) |

### Supplementary Table S7: Modules significant overlaps with KEGG pathways

| Dataset | Module ID | KEGG pathway code | KEGG pathway description | Module size | Pathway size | No. of EC nodes in overlap | No. of metabolite nodes in overlap | Hypergeometric test - FDR |
|---|---|---|---|---|---|---|---|---|
| Franzosa et al. (CD) | 1 | map00220 | Arginine biosynthesis | 8 | 38 | 1 | 2 | 0.0000416 |
| Franzosa et al. (CD) | 1 | map00330 | Arginine and proline metabolism | 8 | 119 | 2 | 2 | 0.0000402 |
| Franzosa et al. (CD) | 2 | map00030 | Pentose phosphate pathway | 14 | 82 | 3 | 2 | 0.0000055 |
| Franzosa et al. (CD) | 2 | map00230 | Purine metabolism | 14 | 170 | 1 | 3 | 0.0013823 |
| Franzosa et al. (CD) | 2 | map00240 | Pyrimidine metabolism | 14 | 113 | 1 | 2 | 0.0042623 |
| Franzosa et al. (CD) | 2 | map00250 | Alanine, aspartate and glutamate metabolism | 14 | 49 | 2 | 1 | 0.0004182 |
| Franzosa et al. (CD) | 2 | map00750 | Vitamin B6 metabolism | 14 | 42 | 2 | 4 | 0 |
| Franzosa et al. (CD) | 2 | map01230 | Biosynthesis of amino acids | 14 | 113 | 0 | 3 | 0.0042623 |
| Franzosa et al. (CD) | 2 | map01240 | Biosynthesis of cofactors | 14 | 297 | 0 | 4 | 0.0095605 |
| Franzosa et al. (CD) | 3 | map00270 | Cysteine and methionine metabolism | 7 | 129 | 3 | 4 | 0 |
| Franzosa et al. (CD) | 5 | map00280 | Valine, leucine and isoleucine degradation | 7 | 65 | 1 | 4 | 0 |
| Franzosa et al. (CD) | 5 | map00290 | Valine, leucine and isoleucine biosynthesis | 7 | 33 | 2 | 5 | 0 |
| Franzosa et al. (CD) | 5 | map00770 | Pantothenate and CoA biosynthesis | 7 | 47 | 1 | 2 | 0.0000475 |
| Franzosa et al. (CD) | 5 | map00966 | Glucosinolate biosynthesis | 7 | 11 | 1 | 4 | 0 |
| Franzosa et al. (CD) | 5 | map00970 | Aminoacyl-tRNA biosynthesis | 7 | 61 | 0 | 3 | 0.0000967 |
| Franzosa et al. (CD) | 5 | map01060 | Biosynthesis of plant secondary metabolites | 7 | 96 | 0 | 3 | 0.0003311 |
| Franzosa et al. (CD) | 5 | map01210 | 2-Oxocarboxylic acid metabolism | 7 | 63 | 0 | 5 | 0 |
| Franzosa et al. (CD) | 5 | map01230 | Biosynthesis of amino acids | 7 | 113 | 0 | 5 | 0.0000005 |
| Franzosa et al. (CD) | 5 | map02010 | ABC transporters | 7 | 76 | 0 | 3 | 0.0001766 |
| Franzosa et al. (CD) | 5 | map04974 | Protein digestion and absorption | 7 | 27 | 0 | 3 | 0.000011 |
| Franzosa et al. (CD) | 5 | map04978 | Mineral absorption | 7 | 17 | 0 | 3 | 0.0000034 |
| Franzosa et al. (CD) | 5 | map05230 | Central carbon metabolism in cancer | 7 | 25 | 0 | 3 | 0.0000092 |
| Franzosa et al. (CD) | 7 | map00230 | Purine metabolism | 26 | 170 | 12 | 9 | 0 |
| Franzosa et al. (CD) | 7 | map00240 | Pyrimidine metabolism | 26 | 113 | 2 | 2 | 0.0033751 |
| Franzosa et al. (CD) | 7 | map00760 | Nicotinate and nicotinamide metabolism | 26 | 93 | 2 | 1 | 0.0139411 |
| Franzosa et al. (CD) | 7 | map01060 | Biosynthesis of plant secondary metabolites | 26 | 96 | 0 | 5 | 0.0001915 |
| Franzosa et al. (CD) | 7 | map01065 | Biosynthesis of alkaloids derived from histidine and purine | 26 | 27 | 0 | 4 | 0.000021 |
| Franzosa et al. (CD) | 7 | map01232 | Nucleotide metabolism | 26 | 57 | 0 | 10 | 0 |
| Franzosa et al. (CD) | 7 | map02010 | ABC transporters | 26 | 76 | 0 | 4 | 0.0008103 |
| Franzosa et al. (CD) | 9 | map00620 | Pyruvate metabolism | 5 | 75 | 2 | 1 | 0.0000551 |
| Franzosa et al. (CD) | 9 | map00640 | Propanoate metabolism | 5 | 79 | 2 | 3 | 0 |
| Franzosa et al. (CD) | 11 | map00565 | Ether lipid metabolism | 6 | 26 | 2 | 1 | 0.0000064 |
| Franzosa et al. (CD) | 11 | map00591 | Linoleic acid metabolism | 6 | 8 | 1 | 2 | 0.0000002 |
| Franzosa et al. (CD) | 11 | map01040 | Biosynthesis of unsaturated fatty acids | 6 | 31 | 0 | 3 | 0.0000101 |
| Franzosa et al. (CD) | 12 | map00120 | Primary bile acid biosynthesis | 6 | 19 | 1 | 5 | 0 |
| Franzosa et al. (CD) | 12 | map00121 | Secondary bile acid biosynthesis | 6 | 9 | 1 | 4 | 0 |
| Franzosa et al. (CD) | 12 | map04976 | Bile secretion | 6 | 34 | 0 | 4 | 0.0000001 |
| Franzosa et al. (CD) | 13 | map00730 | Thiamine metabolism | 8 | 48 | 2 | 1 | 0.0000756 |
| Franzosa et al. (CD) | 13 | map00740 | Riboflavin metabolism | 8 | 47 | 3 | 2 | 0 |
| Franzosa et al. (CD) | 13 | map01240 | Biosynthesis of cofactors | 8 | 297 | 0 | 3 | 0.0110714 |
| Franzosa et al. (CD) | 13 | map04977 | Vitamin digestion and absorption | 8 | 29 | 0 | 3 | 0.000021 |
| Franzosa et al. (CD) | 15 | map00260 | Glycine, serine and threonine metabolism | 5 | 102 | 2 | 3 | 0 |
| Franzosa et al. (CD) | 15 | map00330 | Arginine and proline metabolism | 5 | 119 | 1 | 2 | 0.000193 |
| Franzosa et al. (CD) | 17 | map00350 | Tyrosine metabolism | 11 | 80 | 3 | 1 | 0.0000379 |
| Franzosa et al. (CD) | 17 | map00360 | Phenylalanine metabolism | 11 | 78 | 4 | 2 | 0 |
| Franzosa et al. (CD) | 17 | map00400 | Phenylalanine, tyrosine and tryptophan biosynthesis | 11 | 62 | 3 | 2 | 0.0000005 |
| Franzosa et al. (CD) | 17 | map00401 | Novobiocin biosynthesis | 11 | 21 | 2 | 1 | 0.0000216 |
| Franzosa et al. (CD) | 17 | map00470 | D-Amino acid metabolism | 11 | 84 | 2 | 2 | 0.000043 |
| Franzosa et al. (CD) | 17 | map00960 | Tropane, piperidine and pyridine alkaloid biosynthesis | 11 | 32 | 2 | 2 | 0.0000015 |
| Franzosa et al. (CD) | 21 | map00040 | Pentose and glucuronate interconversions | 18 | 109 | 2 | 1 | 0.0078875 |
| Franzosa et al. (CD) | 21 | map00240 | Pyrimidine metabolism | 18 | 113 | 4 | 2 | 0.0000055 |
| Franzosa et al. (CD) | 21 | map00500 | Starch and sucrose metabolism | 18 | 91 | 4 | 1 | 0.0000281 |
| Franzosa et al. (CD) | 21 | map00670 | One carbon pool by folate | 18 | 29 | 4 | 2 | 0 |
| Franzosa et al. (CD) | 21 | map00760 | Nicotinate and nicotinamide metabolism | 18 | 93 | 3 | 1 | 0.0004182 |
| Franzosa et al. (CD) | 21 | map01240 | Biosynthesis of cofactors | 18 | 297 | 0 | 5 | 0.0043585 |
| Franzosa et al. (CD) | 21 | map01523 | Antifolate resistance | 18 | 13 | 0 | 3 | 0.0000221 |
| Franzosa et al. (CD) | 22 | map00010 | Glycolysis / Gluconeogenesis | 10 | 67 | 2 | 1 | 0.0003711 |
| Franzosa et al. (CD) | 22 | map00051 | Fructose and mannose metabolism | 10 | 116 | 3 | 2 | 0.0000045 |
| Franzosa et al. (CD) | 22 | map00500 | Starch and sucrose metabolism | 10 | 91 | 3 | 2 | 0.0000015 |
| Franzosa et al. (CD) | 22 | map00520 | Amino sugar and nucleotide sugar metabolism | 10 | 215 | 4 | 2 | 0.0000037 |
| Franzosa et al. (CD) | 22 | map00541 | O-Antigen nucleotide sugar biosynthesis | 10 | 118 | 1 | 2 | 0.0017899 |
| Franzosa et al. (CD) | 26 | map00520 | Amino sugar and nucleotide sugar metabolism | 4 | 215 | 3 | 1 | 0.0000087 |
| Franzosa et al. (UC) | 1 | map00260 | Glycine, serine and threonine metabolism | 8 | 102 | 2 | 1 | 0.0006315 |
| Franzosa et al. (UC) | 1 | map00270 | Cysteine and methionine metabolism | 8 | 129 | 2 | 1 | 0.0012005 |
| Franzosa et al. (UC) | 1 | map00310 | Lysine degradation | 8 | 65 | 2 | 1 | 0.0001992 |
| Franzosa et al. (UC) | 1 | map00470 | D-Amino acid metabolism | 8 | 84 | 3 | 3 | 0 |
| Franzosa et al. (UC) | 2 | map00270 | Cysteine and methionine metabolism | 21 | 129 | 9 | 3 | 0 |
| Franzosa et al. (UC) | 2 | map00350 | Tyrosine metabolism | 21 | 80 | 4 | 1 | 0.0000342 |
| Franzosa et al. (UC) | 2 | map00360 | Phenylalanine metabolism | 21 | 78 | 4 | 0 | 0.0004432 |
| Franzosa et al. (UC) | 2 | map00400 | Phenylalanine, tyrosine and tryptophan biosynthesis | 21 | 62 | 4 | 2 | 0.0000004 |
| Franzosa et al. (UC) | 2 | map00401 | Novobiocin biosynthesis | 21 | 21 | 4 | 1 | 0.0000001 |
| Franzosa et al. (UC) | 2 | map00450 | Selenocompound metabolism | 21 | 31 | 5 | 1 | 0 |
| Franzosa et al. (UC) | 2 | map00950 | Isoquinoline alkaloid biosynthesis | 21 | 14 | 2 | 1 | 0.0000438 |
| Franzosa et al. (UC) | 2 | map00960 | Tropane, piperidine and pyridine alkaloid biosynthesis | 21 | 32 | 3 | 0 | 0.0004564 |
| Franzosa et al. (UC) | 2 | map01230 | Biosynthesis of amino acids | 21 | 113 | 0 | 4 | 0.0015885 |
| Franzosa et al. (UC) | 5 | map00010 | Glycolysis / Gluconeogenesis | 11 | 67 | 3 | 1 | 0.0000204 |
| Franzosa et al. (UC) | 5 | map00620 | Pyruvate metabolism | 11 | 75 | 7 | 1 | 0 |
| Franzosa et al. (UC) | 5 | map00640 | Propanoate metabolism | 11 | 79 | 7 | 2 | 0 |
| Franzosa et al. (UC) | 6 | map00230 | Purine metabolism | 30 | 170 | 10 | 8 | 0 |
| Franzosa et al. (UC) | 6 | map00240 | Pyrimidine metabolism | 30 | 113 | 10 | 3 | 0 |
| Franzosa et al. (UC) | 6 | map00983 | Drug metabolism - other enzymes | 30 | 58 | 4 | 1 | 0.0000438 |
| Franzosa et al. (UC) | 6 | map01232 | Nucleotide metabolism | 30 | 57 | 0 | 10 | 0 |

| | | | | | | | | |
|---|---|---|---|---|---|---|---|---|
| Franzosa et al. (UC) | 6 | map02010 | ABC transporters | 30 | 76 | 0 | 5 | 0.0001522 |
| Franzosa et al. (UC) | 8 | map00564 | Glycerophospholipid metabolism | 7 | 81 | 4 | 1 | 0.0000001 |
| Franzosa et al. (UC) | 8 | map00591 | Linoleic acid metabolism | 7 | 8 | 1 | 2 | 0.0000003 |
| Franzosa et al. (UC) | 14 | map00010 | Glycolysis / Gluconeogenesis | 14 | 67 | 3 | 2 | 0.000002 |
| Franzosa et al. (UC) | 14 | map00020 | Citrate cycle (TCA cycle) | 14 | 39 | 5 | 2 | 0 |
| Franzosa et al. (UC) | 14 | map00260 | Glycine, serine and threonine metabolism | 14 | 102 | 2 | 1 | 0.0033262 |
| Franzosa et al. (UC) | 14 | map00280 | Valine, leucine and isoleucine degradation | 14 | 65 | 1 | 2 | 0.0009869 |
| Franzosa et al. (UC) | 14 | map00310 | Lysine degradation | 14 | 65 | 2 | 2 | 0.0000476 |
| Franzosa et al. (UC) | 14 | map00620 | Pyruvate metabolism | 14 | 75 | 3 | 2 | 0.0000033 |
| Franzosa et al. (UC) | 14 | map00640 | Propanoate metabolism | 14 | 79 | 1 | 2 | 0.001666 |
| Franzosa et al. (UC) | 14 | map00785 | Lipoic acid metabolism | 14 | 48 | 9 | 5 | 0 |
| Franzosa et al. (UC) | 14 | map01240 | Biosynthesis of cofactors | 14 | 297 | 0 | 4 | 0.0098416 |
| Franzosa et al. (UC) | 18 | map00010 | Glycolysis / Gluconeogenesis | 13 | 67 | 4 | 2 | 0 |
| Franzosa et al. (UC) | 18 | map00030 | Pentose phosphate pathway | 13 | 82 | 3 | 2 | 0.0000033 |
| Franzosa et al. (UC) | 18 | map00051 | Fructose and mannose metabolism | 13 | 116 | 6 | 2 | 0 |
| Franzosa et al. (UC) | 18 | map00500 | Starch and sucrose metabolism | 13 | 91 | 2 | 1 | 0.0019717 |
| Franzosa et al. (UC) | 18 | map00520 | Amino sugar and nucleotide sugar metabolism | 13 | 215 | 3 | 2 | 0.0002541 |
| Franzosa et al. (UC) | 18 | map00541 | O-Antigen nucleotide sugar biosynthesis | 13 | 118 | 1 | 2 | 0.0039508 |
| Franzosa et al. (UC) | 23 | map00260 | Glycine, serine and threonine metabolism | 12 | 102 | 2 | 1 | 0.0021132 |
| Franzosa et al. (UC) | 23 | map00361 | Chlorocyclohexane and chlorobenzene degradation | 12 | 81 | 2 | 1 | 0.0011536 |
| Franzosa et al. (UC) | 23 | map00625 | Chloroalkane and chloroalkene degradation | 12 | 45 | 2 | 1 | 0.00024 |
| Franzosa et al. (UC) | 23 | map00630 | Glyoxylate and dicarboxylate metabolism | 12 | 86 | 7 | 3 | 0 |
| Franzosa et al. (UC) | 24 | map00250 | Alanine, aspartate and glutamate metabolism | 15 | 49 | 2 | 1 | 0.0005533 |
| Franzosa et al. (UC) | 24 | map00310 | Lysine degradation | 15 | 65 | 2 | 1 | 0.0012005 |
| Franzosa et al. (UC) | 24 | map00330 | Arginine and proline metabolism | 15 | 119 | 3 | 1 | 0.0005421 |
| Franzosa et al. (UC) | 24 | map00410 | beta-Alanine metabolism | 15 | 39 | 4 | 3 | 0 |
| Franzosa et al. (UC) | 24 | map00640 | Propanoate metabolism | 15 | 79 | 2 | 2 | 0.0001341 |
| Franzosa et al. (UC) | 24 | map00650 | Butanoate metabolism | 15 | 88 | 2 | 1 | 0.0027201 |
| Franzosa et al. (UC) | 41 | map00071 | Fatty acid degradation | 13 | 47 | 3 | 3 | 0 |
| Franzosa et al. (UC) | 41 | map00310 | Lysine degradation | 13 | 65 | 1 | 2 | 0.0007883 |
| Franzosa et al. (UC) | 41 | map00362 | Benzoate degradation | 13 | 118 | 2 | 1 | 0.0039508 |
| Franzosa et al. (UC) | 41 | map00650 | Butanoate metabolism | 13 | 88 | 5 | 2 | 0 |
| Franzosa et al. (UC) | 41 | map00720 | Carbon fixation pathways in prokaryotes | 13 | 83 | 1 | 2 | 0.0015492 |
| Franzosa et al. (UC) | 41 | map01200 | Carbon metabolism | 13 | 92 | 0 | 3 | 0.0020222 |
| Franzosa et al. (UC) | 41 | map01212 | Fatty acid metabolism | 13 | 64 | 0 | 3 | 0.0007633 |
| Franzosa et al. (UC) | 45 | map00130 | Ubiquinone and other terpenoid-quinone biosynthesis | 7 | 84 | 2 | 2 | 0.0000063 |
| Franzosa et al. (UC) | 45 | map01053 | Biosynthesis of siderophore group nonribosomal peptides | 7 | 19 | 2 | 2 | 0 |
| Franzosa et al. (UC) | 46 | map00061 | Fatty acid biosynthesis | 12 | 50 | 1 | 2 | 0.000312 |
| Franzosa et al. (UC) | 46 | map00071 | Fatty acid degradation | 12 | 47 | 1 | 2 | 0.0002651 |
| Franzosa et al. (UC) | 46 | map00540 | Lipopolysaccharide biosynthesis | 12 | 77 | 3 | 3 | 0 |
| Franzosa et al. (UC) | 46 | map00770 | Pantothenate and CoA biosynthesis | 12 | 47 | 3 | 2 | 0.0000002 |
| Franzosa et al. (UC) | 48 | map00240 | Pyrimidine metabolism | 10 | 113 | 3 | 1 | 0.0000917 |
| Franzosa et al. (UC) | 48 | map00670 | One carbon pool by folate | 10 | 29 | 6 | 2 | 0 |
| Franzosa et al. (UC) | 48 | map00720 | Carbon fixation pathways in prokaryotes | 10 | 83 | 2 | 2 | 0.0000296 |
| Franzosa et al. (UC) | 61 | map00910 | Nitrogen metabolism | 6 | 41 | 2 | 2 | 0.0000002 |
| Jeffery et al. (IBS) | 2 | map00220 | Arginine biosynthesis | 25 | 38 | 3 | 2 | 0.000003 |
| Jeffery et al. (IBS) | 2 | map00250 | Alanine, aspartate and glutamate metabolism | 25 | 49 | 6 | 1 | 0 |
| Jeffery et al. (IBS) | 2 | map00260 | Glycine, serine and threonine metabolism | 25 | 102 | 6 | 2 | 0.0000001 |
| Jeffery et al. (IBS) | 2 | map00270 | Cysteine and methionine metabolism | 25 | 129 | 2 | 1 | 0.030691 |
| Jeffery et al. (IBS) | 2 | map00300 | Lysine biosynthesis | 25 | 59 | 3 | 2 | 0.0000218 |
| Jeffery et al. (IBS) | 2 | map00310 | Lysine degradation | 25 | 65 | 2 | 1 | 0.0051024 |
| Jeffery et al. (IBS) | 2 | map00330 | Arginine and proline metabolism | 25 | 119 | 2 | 1 | 0.0251289 |
| Jeffery et al. (IBS) | 2 | map00470 | D-Amino acid metabolism | 25 | 84 | 3 | 5 | 0 |
| Jeffery et al. (IBS) | 2 | map00630 | Glyoxylate and dicarboxylate metabolism | 25 | 86 | 2 | 2 | 0.001212 |
| Jeffery et al. (IBS) | 2 | map00680 | Methane metabolism | 25 | 156 | 2 | 1 | 0.0489446 |
| Jeffery et al. (IBS) | 2 | map00910 | Nitrogen metabolism | 25 | 41 | 2 | 1 | 0.0015025 |
| Jeffery et al. (IBS) | 2 | map00970 | Aminoacyl-tRNA biosynthesis | 25 | 61 | 0 | 5 | 0.0000248 |
| Jeffery et al. (IBS) | 2 | map00997 | Biosynthesis of various other secondary metabolites | 25 | 46 | 0 | 3 | 0.0020471 |
| Jeffery et al. (IBS) | 2 | map01060 | Biosynthesis of plant secondary metabolites | 25 | 96 | 0 | 5 | 0.0001732 |
| Jeffery et al. (IBS) | 2 | map01230 | Biosynthesis of amino acids | 25 | 113 | 0 | 6 | 0.0000353 |
| Jeffery et al. (IBS) | 2 | map02010 | ABC transporters | 25 | 76 | 0 | 4 | 0.0007763 |
| Jeffery et al. (IBS) | 2 | map04974 | Protein digestion and absorption | 25 | 27 | 0 | 5 | 0.0000006 |
| Jeffery et al. (IBS) | 2 | map04978 | Mineral absorption | 25 | 17 | 0 | 3 | 0.0001318 |
| Jeffery et al. (IBS) | 2 | map05230 | Central carbon metabolism in cancer | 25 | 25 | 0 | 4 | 0.0000144 |
| Jeffery et al. (IBS) | 4 | map00230 | Purine metabolism | 18 | 170 | 6 | 3 | 0 |
| Jeffery et al. (IBS) | 4 | map00760 | Nicotinate and nicotinamide metabolism | 18 | 93 | 7 | 3 | 0 |
| Jeffery et al. (IBS) | 4 | map01232 | Nucleotide metabolism | 18 | 57 | 0 | 3 | 0.0014778 |
| Jeffery et al. (IBS) | 16 | map00071 | Fatty acid degradation | 12 | 47 | 2 | 3 | 0.0000002 |
| Jeffery et al. (IBS) | 16 | map00280 | Valine, leucine and isoleucine degradation | 12 | 65 | 2 | 1 | 0.000644 |
| Jeffery et al. (IBS) | 16 | map00310 | Lysine degradation | 12 | 65 | 1 | 3 | 0.0000272 |
| Jeffery et al. (IBS) | 16 | map00362 | Benzoate degradation | 12 | 118 | 1 | 2 | 0.0032695 |
| Jeffery et al. (IBS) | 16 | map00630 | Glyoxylate and dicarboxylate metabolism | 12 | 86 | 1 | 2 | 0.001415 |
| Jeffery et al. (IBS) | 16 | map00650 | Butanoate metabolism | 12 | 88 | 7 | 4 | 0 |
| Jeffery et al. (IBS) | 16 | map01200 | Carbon metabolism | 12 | 92 | 0 | 3 | 0.0016749 |
| Jeffery et al. (IBS) | 16 | map01212 | Fatty acid metabolism | 12 | 64 | 0 | 3 | 0.0006187 |
| Jeffery et al. (IBS) | 33 | map00240 | Pyrimidine metabolism | 16 | 113 | 10 | 6 | 0 |
| Jeffery et al. (IBS) | 33 | map00410 | beta-Alanine metabolism | 16 | 39 | 2 | 1 | 0.000367 |
| Jeffery et al. (IBS) | 33 | map00770 | Pantothenate and CoA biosynthesis | 16 | 47 | 2 | 1 | 0.0006143 |
| Jeffery et al. (IBS) | 33 | map00983 | Drug metabolism - other enzymes | 16 | 58 | 4 | 0 | 0.0000547 |
| Jeffery et al. (IBS) | 33 | map01232 | Nucleotide metabolism | 16 | 57 | 0 | 5 | 0.0000022 |
| Jeffery et al. (IBS) | 38 | map00730 | Thiamine metabolism | 15 | 48 | 4 | 2 | 0 |
| Jeffery et al. (IBS) | 38 | map00740 | Riboflavin metabolism | 15 | 47 | 5 | 3 | 0 |
| Jeffery et al. (IBS) | 38 | map01240 | Biosynthesis of cofactors | 15 | 297 | 0 | 5 | 0.0020407 |
| Jeffery et al. (IBS) | 38 | map04977 | Vitamin digestion and absorption | 15 | 29 | 0 | 3 | 0.0001381 |
| Jeffery et al. (IBS) | 40 | map00860 | Porphyrin metabolism | 5 | 186 | 3 | 2 | 0.0000003 |
| Jeffery et al. (IBS) | 47 | map00790 | Folate biosynthesis | 5 | 97 | 3 | 2 | 0 |
| Jeffery et al. (IBS) | 48 | map00730 | Thiamine metabolism | 5 | 48 | 3 | 2 | 0 |
| Jeffery et al. (IBS) | 51 | map00780 | Biotin metabolism | 7 | 34 | 4 | 3 | 0 |
| Jeffery et al. (IBS) | 51 | map01240 | Biosynthesis of cofactors | 7 | 297 | 0 | 3 | 0.0076235 |
| Jeffery et al. (IBS) | 53 | map00040 | Pentose and glucuronate interconversions | 25 | 109 | 1 | 2 | 0.0200421 |

| | | | | | | | | |
|---|---|---|---|---|---|---|---|---|
| Jeffery et al. (IBS) | 53 | map00052 | Galactose metabolism | 25 | 73 | 3 | 4 | 0.0000002 |
| Jeffery et al. (IBS) | 53 | map00500 | Starch and sucrose metabolism | 25 | 91 | 15 | 6 | 0 |
| Jeffery et al. (IBS) | 53 | map00520 | Amino sugar and nucleotide sugar metabolism | 25 | 215 | 4 | 3 | 0.0001362 |
| Jeffery et al. (IBS) | 53 | map00541 | O-Antigen nucleotide sugar biosynthesis | 25 | 118 | 3 | 3 | 0.0000432 |
| Jeffery et al. (IBS) | 53 | map00561 | Glycerolipid metabolism | 25 | 56 | 1 | 2 | 0.0034615 |
| Jeffery et al. (IBS) | 53 | map01250 | Biosynthesis of nucleotide sugars | 25 | 139 | 0 | 3 | 0.0368771 |
| Jeffery et al. (IBS) | 59 | map00900 | Terpenoid backbone biosynthesis | 5 | 50 | 4 | 1 | 0 |
| Jeffery et al. (IBS) | 60 | map00860 | Porphyrin metabolism | 8 | 186 | 5 | 3 | 0 |
| Jeffery et al. (IBS) | 68 | map00220 | Arginine biosynthesis | 5 | 38 | 3 | 2 | 0 |
| Jeffery et al. (IBS) | 68 | map00791 | Atrazine degradation | 5 | 18 | 3 | 2 | 0 |
| Wang et al. (ESRD) | 6 | map00260 | Glycine, serine and threonine metabolism | 6 | 102 | 4 | 2 | 0 |
| Wang et al. (ESRD) | 7 | map00040 | Pentose and glucuronate interconversions | 8 | 109 | 3 | 2 | 0.0000008 |
| Wang et al. (ESRD) | 7 | map00051 | Fructose and mannose metabolism | 8 | 116 | 2 | 1 | 0.0008585 |
| Wang et al. (ESRD) | 7 | map00052 | Galactose metabolism | 8 | 73 | 2 | 1 | 0.0002393 |
| Wang et al. (ESRD) | 8 | map00561 | Glycerolipid metabolism | 6 | 56 | 3 | 1 | 0.0000007 |
| Wang et al. (ESRD) | 8 | map00930 | Caprolactam degradation | 6 | 20 | 2 | 1 | 0.0000031 |
| Wang et al. (ESRD) | 9 | map00340 | Histidine metabolism | 8 | 44 | 4 | 3 | 0 |
| Wang et al. (ESRD) | 10 | map00750 | Vitamin B6 metabolism | 6 | 42 | 4 | 2 | 0 |
| Wang et al. (ESRD) | 11 | map00730 | Thiamine metabolism | 8 | 48 | 3 | 2 | 0 |
| Wang et al. (ESRD) | 11 | map00900 | Terpenoid backbone biosynthesis | 8 | 50 | 3 | 2 | 0 |
| Wang et al. (ESRD) | 12 | map00260 | Glycine, serine and threonine metabolism | 9 | 102 | 2 | 2 | 0.0000395 |
| Wang et al. (ESRD) | 12 | map00620 | Pyruvate metabolism | 9 | 75 | 3 | 1 | 0.0000134 |
| Wang et al. (ESRD) | 12 | map00630 | Glyoxylate and dicarboxylate metabolism | 9 | 86 | 3 | 2 | 0.0000006 |
| Wang et al. (ESRD) | 14 | map00620 | Pyruvate metabolism | 6 | 75 | 3 | 1 | 0.0000022 |
| Wang et al. (ESRD) | 14 | map00680 | Methane metabolism | 6 | 156 | 2 | 2 | 0.0000288 |
| Wang et al. (ESRD) | 14 | map00710 | Carbon fixation in photosynthetic organisms | 6 | 35 | 2 | 1 | 0.0000139 |
| Wang et al. (ESRD) | 15 | map00030 | Pentose phosphate pathway | 10 | 82 | 4 | 2 | 0 |
| Wang et al. (ESRD) | 15 | map00480 | Glutathione metabolism | 10 | 47 | 4 | 1 | 0.0000001 |
| Wang et al. (ESRD) | 16 | map00650 | Butanoate metabolism | 6 | 88 | 2 | 2 | 0.0000039 |
| Wang et al. (ESRD) | 16 | map00720 | Carbon fixation pathways in prokaryotes | 6 | 83 | 3 | 2 | 0 |
| Wang et al. (ESRD) | 17 | map00010 | Glycolysis / Gluconeogenesis | 15 | 67 | 2 | 2 | 0.000069 |
| Wang et al. (ESRD) | 17 | map00020 | Citrate cycle (TCA cycle) | 15 | 39 | 2 | 2 | 0.0000105 |
| Wang et al. (ESRD) | 17 | map00280 | Valine, leucine and isoleucine degradation | 15 | 65 | 4 | 5 | 0 |
| Wang et al. (ESRD) | 17 | map00290 | Valine, leucine and isoleucine biosynthesis | 15 | 33 | 1 | 2 | 0.0001782 |
| Wang et al. (ESRD) | 17 | map00310 | Lysine degradation | 15 | 65 | 1 | 2 | 0.0011322 |
| Wang et al. (ESRD) | 17 | map00620 | Pyruvate metabolism | 15 | 75 | 2 | 2 | 0.0001023 |
| Wang et al. (ESRD) | 17 | map00640 | Propanoate metabolism | 15 | 79 | 3 | 2 | 0.0000066 |
| Wang et al. (ESRD) | 17 | map00785 | Lipoic acid metabolism | 15 | 48 | 5 | 6 | 0 |
| Wang et al. (ESRD) | 17 | map01210 | 2-Oxocarboxylic acid metabolism | 15 | 63 | 0 | 3 | 0.0010414 |
| Wang et al. (ESRD) | 17 | map01240 | Biosynthesis of cofactors | 15 | 297 | 0 | 4 | 0.012395 |
| Wang et al. (ESRD) | 19 | map00300 | Lysine biosynthesis | 10 | 59 | 7 | 3 | 0 |
| Wang et al. (ESRD) | 19 | map01230 | Biosynthesis of amino acids | 10 | 113 | 0 | 3 | 0.0015409 |
| Wang et al. (ESRD) | 24 | map00400 | Phenylalanine, tyrosine and tryptophan biosynthesis | 4 | 62 | 3 | 1 | 0.0000001 |
| Wang et al. (ESRD) | 26 | map00860 | Porphyrin metabolism | 8 | 186 | 5 | 3 | 0 |
| Wang et al. (ESRD) | 26 | map01240 | Biosynthesis of cofactors | 8 | 297 | 0 | 3 | 0.0112675 |
| Wang et al. (ESRD) | 27 | map00561 | Glycerolipid metabolism | 8 | 56 | 2 | 1 | 0.0001198 |
| Wang et al. (ESRD) | 27 | map00760 | Nicotinate and nicotinamide metabolism | 8 | 93 | 2 | 2 | 0.0000172 |
| Wang et al. (ESRD) | 29 | map00030 | Pentose phosphate pathway | 9 | 82 | 2 | 1 | 0.0004719 |
| Wang et al. (ESRD) | 29 | map00230 | Purine metabolism | 9 | 170 | 2 | 2 | 0.0002285 |
| Wang et al. (ESRD) | 29 | map00630 | Glyoxylate and dicarboxylate metabolism | 9 | 86 | 2 | 1 | 0.0005343 |
| Wang et al. (ESRD) | 29 | map00750 | Vitamin B6 metabolism | 9 | 42 | 1 | 2 | 0.0000778 |
| Wang et al. (ESRD) | 29 | map00910 | Nitrogen metabolism | 9 | 41 | 2 | 1 | 0.0000746 |
| Wang et al. (ESRD) | 30 | map00730 | Thiamine metabolism | 11 | 48 | 3 | 1 | 0.0000064 |
| Wang et al. (ESRD) | 30 | map00740 | Riboflavin metabolism | 11 | 47 | 5 | 1 | 0 |
| Wang et al. (ESRD) | 34 | map00220 | Arginine biosynthesis | 11 | 38 | 6 | 4 | 0 |
| Wang et al. (ESRD) | 34 | map00330 | Arginine and proline metabolism | 11 | 119 | 2 | 2 | 0.0001517 |
| Wang et al. (ESRD) | 34 | map01230 | Biosynthesis of amino acids | 11 | 113 | 0 | 4 | 0.0001251 |
| Wang et al. (ESRD) | 36 | map00780 | Biotin metabolism | 5 | 34 | 3 | 2 | 0 |
| Wang et al. (ESRD) | 37 | map00561 | Glycerolipid metabolism | 9 | 56 | 4 | 2 | 0 |
| Wang et al. (ESRD) | 37 | map00562 | Inositol phosphate metabolism | 9 | 51 | 2 | 2 | 0.0000035 |
| Wang et al. (ESRD) | 37 | map00564 | Glycerophospholipid metabolism | 9 | 81 | 3 | 2 | 0.0000005 |
| Wang et al. (ESRD) | 37 | map04070 | Phosphatidylinositol signaling system | 9 | 15 | 0 | 3 | 0.0000049 |
| Wang et al. (ESRD) | 38 | map00330 | Arginine and proline metabolism | 7 | 119 | 5 | 2 | 0 |
| Wang et al. (ESRD) | 38 | map00480 | Glutathione metabolism | 7 | 47 | 1 | 2 | 0.0000498 |
| Wang et al. (ESRD) | 40 | map00520 | Amino sugar and nucleotide sugar metabolism | 4 | 215 | 2 | 1 | 0.0004356 |
| Wang et al. (ESRD) | 41 | map00470 | D-Amino acid metabolism | 5 | 84 | 2 | 1 | 0.0000778 |
| Wang et al. (ESRD) | 41 | map00552 | Teichoic acid biosynthesis | 5 | 32 | 2 | 1 | 0.0000062 |
| Wang et al. (ESRD) | 42 | map00240 | Pyrimidine metabolism | 9 | 113 | 6 | 3 | 0 |
| Wang et al. (ESRD) | 42 | map00410 | beta-Alanine metabolism | 9 | 39 | 2 | 1 | 0.0000655 |
| Wang et al. (ESRD) | 42 | map00770 | Pantothenate and CoA biosynthesis | 9 | 47 | 2 | 1 | 0.0001062 |
| Wang et al. (ESRD) | 42 | map00983 | Drug metabolism - other enzymes | 9 | 58 | 3 | 0 | 0.0001852 |
| Wang et al. (ESRD) | 43 | map00010 | Glycolysis / Gluconeogenesis | 8 | 67 | 3 | 1 | 0.0000056 |
| Wang et al. (ESRD) | 43 | map00020 | Citrate cycle (TCA cycle) | 8 | 39 | 2 | 1 | 0.0000454 |
| Wang et al. (ESRD) | 43 | map00400 | Phenylalanine, tyrosine and tryptophan biosynthesis | 8 | 62 | 2 | 2 | 0.0000042 |
| Wang et al. (ESRD) | 43 | map00620 | Pyruvate metabolism | 8 | 75 | 4 | 1 | 0.0000002 |
| Wang et al. (ESRD) | 43 | map00680 | Methane metabolism | 8 | 156 | 2 | 1 | 0.0018833 |
| Wang et al. (ESRD) | 43 | map00710 | Carbon fixation in photosynthetic organisms | 8 | 35 | 2 | 1 | 0.0000341 |
| Wang et al. (ESRD) | 43 | map00720 | Carbon fixation pathways in prokaryotes | 8 | 83 | 2 | 1 | 0.0003446 |
| Wang et al. (ESRD) | 45 | map00520 | Amino sugar and nucleotide sugar metabolism | 5 | 215 | 3 | 2 | 0.0000005 |
| Wang et al. (ESRD) | 45 | map00541 | O-Antigen nucleotide sugar biosynthesis | 5 | 118 | 3 | 2 | 0 |
| Wang et al. (ESRD) | 46 | map00270 | Cysteine and methionine metabolism | 6 | 129 | 4 | 2 | 0 |
| Wang et al. (ESRD) | 49 | map00230 | Purine metabolism | 11 | 170 | 4 | 3 | 0.0000001 |
| Wang et al. (ESRD) | 49 | map00730 | Thiamine metabolism | 11 | 48 | 3 | 2 | 0.0000001 |
| Wang et al. (ESRD) | 51 | map00410 | beta-Alanine metabolism | 8 | 39 | 3 | 2 | 0 |
| Wang et al. (ESRD) | 51 | map00770 | Pantothenate and CoA biosynthesis | 8 | 47 | 2 | 1 | 0.000075 |
| Wang et al. (ESRD) | 51 | map04080 | Neuroactive ligand-receptor interaction | 8 | 27 | 0 | 3 | 0.0000171 |

* Only overlaps with significant modules are shown.

### *Supplementary Table S8: Pathway-disease associations (KEGG pathways) using multiple analysis approaches*

| Dataset | Test name | KEGG pathway ID | Pathway name | P-value | FDR | Direction |
|---|---|---|---|---|---|---|
| Wang et al. (ESRD) | Pathway-level DA | map00730 | Thiamine metabolism | 1.4E-10 | 6.97E-08 | Under-represented in disease |
| Wang et al. (ESRD) | Pathway-level DA | map00740 | Riboflavin metabolism | 5.62E-08 | 1.39E-05 | Under-represented in disease |
| Wang et al. (ESRD) | Pathway-level DA | map00930 | Caprolactam degradation | 5.7E-07 | 9.43E-05 | Under-represented in disease |
| Wang et al. (ESRD) | Pathway-level DA | map00562 | Inositol phosphate metabolism | 3.66E-06 | 0.000453 | Under-represented in disease |
| Wang et al. (ESRD) | Pathway-level DA | map00250 | Alanine, aspartate and glutamate metabolism | 1.24E-05 | 0.001226 | Under-represented in disease |
| Wang et al. (ESRD) | Pathway-level DA | map00332 | Carbapenem biosynthesis | 2.81E-05 | 0.002319 | Under-represented in disease |
| Wang et al. (ESRD) | Pathway-level DA | map00470 | D-Amino acid metabolism | 6.52E-05 | 0.00404 | Under-represented in disease |
| Wang et al. (ESRD) | Pathway-level DA | map01055 | Biosynthesis of vancomycin group antibiotics | 0.00022 | 0.008388 | Under-represented in disease |
| Wang et al. (ESRD) | Pathway-level DA | map00710 | Carbon fixation in photosynthetic organisms | 0.000269 | 0.008893 | Under-represented in disease |
| Wang et al. (ESRD) | Pathway-level DA | map00790 | Folate biosynthesis | 0.000268 | 0.008893 | Under-represented in disease |
| Wang et al. (ESRD) | Pathway-level DA | map00650 | Butanoate metabolism | 3.14E-04 | 0.009745 | Over-represented in disease |
| Wang et al. (ESRD) | Pathway-level DA | map00410 | beta-Alanine metabolism | 3.58E-04 | 0.009869 | Under-represented in disease |
| Wang et al. (ESRD) | Pathway-level DA | map00230 | Purine metabolism | 0.000473 | 0.01235 | Under-represented in disease |
| Wang et al. (ESRD) | Pathway-level DA | map00300 | Lysine biosynthesis | 6.19E-04 | 0.013698 | Under-represented in disease |
| Wang et al. (ESRD) | Pathway-level DA | map00561 | Glycerolipid metabolism | 5.91E-04 | 0.013698 | Over-represented in disease |
| Wang et al. (ESRD) | Pathway-level DA | map00625 | Chloroalkane and chloroalkene degradation | 6.54E-04 | 0.013698 | Under-represented in disease |
| Wang et al. (ESRD) | Pathway-level DA | map00900 | Terpenoid backbone biosynthesis | 5.94E-04 | 0.013698 | Under-represented in disease |
| Wang et al. (ESRD) | Pathway-level DA | map00525 | Acarbose and validamycin biosynthesis | 7.29E-04 | 0.013906 | Under-represented in disease |
| Wang et al. (ESRD) | Pathway-level DA | map00290 | Valine, leucine and isoleucine biosynthesis | 0.001071 | 0.018309 | Under-represented in disease |
| Wang et al. (ESRD) | Pathway-level DA | map00363 | Bisphenol degradation | 0.001315 | 0.019305 | Over-represented in disease |
| Wang et al. (ESRD) | Pathway-level DA | map00565 | Ether lipid metabolism | 0.001323 | 0.019305 | Over-represented in disease |
| Wang et al. (ESRD) | Pathway-level DA | map00220 | Arginine biosynthesis | 0.001624 | 0.022375 | Under-represented in disease |
| Wang et al. (ESRD) | Pathway-level DA | map00380 | Tryptophan metabolism | 0.001965 | 0.026335 | Over-represented in disease |
| Wang et al. (ESRD) | Pathway-level DA | map00195 | Photosynthesis | 0.00214 | 0.027162 | Under-represented in disease |
| Wang et al. (ESRD) | Pathway-level DA | map00280 | Valine, leucine and isoleucine degradation | 0.00219 | 0.027162 | Over-represented in disease |
| Wang et al. (ESRD) | Pathway-level DA | map00400 | Phenylalanine, tyrosine and tryptophan biosynthesis | 0.002721 | 0.030676 | Under-represented in disease |
| Wang et al. (ESRD) | Pathway-level DA | map00190 | Oxidative phosphorylation | 0.003417 | 0.036062 | Under-represented in disease |
| Wang et al. (ESRD) | Pathway-level DA | map00760 | Nicotinate and nicotinamide metabolism | 0.004138 | 0.039475 | Under-represented in disease |
| Wang et al. (ESRD) | Pathway-level DA | map00254 | Aflatoxin biosynthesis | 0.004274 | 0.039999 | Over-represented in disease |
| Wang et al. (ESRD) | Pathway-level DA | map00523 | Polyketide sugar unit biosynthesis | 0.004913 | 0.044308 | Under-represented in disease |
| Wang et al. (ESRD) | Pathway-level DA | map00030 | Pentose phosphate pathway | 0.005153 | 0.045639 | Over-represented in disease |
| Wang et al. (ESRD) | Pathway-level DA | map00640 | Propanoate metabolism | 0.005369 | 0.045914 | Over-represented in disease |
| Wang et al. (ESRD) | Pathway-level DA | map00460 | Cyanoamino acid metabolism | 0.006887 | 0.053934 | Under-represented in disease |
| Wang et al. (ESRD) | Pathway-level DA | map00660 | C5-Branched dibasic acid metabolism | 0.007164 | 0.054667 | Under-represented in disease |
| Wang et al. (ESRD) | Pathway-level DA | map00680 | Methane metabolism | 0.008692 | 0.060723 | Over-represented in disease |
| Wang et al. (ESRD) | Pathway-level DA | map00360 | Phenylalanine metabolism | 0.010326 | 0.068292 | Over-represented in disease |
| Wang et al. (ESRD) | Pathway-level DA | map00980 | Metabolism of xenobiotics by cytochrome P450 | 0.010259 | 0.068292 | Over-represented in disease |
| Wang et al. (ESRD) | Pathway-level DA | map00785 | Lipoic acid metabolism | 0.011633 | 0.074935 | Over-represented in disease |
| Wang et al. (ESRD) | Pathway-level DA | map00550 | Peptidoglycan biosynthesis | 0.012094 | 0.075929 | Under-represented in disease |
| Wang et al. (ESRD) | Pathway-level DA | map00480 | Glutathione metabolism | 0.013365 | 0.07987 | Over-represented in disease |
| Wang et al. (ESRD) | Pathway-level DA | map00520 | Amino sugar and nucleotide sugar metabolism | 0.014646 | 0.084471 | Over-represented in disease |
| Wang et al. (ESRD) | Pathway-level DA | map00908 | Zeatin biosynthesis | 0.015252 | 0.084982 | Under-represented in disease |
| Wang et al. (ESRD) | Pathway-level DA | map00052 | Galactose metabolism | 0.017638 | 0.095094 | Over-represented in disease |
| Wang et al. (ESRD) | Pathway-level DA | map00903 | Limonene degradation | 0.018196 | 0.097047 | Under-represented in disease |
| Franzosa et al. (CD) | Pathway-level DA | map00220 | Arginine biosynthesis | 3.87E-09 | 4.32E-06 | Under-represented in disease |
| Franzosa et al. (CD) | Pathway-level DA | map00332 | Carbapenem biosynthesis | 1.59E-06 | 4.45E-04 | Under-represented in disease |
| Franzosa et al. (CD) | Pathway-level DA | map00400 | Phenylalanine, tyrosine and tryptophan biosynthesis | 3.41E-06 | 7.60E-04 | Under-represented in disease |
| Franzosa et al. (CD) | Pathway-level DA | map00290 | Valine, leucine and isoleucine biosynthesis | 1.34E-05 | 0.002486 | Under-represented in disease |
| Franzosa et al. (CD) | Pathway-level DA | map00340 | Histidine metabolism | 7.88E-05 | 0.007703 | Under-represented in disease |
| Franzosa et al. (CD) | Pathway-level DA | map00030 | Pentose phosphate pathway | 1.03E-04 | 0.008693 | Over-represented in disease |
| Franzosa et al. (CD) | Pathway-level DA | map00520 | Amino sugar and nucleotide sugar metabolism | 0.001065 | 0.042465 | Over-represented in disease |
| Franzosa et al. (CD) | Pathway-level DA | map00052 | Galactose metabolism | 0.001401 | 0.052111 | Over-represented in disease |
| Franzosa et al. (CD) | Pathway-level DA | map00254 | Aflatoxin biosynthesis | 0.001929 | 0.061446 | Over-represented in disease |
| Franzosa et al. (CD) | Pathway-level DA | map00660 | C5-Branched dibasic acid metabolism | 0.001771 | 0.061446 | Under-represented in disease |
| Franzosa et al. (CD) | Pathway-level DA | map00730 | Thiamine metabolism | 0.001862 | 0.061446 | Under-represented in disease |
| Franzosa et al. (CD) | Pathway-level DA | map00480 | Glutathione metabolism | 0.002442 | 0.064888 | Over-represented in disease |
| Franzosa et al. (CD) | Pathway-level DA | map00950 | Isoquinoline alkaloid biosynthesis | 0.002879 | 0.074716 | Over-represented in disease |
| Franzosa et al. (UC) | Pathway-level DA | map00625 | Chloroalkane and chloroalkene degradation | 5.59E-06 | 0.002424 | Under-represented in disease |
| Franzosa et al. (UC) | Pathway-level DA | map00310 | Lysine degradation | 8.58E-05 | 0.011419 | Over-represented in disease |
| Franzosa et al. (UC) | Pathway-level DA | map00999 | Biosynthesis of various plant secondary metabolites | 1.72E-04 | 0.014893 | Under-represented in disease |
| Franzosa et al. (UC) | Pathway-level DA | map00400 | Phenylalanine, tyrosine and tryptophan biosynthesis | 2.11E-04 | 0.015238 | Under-represented in disease |
| Franzosa et al. (UC) | Pathway-level DA | map00270 | Cysteine and methionine metabolism | 3.39E-04 | 0.021011 | Under-represented in disease |
| Franzosa et al. (UC) | Pathway-level DA | map00450 | Selenocompound metabolism | 4.75E-04 | 0.021804 | Under-represented in disease |
| Franzosa et al. (UC) | Pathway-level DA | map00785 | Lipoic acid metabolism | 4.77E-04 | 0.021804 | Over-represented in disease |
| Franzosa et al. (UC) | Pathway-level DA | map00290 | Valine, leucine and isoleucine biosynthesis | 7.48E-04 | 0.029527 | Under-represented in disease |
| Franzosa et al. (UC) | Pathway-level DA | map00195 | Photosynthesis | 8.95E-04 | 0.032357 | Under-represented in disease |
| Franzosa et al. (UC) | Pathway-level DA | map00190 | Oxidative phosphorylation | 0.001223 | 0.040829 | Under-represented in disease |
| Franzosa et al. (UC) | Pathway-level DA | map00540 | Lipopolysaccharide biosynthesis | 0.001463 | 0.045522 | Over-represented in disease |
| Franzosa et al. (UC) | Pathway-level DA | map00430 | Taurine and hypotaurine metabolism | 0.001528 | 0.045747 | Over-represented in disease |
| Franzosa et al. (UC) | Pathway-level DA | map00440 | Phosphonate and phosphinate metabolism | 0.00206 | 0.049871 | Over-represented in disease |
| Franzosa et al. (UC) | Pathway-level DA | map00480 | Glutathione metabolism | 0.002069 | 0.049871 | Over-represented in disease |
| Franzosa et al. (UC) | Pathway-level DA | map00730 | Thiamine metabolism | 0.002126 | 0.049871 | Under-represented in disease |
| Franzosa et al. (UC) | Pathway-level DA | map00650 | Butanoate metabolism | 0.002634 | 0.060167 | Over-represented in disease |
| Franzosa et al. (UC) | Pathway-level DA | map00592 | alpha-Linolenic acid metabolism | 0.002773 | 0.061707 | Over-represented in disease |
| Franzosa et al. (UC) | Pathway-level DA | map00680 | Methane metabolism | 0.003179 | 0.064171 | Over-represented in disease |
| Franzosa et al. (UC) | Pathway-level DA | map00020 | Citrate cycle (TCA cycle) | 0.003327 | 0.065627 | Over-represented in disease |
| Franzosa et al. (UC) | Pathway-level DA | map00030 | Pentose phosphate pathway | 0.004094 | 0.071889 | Over-represented in disease |
| Franzosa et al. (UC) | Pathway-level DA | map00053 | Ascorbate and aldarate metabolism | 0.004141 | 0.071889 | Over-represented in disease |
| Franzosa et al. (UC) | Pathway-level DA | map00332 | Carbapenem biosynthesis | 0.004228 | 0.071961 | Under-represented in disease |
| Franzosa et al. (UC) | Pathway-level DA | map00220 | Arginine biosynthesis | 0.004753 | 0.075299 | Under-represented in disease |
| Franzosa et al. (UC) | Pathway-level DA | map01053 | Biosynthesis of siderophore group nonribosomal peptides | 0.004771 | 0.075299 | Over-represented in disease |
| Franzosa et al. (UC) | Pathway-level DA | map00401 | Novobiocin biosynthesis | 0.006934 | 0.096367 | Under-represented in disease |
| Franzosa et al. (UC) | Pathway-level DA | map00770 | Pantothenate and CoA biosynthesis | 0.006869 | 0.096367 | Under-represented in disease |
| Jeffery et al. (IBS) | Pathway-level DA | map00470 | D-Amino acid metabolism | 8.13E-05 | 0.002804 | Under-represented in disease |
| Jeffery et al. (IBS) | Pathway-level DA | map00550 | Peptidoglycan biosynthesis | 1.39E-04 | 0.002804 | Under-represented in disease |

| | | | | | | |
|---|---|---|---|---|---|---|
| Jeffery et al. (IBS) | Pathway-level DA | map00900 | Terpenoid backbone biosynthesis | 5.14E-05 | 0.002804 | Under-represented in disease |
| Jeffery et al. (IBS) | Pathway-level DA | map00908 | Zeatin biosynthesis | 1.17E-04 | 0.002804 | Under-represented in disease |
| Jeffery et al. (IBS) | Pathway-level DA | map00970 | Aminoacyl-tRNA biosynthesis | 9.24E-05 | 0.002804 | Under-represented in disease |
| Jeffery et al. (IBS) | Pathway-level DA | map00983 | Drug metabolism - other enzymes | 9.48E-05 | 0.002804 | Under-represented in disease |
| Jeffery et al. (IBS) | Pathway-level DA | map00561 | Glycerolipid metabolism | 1.76E-04 | 0.003047 | Over-represented in disease |
| Jeffery et al. (IBS) | Pathway-level DA | map00010 | Glycolysis / Gluconeogenesis | 2.27E-04 | 0.00343 | Under-represented in disease |
| Jeffery et al. (IBS) | Pathway-level DA | map00240 | Pyrimidine metabolism | 2.61E-04 | 0.003507 | Under-represented in disease |
| Jeffery et al. (IBS) | Pathway-level DA | map00620 | Pyruvate metabolism | 6.19E-04 | 0.007487 | Over-represented in disease |
| Jeffery et al. (IBS) | Pathway-level DA | map00052 | Galactose metabolism | 7.26E-04 | 0.007987 | Over-represented in disease |
| Jeffery et al. (IBS) | Pathway-level DA | map00640 | Propanoate metabolism | 7.92E-04 | 0.007987 | Over-represented in disease |
| Jeffery et al. (IBS) | Pathway-level DA | map00254 | Aflatoxin biosynthesis | 9.56E-04 | 0.008896 | Over-represented in disease |
| Jeffery et al. (IBS) | Pathway-level DA | map00520 | Amino sugar and nucleotide sugar metabolism | 0.001148 | 0.009275 | Over-represented in disease |
| Jeffery et al. (IBS) | Pathway-level DA | map00650 | Butanoate metabolism | 0.00115 | 0.009275 | Over-represented in disease |
| Jeffery et al. (IBS) | Pathway-level DA | map00910 | Nitrogen metabolism | 0.002467 | 0.018653 | Over-represented in disease |
| Jeffery et al. (IBS) | Pathway-level DA | map00051 | Fructose and mannose metabolism | 0.003 | 0.021354 | Over-represented in disease |
| Jeffery et al. (IBS) | Pathway-level DA | map00140 | Steroid hormone biosynthesis | 0.004404 | 0.029147 | Over-represented in disease |
| Jeffery et al. (IBS) | Pathway-level DA | map00600 | Sphingolipid metabolism | 0.004577 | 0.029147 | Over-represented in disease |
| Jeffery et al. (IBS) | Pathway-level DA | map00627 | Aminobenzoate degradation | 0.005181 | 0.031343 | Over-represented in disease |
| Jeffery et al. (IBS) | Pathway-level DA | map00630 | Glyoxylate and dicarboxylate metabolism | 0.007873 | 0.045366 | Over-represented in disease |
| Jeffery et al. (IBS) | Pathway-level DA | map00770 | Pantothenate and CoA biosynthesis | 0.008449 | 0.046469 | Under-represented in disease |
| Jeffery et al. (IBS) | Pathway-level DA | map00332 | Carbapenem biosynthesis | 0.009367 | 0.049279 | Under-represented in disease |
| Jeffery et al. (IBS) | Pathway-level DA | map00071 | Fatty acid degradation | 0.01074 | 0.052993 | Over-represented in disease |
| Jeffery et al. (IBS) | Pathway-level DA | map00670 | One carbon pool by folate | 0.010949 | 0.052993 | Under-represented in disease |
| Jeffery et al. (IBS) | Pathway-level DA | map00710 | Carbon fixation in photosynthetic organisms | 0.013107 | 0.061 | Under-represented in disease |
| Jeffery et al. (IBS) | Pathway-level DA | map00280 | Valine, leucine and isoleucine degradation | 0.014809 | 0.063997 | Over-represented in disease |
| Jeffery et al. (IBS) | Pathway-level DA | map00511 | Other glycan degradation | 0.014519 | 0.063997 | Over-represented in disease |
| Jeffery et al. (IBS) | Pathway-level DA | map00780 | Biotin metabolism | 0.016594 | 0.069238 | Over-represented in disease |
| Jeffery et al. (IBS) | Pathway-level DA | map00531 | Glycosaminoglycan degradation | 0.018977 | 0.075413 | Over-represented in disease |
| Jeffery et al. (IBS) | Pathway-level DA | map00760 | Nicotinate and nicotinamide metabolism | 0.019321 | 0.075413 | Under-represented in disease |
| Jeffery et al. (IBS) | Pathway-level DA | map00030 | Pentose phosphate pathway | 0.022655 | 0.085346 | Over-represented in disease |
| Jeffery et al. (IBS) | Pathway-level DA | map00053 | Ascorbate and aldarate metabolism | 0.024009 | 0.085346 | Over-represented in disease |
| Jeffery et al. (IBS) | Pathway-level DA | map00720 | Carbon fixation pathways in prokaryotes | 0.023691 | 0.085346 | Under-represented in disease |
| Jeffery et al. (IBS) | Pathway-level DA | map01053 | Biosynthesis of siderophore group nonribosomal peptides | 0.024687 | 0.085346 | Over-represented in disease |
| Jeffery et al. (IBS) | Pathway-level DA | map00380 | Tryptophan metabolism | 0.026153 | 0.087903 | Over-represented in disease |
| Jeffery et al. (IBS) | Pathway-level DA | map00195 | Photosynthesis | 0.027123 | 0.088699 | Under-represented in disease |
| Jeffery et al. (IBS) | Pathway-level DA | map00230 | Purine metabolism | 0.028699 | 0.08998 | Under-represented in disease |
| Jeffery et al. (IBS) | Pathway-level DA | map00310 | Lysine degradation | 0.029002 | 0.08998 | Over-represented in disease |
| Jeffery et al. (IBS) | Pathway-level DA | map00190 | Oxidative phosphorylation | 0.029755 | 0.090008 | Under-represented in disease |
| Jeffery et al. (IBS) | Pathway-level DA | map00604 | Glycosphingolipid biosynthesis - ganglio series | 0.030845 | 0.091031 | Over-represented in disease |
| Wang et al. (ESRD) | ORA-ECs | map00730 | Thiamine metabolism | 4.08E-04 | 0.022413 | Under-represented in disease |
| Franzosa et al. (CD) | ORA-ECs | map00230 | Purine metabolism | 0.001781 | 0.099747 | Over-represented in disease |
| Franzosa et al. (CD) | ORA-ECs | map00220 | Arginine biosynthesis | 8.11E-05 | 0.00454 | Under-represented in disease |
| Franzosa et al. (CD) | ORA-ECs | map00340 | Histidine metabolism | 1.97E-04 | 0.005519 | Under-represented in disease |
| Franzosa et al. (UC) | ORA-ECs | map00540 | Lipopolysaccharide biosynthesis | 5.62E-06 | 3.15E-04 | Over-represented in disease |
| Franzosa et al. (UC) | ORA-ECs | map00785 | Lipoic acid metabolism | 0.001267 | 0.035478 | Over-represented in disease |
| Franzosa et al. (UC) | ORA-ECs | map00480 | Glutathione metabolism | 0.002465 | 0.046014 | Over-represented in disease |
| Franzosa et al. (UC) | ORA-ECs | map00400 | Phenylalanine, tyrosine and tryptophan biosynthesis | 4.34E-04 | 0.024307 | Under-represented in disease |
| Franzosa et al. (UC) | ORA-ECs | map00340 | Histidine metabolism | 0.001357 | 0.03799 | Under-represented in disease |
| Franzosa et al. (UC) | ORA-ECs | map00540 | Lipopolysaccharide biosynthesis | 1.84E-04 | 0.010308 | Associated with disease |
| Franzosa et al. (CD) | ORA-Metabolites | map04978 | Mineral absorption | 2.27E-04 | 0.014999 | Over-represented in disease |
| Franzosa et al. (CD) | ORA-Metabolites | map00120 | Primary bile acid biosynthesis | 0.0012 | 0.039608 | Over-represented in disease |
| Franzosa et al. (CD) | ORA-Metabolites | map04974 | Protein digestion and absorption | 0.002147 | 0.047235 | Over-represented in disease |
| Jeffery et al. (IBS) | ORA-Metabolites | map04974 | Protein digestion and absorption | 1.24E-05 | 7.09E-04 | Over-represented in disease |
| Jeffery et al. (IBS) | ORA-Metabolites | map00970 | Aminoacyl-tRNA biosynthesis | 1.12E-04 | 0.003182 | Over-represented in disease |
| Jeffery et al. (IBS) | ORA-Metabolites | map00270 | Cysteine and methionine metabolism | 1.98E-04 | 0.003761 | Over-represented in disease |
| Jeffery et al. (IBS) | ORA-Metabolites | map04978 | Mineral absorption | 3.43E-04 | 0.004885 | Over-represented in disease |
| Jeffery et al. (IBS) | ORA-Metabolites | map00997 | Biosynthesis of various other secondary metabolites | 0.002531 | 0.026665 | Over-represented in disease |
| Jeffery et al. (IBS) | ORA-Metabolites | map00470 | D-Amino acid metabolism | 0.003275 | 0.026665 | Over-represented in disease |
| Jeffery et al. (IBS) | ORA-Metabolites | map05230 | Central carbon metabolism in cancer | 0.003275 | 0.026665 | Over-represented in disease |
| Jeffery et al. (IBS) | ORA-Metabolites | map01230 | Biosynthesis of amino acids | 0.005957 | 0.042447 | Over-represented in disease |
| Jeffery et al. (IBS) | ORA-Metabolites | map00260 | Glycine, serine and threonine metabolism | 0.015949 | 0.095025 | Over-represented in disease |
| Jeffery et al. (IBS) | ORA-Metabolites | map01210 | 2-Oxocarboxylic acid metabolism | 0.016671 | 0.095025 | Over-represented in disease |
| Jeffery et al. (IBS) | ORA-Metabolites | map04974 | Protein digestion and absorption | 7.70E-04 | 0.043874 | Associated with disease |
| Jeffery et al. (IBS) | ORA-Metabolites | map00970 | Aminoacyl-tRNA biosynthesis | 0.002521 | 0.071859 | Associated with disease |
